# Bayesian Parameter Balancing Enables Robust and Consistent Estimation of Kinetic Parameter Uncertainty

**DOI:** 10.64898/2026.05.18.726043

**Authors:** TrungTin Nguyen, Binh H. Ho, Michael Pan, Jennifer A. Flegg, Michael J. McDonald, Christopher Drovandi

## Abstract

**Motivation:** Kinetic models are central to systems biology, but enzyme-kinetic parameters compiled from the literature and databases are often incomplete, inconsistent, and measured under heterogeneous conditions. Classical parameter balancing helps infer missing parameters, yet it often lacks calibrated uncertainty, robustness to misspecification, and explicit treatment of source-level heterogeneity.

**Results:** We develop a formal Bayesian parameter balancing framework that enforces thermodynamic constraints, estimates full posterior uncertainty, and validates calibration using leave-one-out cross-validation and posterior-predictive coverage. Beyond the classical Gaussian formulation, we introduce robust Student-*t* and skewed error models to improve reliability under outliers and model misspecification, and incorporate random effects to account for source-level or group-level variability across studies. The resulting approach yields thermodynamically consistent parameter sets with well-calibrated credible intervals on held-out data, offering a Bayesian parameter balancing approach useful to systems biology researchers.

**Availability and implementation:** Source code, data, workflows, a Julia package and command-line usage are available at [the project GitHub repository](https://github.com/Trung-TinNGUYEN/Robust_Bayesian_Parameter_Balancing).

**Graphical Abstract:** 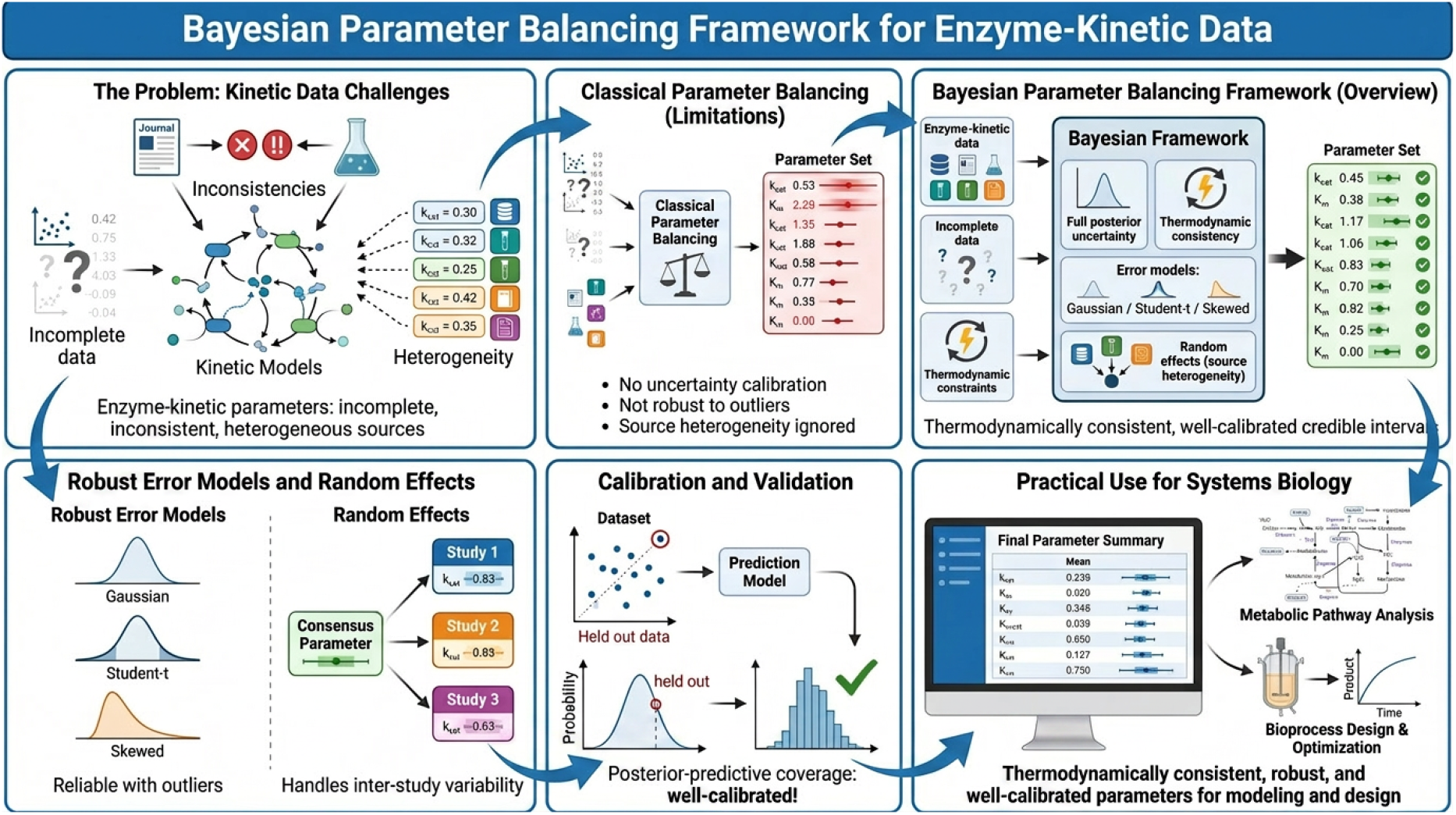

## 1 Introduction

Mathematical models of cellular biochemistry require many kinetic parameters, including Michaelis constants, turnover rates, and elementary kinetic constants. Although such quantities are available from literature and curated resources such as BRENDA, SABIO-RK, and eQuilibrator [1–5], the resulting data are often sparse, noisy, heterogeneous, and mutually inconsistent. Values measured under different assay conditions may not be directly comparable, and naive aggregation can violate thermodynamic consistency. This issue becomes especially important in large reconstructions such as Yeast9 [6, 7] and other genome-scale models [8, 9], where local inconsistencies can compromise downstream simulation.

Systems biology has developed several ways to enforce physical consistency, including thermodynamic and detailed-balance constraints [10, 11], modular thermodynamic-kinetic formulations [12–14], and bond-graph methods [15–19]. Parameter balancing is a practical member of this family: it defines independent basis parameters and derives all remaining quantities through algebraic and thermodynamic relationships, so equilibrium constraints such as Haldane relationships are satisfied by construction [20, 21]. It has been used to parameterize pathway, genome-scale, and whole-cell models [22–25].

Classical parameter balancing is useful, but its statistical assumptions are often implicit. Likelihoods, priors on transformed parameters, and observation-noise models are not always stated as a complete probabilistic model, and reported intervals are rarely checked for calibration. Gaussian formulations are also fragile when curated biochemical data contain outliers, heavy tails, skewness, or systematic laboratory and source effects. These limitations motivate a more explicit Bayesian treatment with robust likelihoods and hierarchical structure.

Here we formulate parameter balancing as Bayesian inference for log-basis parameters ***q*** through a constrained linear observation map ***Q***_obs_***q***. The resulting posterior gives both balanced parameter estimates and uncertainty, and the Gaussian case recovers classical parameter balancing as a conjugate baseline. We then validate uncertainty using leave-one-record-out and leave-one-source-out posterior prediction, so nominal intervals are assessed on held-out data rather than only by in-sample variance formulas.

A central goal of this work is to move beyond point estimation and rough uncertainty summaries toward calibrated uncertainty quantification. These diagnostics allow us to check whether credible intervals behave as advertised on held-out data, rather than relying only on in-sample fit or posterior variance formulas. We show that when Gaussian error assumptions are violated, classical parameter balancing can become substantially overconfident.

We therefore extend the Gaussian baseline in two directions. First, we introduce robust likelihoods based on Student-*t* and skew-normal error models for heavy-tailed and asymmetric measurement errors. Second, we incorporate random effects to model source- or group-level variation, allowing the method to better reconcile measurements collected from different laboratories, databases, or assay settings. The resulting framework retains the biochemical consistency enforced by parameter balancing while delivering more reliable uncertainty.

### Main contributions

We connect parameter balancing to robust and calibrated Bayesian inference [26–28]. We provide an explicit Bayesian formulation of classical balancing; develop Student-*t*, skew-normal, and source random-effects extensions; evaluate point accuracy separately from interval calibration; and study computational scaling, including thread-level parallelism and an exact-evidence lower bound (ELBO) structured variational approximation. The results show that Gaussian balancing can remain competitive in point error while failing in uncertainty calibration, whereas robust and hierarchical alternatives yield more reliable predictive intervals.

### Paper organisation

Section 2 presents the Bayesian parameter-balancing framework, including the thermodynamic dependency structure, robust likelihoods, and hierarchical source effects. Section 3 reports simulation studies and held-out pathway analyses for glycolysis and the tricarboxylic acid (TCA) cycle. Additional notation is provided in Supplementary Section B, kinetic background in Supplementary Section C, robust and hierarchical derivations in Supplementary Section D, evaluation definitions in Supplementary Section E, and computational, reproducibility, and extended empirical results in Supplementary Sections F, F.2 and F.2.3.

### Notation and abbreviations

Table 1 summarizes the main notation; additional notation for the robust likelihoods, hierarchical source effects, and fold-wise predictive diagnostics is provided in Supplementary Tables S1 and S2.

**Table 1:**
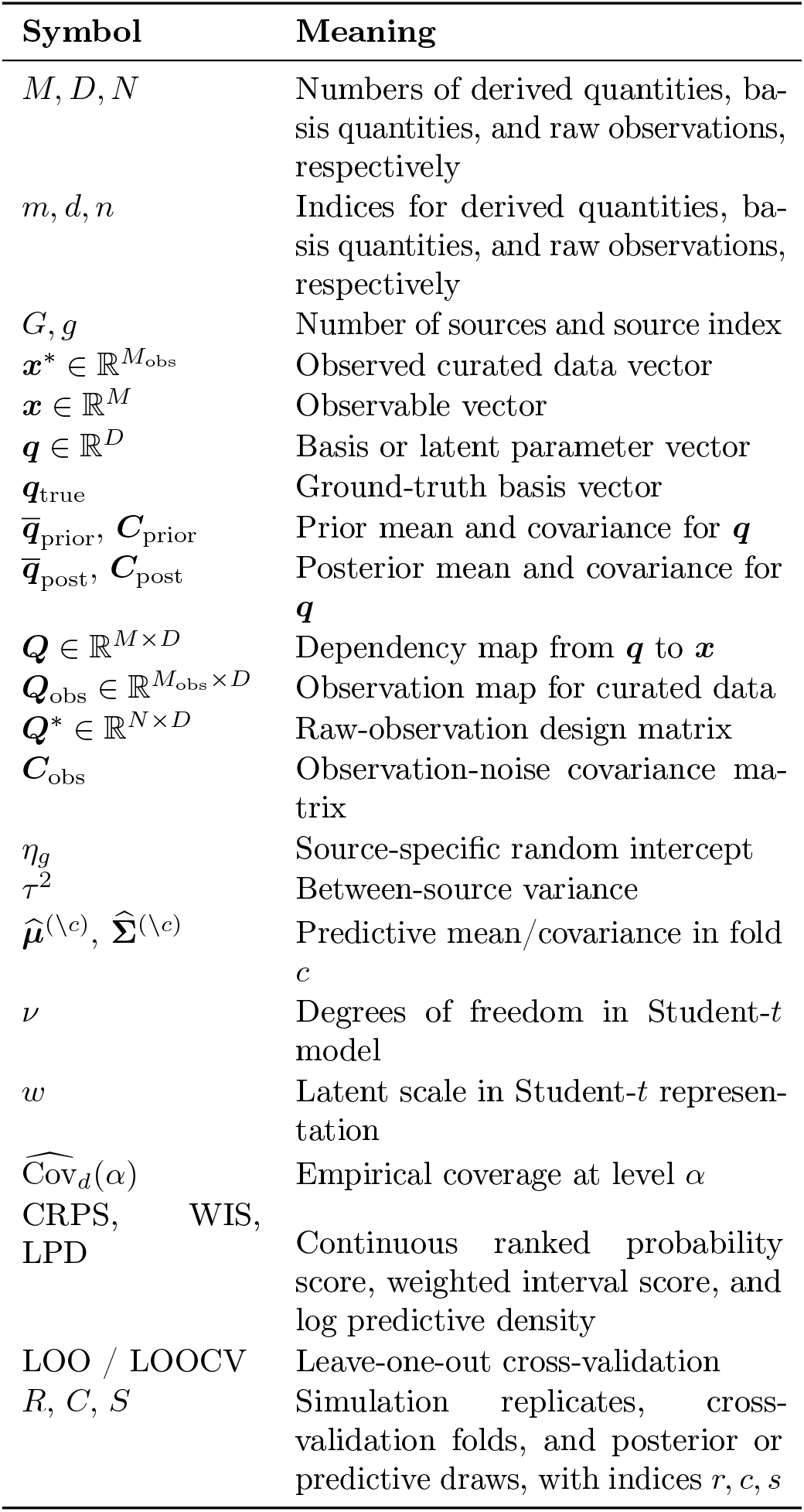
Key notation and abbreviations used in the main text.

| Symbol | Meaning |
| --- | --- |
| $M, D, N$ | Numbers of derived quantities, basis quantities, and raw observations, respectively |
| $m, d, n$ | Indices for derived quantities, basis quantities, and raw observations, respectively |
| $G, g$ | Number of sources and source index |
| $\mathbf{x}^* \in \mathbb{R}^{M_{\text{obs}}}$ | Observed curated data vector |
| $\mathbf{x} \in \mathbb{R}^M$ | Observable vector |
| $\mathbf{q} \in \mathbb{R}^D$ | Basis or latent parameter vector |
| $\mathbf{q}_{\text{true}}$ | Ground-truth basis vector |
| $\bar{\mathbf{q}}_{\text{prior}}, \mathbf{C}_{\text{prior}}$ | Prior mean and covariance for $\mathbf{q}$ |
| $\bar{\mathbf{q}}_{\text{post}}, \mathbf{C}_{\text{post}}$ | Posterior mean and covariance for $\mathbf{q}$ |
| $\mathbf{Q} \in \mathbb{R}^{M \times D}$ | Dependency map from $\mathbf{q}$ to $\mathbf{x}$ |
| $\mathbf{Q}_{\text{obs}} \in \mathbb{R}^{M_{\text{obs}} \times D}$ | Observation map for curated data |
| $\mathbf{Q}^* \in \mathbb{R}^{N \times D}$ | Raw-observation design matrix |
| $\mathbf{C}_{\text{obs}}$ | Observation-noise covariance matrix |
| $\eta_g$ | Source-specific random intercept |
| $\tau^2$ | Between-source variance |
| $\hat{\mu}^{(\setminus c)}, \hat{\Sigma}^{(\setminus c)}$ | Predictive mean/covariance in fold $c$ |
| $\nu$ | Degrees of freedom in Student-$t$ model |
| $w$ | Latent scale in Student-$t$ representation |
| $\widehat{\text{Cov}}_d(\alpha)$ | Empirical coverage at level $\alpha$ |
| CRPS, WIS, LPD | Continuous ranked probability score, weighted interval score, and log predictive density |
| LOO / LOOCV | Leave-one-out cross-validation |
| $R, C, S$ | Simulation replicates, cross-validation folds, and posterior or predictive draws, with indices $r, c, s$ |

## 2 Materials and methods

### 2.1 Pathway models and curated data

We study two benchmark pathways, glycolysis and the TCA cycle, because both are well characterized and contain sufficient thermodynamic and kinetic information for controlled parameter balancing experiments. For each pathway, we constructed a thermodynamically consistent reference parameter set by combining curated literature values from pathway-specific kinetic datasets with equilibrium information from eQuilibrator [3–5]. The resulting reference values, shown in Figures 1 and 2 (blue bars), serve as the baseline parameter sets for downstream simulation and validation.

**Figure 1:**
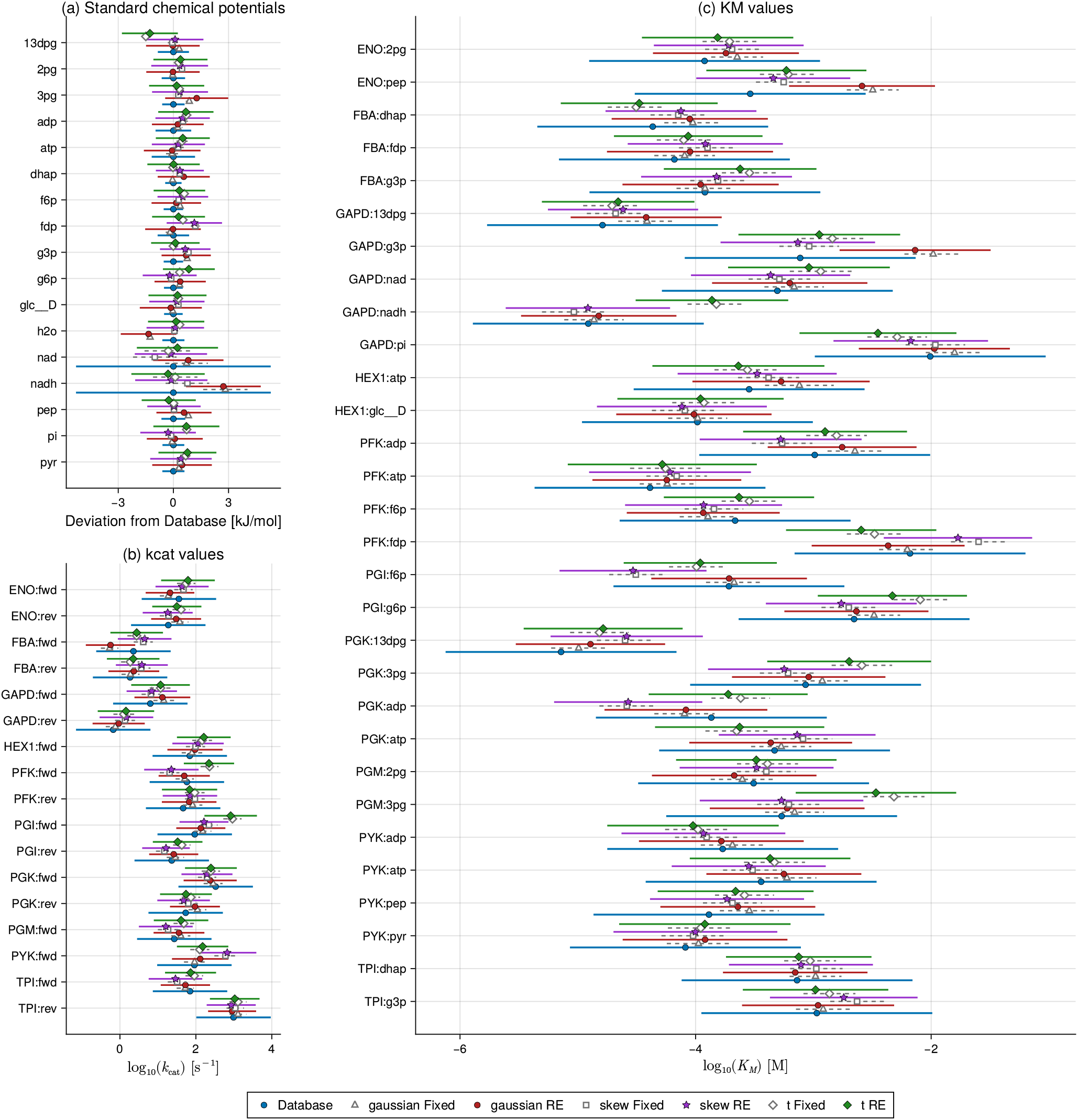
Synthetic glycolysis: all fitted models versus the reference observable profile. Panel (a) shows standard chemical potentials Δ*μ°′*, panel (b) shows log_10_(*k*_cat_), and panel (c) shows log_10_(*K*_*M*_). Here, Fix denotes the fixed-effects specification and RE denotes the random-effects specification. Across all three observable blocks, the six fitted models recover the reference structure closely. The dominant between-model differences appear in interval width rather than in posterior means, indicating that the model families largely agree on the central biochemical profile but differ in how uncertainty is distributed across weakly identified kinetic quantities. In particular, the hierarchical RE fits widen uncertainty selectively rather than uniformly, which is consistent with source-level heterogeneity affecting only part of the pathway.

**Figure 2:**
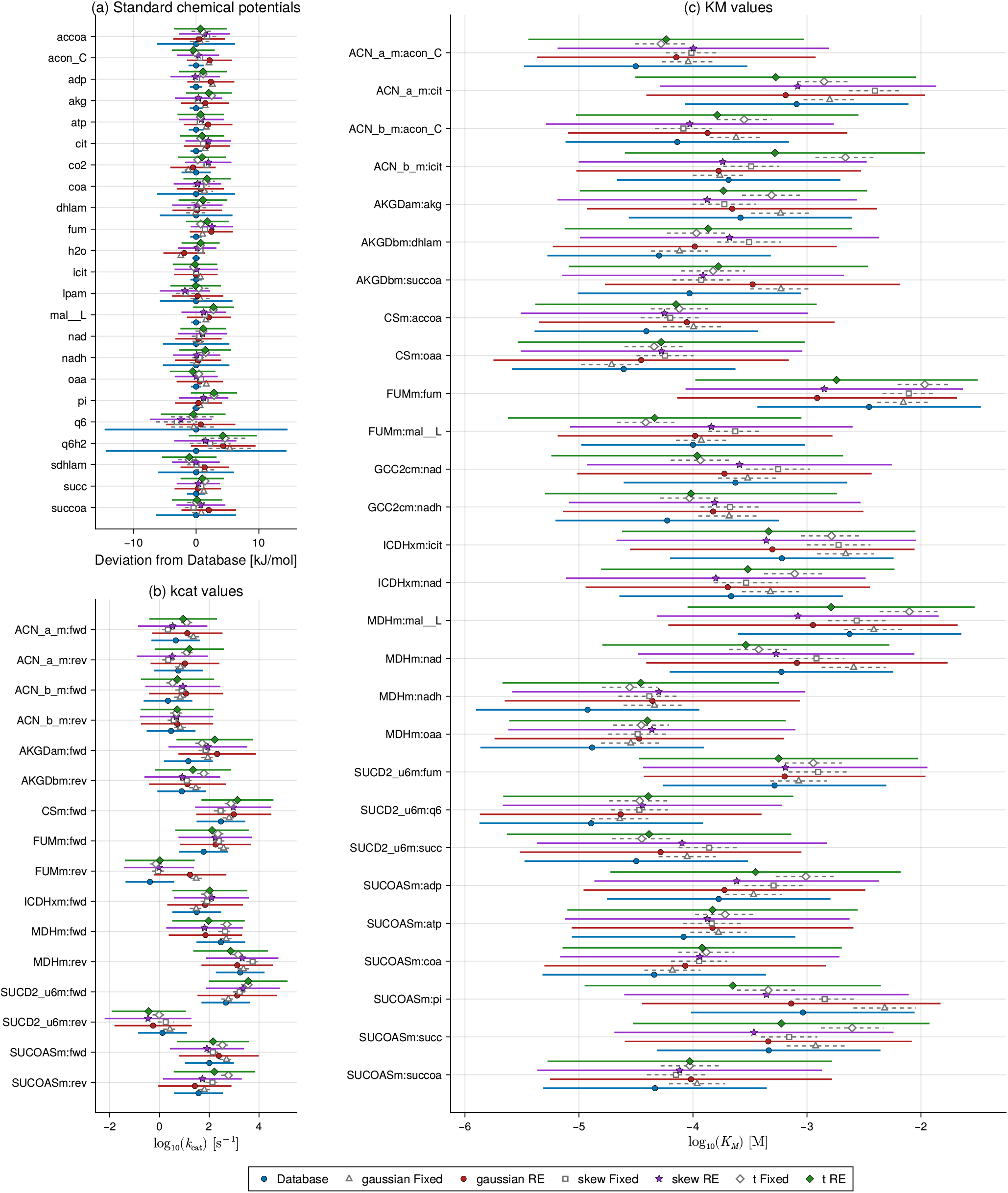
Synthetic TCA cycle: all fitted models versus the reference observable profile. Panel (a) shows standard chemical potentials Δ*μ°′*, panel (b) shows log_10_(*k*_cat_), and panel (c) shows log_10_(*K*_*M*_). As in glycolysis, all six specifications recover the reference observable structure well, with the largest between-model differences concentrated in interval width rather than in posterior means. The pattern is especially clear for selected kinetic quantities where random-effects fits redistribute uncertainty more broadly than their fixed-effects counterparts. Under heterogeneous synthetic data, the hierarchical extension therefore preserves thermodynamic consistency while quantifying source-level variability more honestly.

The kinetic context is a metabolic reaction network with *N*_met_ metabolites and *N*_rxn_ reactions. Let 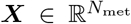 denote the metabolite concentration vector, 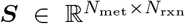 the stoichiometric matrix, and 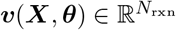 the reaction-rate vector, where ***θ*** ∈ ℝ^*P*^ collects the kinetic parameters governing the rates. The corresponding dynamics can be written as

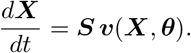

In typical biochemical models, the entries of ***θ*** include catalytic rate constants, Michaelis constants, enzyme abundances, metabolite concentrations, and thermodynamic quantities such as standard chemical potentials or equilibrium constants. Although our inference does not fit the ordinary differential equation dynamics directly, this mechanistic setting determines which parameters are linked and which physical constraints must be respected.

Following the classical parameter balancing literature [13, 20, 21], we work on a transformed scale on which the relevant parameter dependencies become linear. We denote the full collection of transformed model quantities by ***x*** ∈ ℝ^*M*^, and a lower-dimensional set of independent basis quantities by ***q*** ∈ ℝ^*D*^, *D* ≤ *M*. These are linked through the dependency map

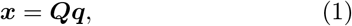

where ***Q*** ∈ ℝ^*M ×D*^ is determined by the pathway structure and the chosen thermodynamic parameterization. Observed quantities are extracted through the corresponding observation map 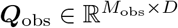, yielding the curated data vector 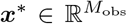. In practice, ***Q***_obs_ is obtained from ***Q*** by selecting, deleting, and, when needed, duplicating rows to align the dependency structure with the available observations. The associated observation-noise covariance is 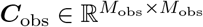. This construction ensures that inferred parameter sets remain thermodynamically consistent by design.

### 2.2 Metabolic parameter dependencies and thermodynamic consistency

The role of ***Q*** is to encode biochemical relationships among kinetic and thermodynamic quantities. Forward and reverse catalytic constants, Michaelis constants, and equilibrium constants cannot in general be chosen independently; they must satisfy thermodynamic identities such as Haldane relationships, which link kinetics to equilibrium thermodynamics. As a result, naive assembly of literature values can produce parameter sets that are numerically plausible in isolation but physically incompatible when combined.

Parameter balancing resolves this by reparameterizing the model in terms of independent basis quantities ***q***, from which all derived quantities ***x*** are obtained through ***Q***. In practice, ***q*** includes transformed rate, affinity, binding, and thermodynamic terms, while ***x*** contains the full set of kinetic and thermodynamic quantities of interest. After transformation, these dependencies are linear, which makes it possible to formulate parameter balancing as a constrained statistical inference problem while preserving biochemical plausibility. Additional kinetic background, including an illustrative reversible reaction, convenience kinetics, and the induced Haldane constraint, is provided in Supplementary Sections C.2 to C.4.

### 2.3 Formal Bayesian parameter balancing

We formulate parameter balancing as Bayesian inference on the basis parameters ***q***. In classical parameter balancing, one first assembles a curated data vector ***x***\* containing all available numerical information about the model quantities, including literature values for *K*_*M*_, *k*_cat_, metabolite concentrations, and equilibrium-related quantities. Some entries of ***x***\* may be missing, while others may summarize multiple measurements with associated uncertainty. The aim is to infer a complete, self-consistent parameter vector ***x*** that matches the observed entries as closely as possible while respecting the structural constraints encoded by ***Q***.

The forward model is defined as in Equation (1) and the observed quantities are linked to the basis parameters through the observation map ***Q***_obs_. The Gaussian baseline model is

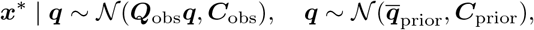

where ***C***_obs_ encodes observation uncertainty and 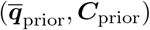 specify prior biochemical plausibility on the transformed scale. Equivalently, the observation model can be written in regression form as

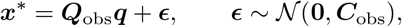

so that parameter balancing is a Bayesian linear regression problem in the latent basis vector ***q***, with design matrix ***Q***_obs_, response vector ***x***\*, and Gaussian prior on the regression coefficients. This makes explicit the probabilistic structure that is only implicit in classical parameter balancing.

Under this linear-Gaussian model, the posterior is again Gaussian and is available in closed form:

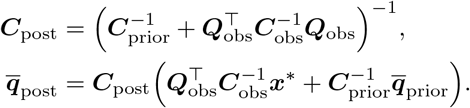

The posterior mean 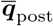 gives the balanced basis parameter estimates, while ***C***_post_ quantifies their uncertainty. Posterior summaries for the full parameter vector follow by propagation through ***Q***, giving

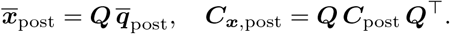

Thus, the balanced parameter set is obtained together with uncertainty for both basis and derived quantities.

Parameters without direct observations remain prior-informed but constrained through ***Q***, while conflicting measurements are reconciled according to their uncertainties. The posterior predictive distribution also provides a basis for held-out calibration checks, consistent with the broader Bayesian workflow perspective on model checking and evaluation [29, 30].

### 2.4 Robust likelihoods and hierarchical source effects

To address misspecification, outliers, and source heterogeneity, we extended the Gaussian baseline with Student-*t* errors for heavy tails, skew-normal errors for asymmetric noise, and source-level random effects for between-source variation. The robust likelihoods use conditional Gaussian representations, while the hierarchical term preserves the thermodynamic structure encoded by ***Q***.

For the raw-data analyses, let 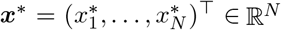 denote the log-transformed records, and let ***Q***\* ∈ ℝ^*N×D*^ be the raw-observation design matrix whose *n*-th row maps record *n* to the corresponding derived quantity. We write

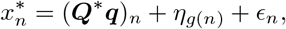

where *g*(*n*) denotes the publication or curated dataset from which record *n* was extracted. The source-specific random intercept *η*_*g*(*n*)_ captures broad publication- or dataset-level shifts without introducing a separate random effect for every source–parameter combination. We assume

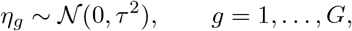

independently across sources. The Student-*t* model uses a degrees-of-freedom parameter *v >* 2, while the skew-normal model uses shape parameter *α*. In raw BRENDA analyses, *v* was fixed at a weakly heavy-tailed value and *α* was profiled within each training fold; in synthetic experiments, the robust fits used the corresponding data-generating or sweep values described in Supplementary Sections D.3 and F.1. The thermodynamic block is included in every fold through the same Gaussian contribution used in the baseline model.

### 2.5 Fold-wise posterior predictive evaluation

We evaluated predictive calibration on raw BRENDA records using leakage-free fold-wise cross-validation. Let *c*(*n*) denote the fold key for record *n*, which may correspond to the record index, the source label *g*(*n*), the Enzyme Commission classification of the associated enzyme, or the derived-quantity index, depending on the experiment. The mapping *n* ↦ *c*(*n*) partitions the observation indices into disjoint folds, and each fold is evaluated by holding out one partition at a time and fitting the model on its complement. For fold *c*, define the held-out and training sets by

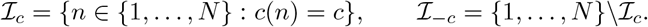

All fitting, hyperparameter learning, and robustness-parameter handling used only the training records indexed by ℐ_*−*c_, together with the always-included thermodynamic block.

For each held-out record *n* ∈ ℐ_c_, let *g*\* = *g*(*n*) denote its source. If *g*\* appears in the training fold, prediction conditions on the sampled source effect 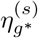. If *g*\* is absent from training, as in leave-one-source-out cross-validation, we instead draw

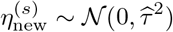

independently across posterior draws, where 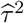 is the fitted between-source variance. This is the appropriate hierarchical predictive rule for unseen sources.

Given a posterior draw ***q***^(*s*)^ and the corresponding source effect, the predictive mean for held-out record *n* is

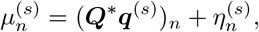

where 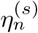 denotes either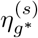 or 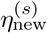, depending on whether the source is represented in training. We then generated predictive draws from the same observation family used in the fitted model, using the fitted robustness parameters 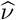 or 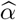 where applicable, together with the record-specific variance 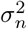. For the Gaussian model with an unseen source, the source effect can be integrated out analytically, yielding

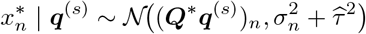

in the non-contaminated case. For Student-*t* and skew-normal models, we estimated the predictive law by Monte Carlo over the source effect and the likelihood-specific latent variables.

From the *S* posterior predictive draws 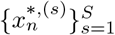, we compute central predictive intervals, probability integral transform (PIT) values, continuous ranked probability score (CRPS), weighted interval score (WIS), and log predictive density (LPD); see Supplementary Sections E.4 to E.6. For nominal level *p*, if [*L*_*n,p*_, *U*_*n,p*_] is the central 100*p*% predictive interval, then

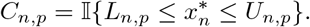

Empirical coverage averages *C*_n,p_ over held-out records.

### 2.6 Simulation design

We evaluated inference under controlled misspecification by generating synthetic datasets from the reference pathway parameter sets. Let ***q***_true_ denote the reference basis vector. Synthetic observations were generated from

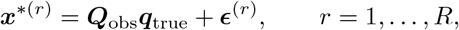

under three data-generating families: Gaussian, Student-*t*, and skew-normal noise. This allowed us to compare well-specified settings with heavy-tailed and asymmetric misspecification. For each simulated dataset, we fitted the Gaussian baseline and, where appropriate, the corresponding robust alternative. The main synthetic recovery experiments use the variance-matched robust families defined in Supplementary Section D.3. The controlled misspecification sweeps are deliberately more diagnostic: the degrees-of-freedom sweep varies the canonical Student-*t* tail parameter while matching the robust comparator to the corresponding sweep variance, and the skewness sweep uses the location-parameterized skew-normal family with the robust comparator fixed at the corresponding sweep value.

### 2.7 Evaluation protocol

We evaluated both parameter recovery and predictive calibration. For synthetic experiments, we summarized point estimation by bias and root mean squared error, and uncertainty calibration by empirical coverage of nominal credible intervals across simulation replicates. For held-out prediction, we used leave-one-out and leave-one-source-out cross-validation with posterior-predictive intervals to assess whether uncertainty was well calibrated on unseen observations. In the main text, we focus on the primary calibration and predictive findings; full interval definitions are provided in Supplementary Sections E.2 and E.4, predictive scoring rules in Supplementary Section E.5, additional coverage summaries in Supplementary Section F.1, and computational scaling results in Supplementary Section F.2.

For the Gaussian baseline, posterior means and covariances were computed analytically. Robust and hierarchical fits used conditional Gaussian representations with Gibbs-type MCMC updates for latent variables and source effects, embedded in the fold-wise MCEM procedure [31, 32]; see Supplementary Sections D.3, D.4, D.6 and D.7.

## 3 Results

We evaluated classical and robust Bayesian parameter balancing on glycolysis and the TCA cycle using synthetic experiments and held-out prediction. Across settings, the Gaussian baseline often remained competitive in point accuracy, but robust likelihoods and hierarchical source effects improved uncertainty calibration.

We begin with a controlled synthetic experiment designed to assess how the fitted models recover a chemically coherent reference profile in observable space. For each pathway, the reference quantities are the synthetic database-level observables implied by the balanced parameter set used to generate the data. Figures 1 and 2 compare these reference values with posterior summaries from the six fitted models: Gaussian, Student-*t*, and skew-normal likelihoods, each with and without random effects.

Two features are immediately apparent. First, across both glycolysis and the TCA cycle, all six specifications recover the main thermodynamic and kinetic structure of the reference profile. Posterior means remain close to the database values for most standard chemical potentials, catalytic constants, and Michaelis constants. This indicates that the robust and hierarchical extensions preserve the core biochemical reconstruction achieved by classical parameter balancing rather than replacing it with a qualitatively different fitted state.

Second, the main between-model differences appear in the uncertainty bands rather than in systematic shifts of the posterior means. In both pathways, the largest contrasts are concentrated in a subset of kinetic observables, especially within the *K*_*M*_ and *k*_cat_ blocks, where the random-effects fits redistribute uncertainty more flexibly than their fixed-effects counterparts. Thus, the synthetic forest plots should be read primarily as recovery diagnostics: they show that the model families agree on the broad observable structure, while differing in how uncertainty is allocated across weakly identified or heterogeneity-sensitive quantities.

This interpretation is important for the remainder of the Results section. The practical advantage of the robust and hierarchical formulations is not that they dramatically shift the central pathway estimates, but that they quantify uncertainty more faithfully when the data-generating process departs from the Gaussian fixed-effects idealization. That distinction is developed more directly in the coverage and held-out predictive analyses below.

### 3.1 Robust likelihoods improve interval calibration under misspecification

We first examined interval calibration under controlled synthetic misspecification. When data were generated under the Gaussian model, the classical Gaussian formulation achieved close-to-nominal coverage, as expected. However, when the data-generating mechanism was heavy-tailed or skewed, the Gaussian model systematically under-covered. This pattern was observed in both glycolysis and the TCA cycle: credible intervals from the classical model became too narrow as tails thickened or skew increased.

This pattern is shown clearly in Figure 3. For glycolysis, the Gaussian intervals remain close to the nominal 95% target when the data are nearly Gaussian, but coverage drops steadily as the tails thicken (i.e., as the degrees of freedom decrease) in panel (a) or as skew increases (*α* diverges from zero) in panel (b). The same qualitative behaviour appears for the TCA cycle in Figure 4. The decline is not subtle: under strong misspecification, the Gaussian baseline can fall well below the nominal level, indicating systematic overconfidence rather than mere Monte Carlo fluctuation.

**Figure 3:**
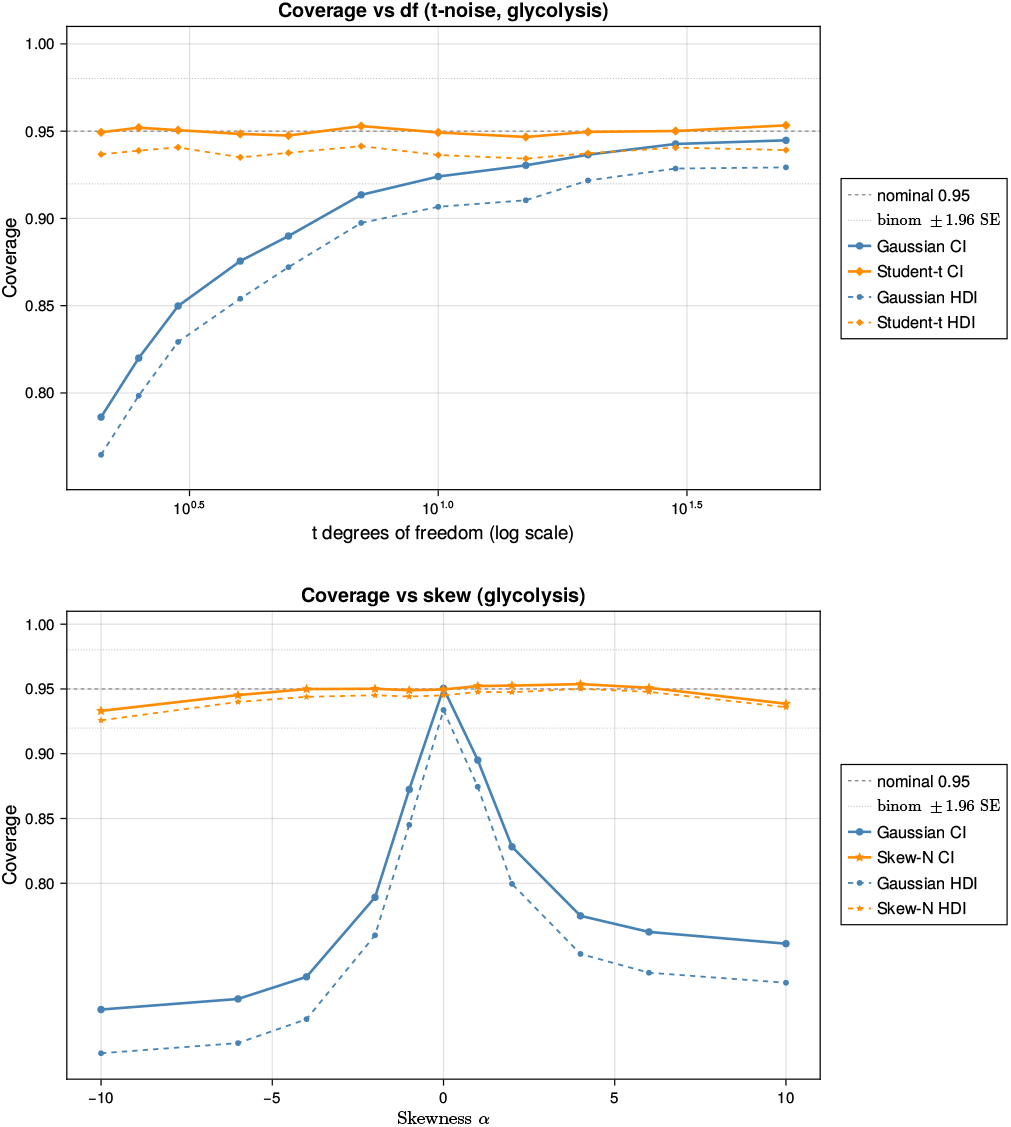
Glycolysis coverage under misspecification. The upper plot studies heavy-tailed misspecification under a Student-*t* data-generating mechanism, and the lower plot studies asymmetric misspecification under a skew-normal data-generating mechanism. In both cases, Gaussian intervals increasingly under-cover as misspecification becomes more pronounced, whereas the corresponding robust likelihoods remain much closer to the nominal 95% target. This same qualitative pattern later reappears in held-out prediction on raw BRENDA data, indicating that the synthetic misspecification study is informative about the real-data calibration behaviour.

**Figure 4:**
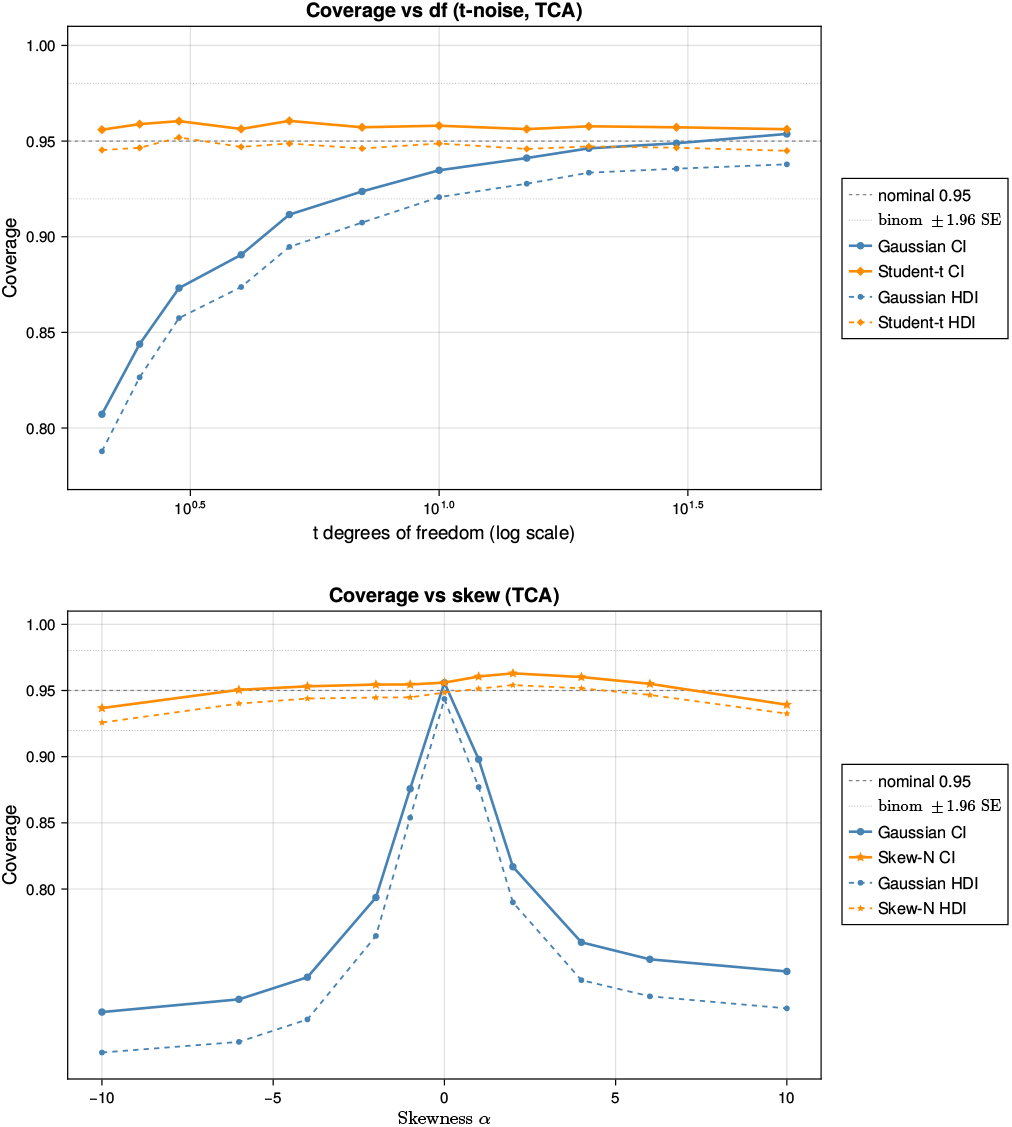
TCA-cycle coverage under misspecification. The top panel shows heavy-tailed misspecification and the bottom panel shows asymmetric misspecification. The same qualitative pattern as in Figure 3 is recovered: Gaussian intervals become increasingly overconfident as misspecification grows, whereas the robust likelihoods remain substantially better-calibrated across the range of tail-heaviness and skewness considered.

By contrast, robust likelihoods substantially improved calibration. Student-*t* likelihoods restored coverage under heavy-tailed noise, and skew-normal likelihoods improved performance under asymmetric noise. The main effect was not a dramatic change in point estimates, but a correction of posterior uncertainty so that the intervals behaved much more nearly as advertised.

### 3.2 Point accuracy remains stable while uncertainty quality changes

Across both pathways, bias and root mean squared error (RMSE) were broadly similar across Gaussian and robust fits in well-specified and mildly misspecified settings. Thus, classical and robust parameter balancing often produced comparable point estimates even when their uncertainty quantification differed sharply. This separation between point accuracy and interval calibration is important for interpretation. A method can remain competitive in RMSE while still providing misleadingly narrow intervals.

The complete numerical summaries for the 200-replicate misspecification sweeps are reported in Supplementary Section F.1, together with the coverage curves in Figures 3 and 4 and the point-accuracy diagnostics in Supplementary Figure S2. These summaries confirm that robust fits can substantially improve calibration under heavy-tailed or asymmetric misspecification while preserving broadly similar point accuracy in well-specified and mildly misspecified regimes.

### 3.3 Held-out predictive assessment and source heterogeneity

We next turn to held-out predictive assessment based on leave-one-record-out and leave-one-source-out analyses. Detailed numerical summaries for these experiments, including source-level 95% predictive coverage and predictive RMSE ranges, are reported in Supplementary Section F.1; the main text focuses on the diagnostic patterns in the source-level panels.

For glycolysis, the first source-level result is shown in Figure 5. The fixed-effects models under-cover across most nominal predictive levels, especially in the upper tail, indicating that the fitted predictive intervals are again too narrow. Once source-level random effects are introduced, all three observation families move markedly closer to the ideal diagonal. This is an important empirical finding, because it shows that a substantial share of the miscalibration arises not merely from heavy tails or skewness, but from systematic variation across laboratories, assays, or data sources. In this setting, accounting explicitly for source heterogeneity yields a larger gain in predictive reliability than changing the likelihood family alone.

**Figure 5:**
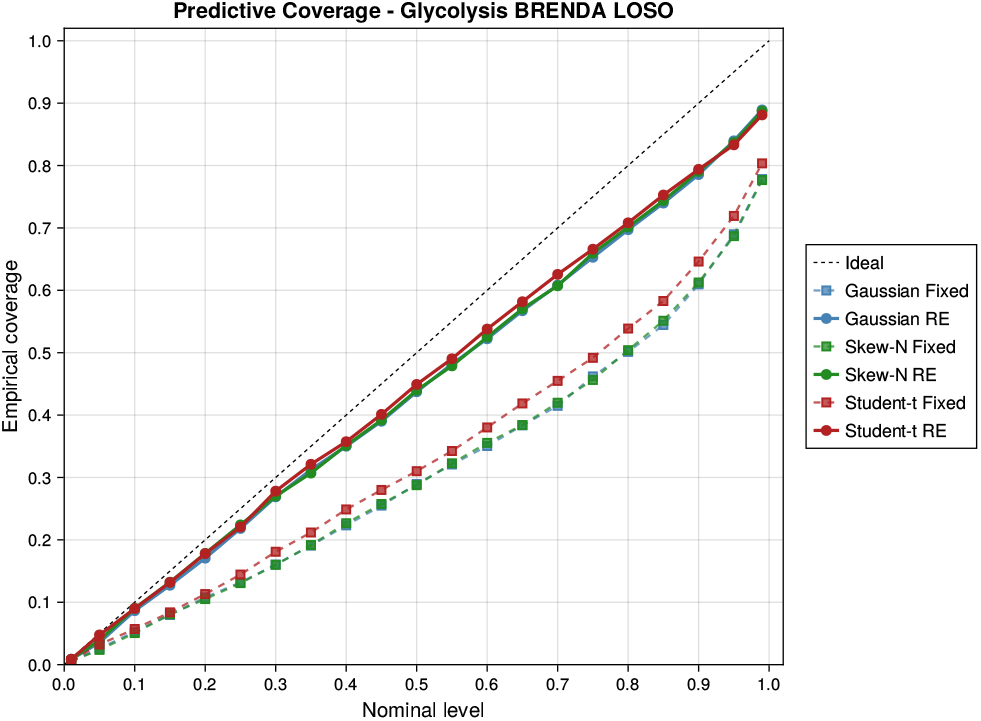
Predictive coverage for glycolysis under leave-one-source-out (LOSO) cross-validation on raw BRENDA data. The dashed diagonal denotes exact calibration. Fixed-effects models under-cover across most nominal levels, especially in the upper tail of the predictive interval spectrum. Introducing source-level random effects moves all three model families substantially closer to the ideal line. Among the hierarchical fits, the Gaussian and skew-normal models are nearly indistinguishable and provide the closest agreement with nominal coverage, while the Student-*t* random-effects model also improves markedly over its fixed-effects analogue and remains competitive, though slightly below nominal at the highest confidence levels. The figure shows that, for glycolysis, the primary calibration gain comes from modeling source heterogeneity rather than from large changes in mean predictive accuracy.

Figure 6 makes the same point from a grouped perspective. Coverage at the nominal 95% level is shown by derived-quantity group together with exact binomial acceptance bands. Random-effects formulations improve calibration most clearly for the small and medium groups, where source-level variability is more visible relative to within-group sample size. This grouped view supports the interpretation from Figure 5: the calibration gain is systematic and not confined to a single subset of the pathway.

**Figure 6:**
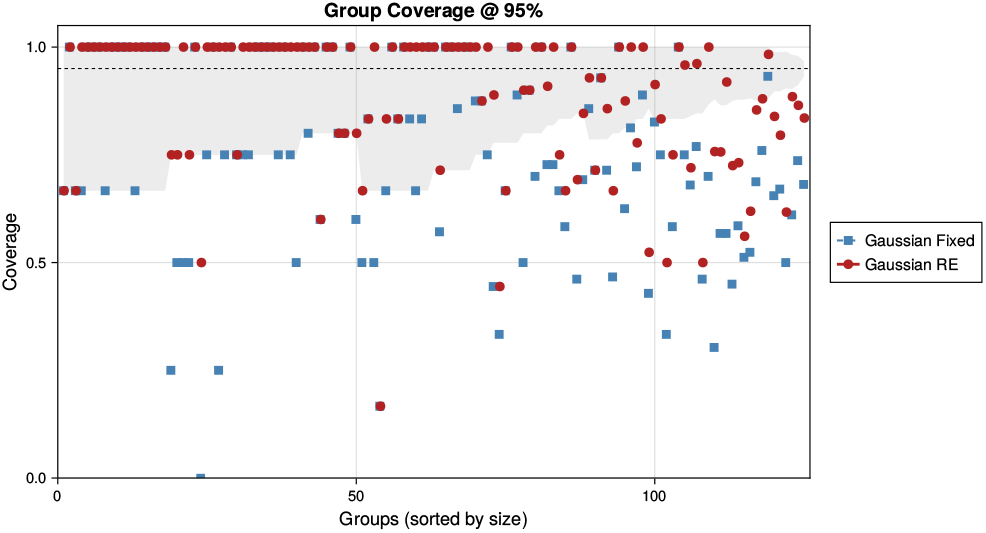
Group-wise predictive coverage for glycolysis at the nominal 95% level. Coverage is shown by derived-quantity group together with exact binomial acceptance bands. The horizontal target line marks ideal calibration, while the group-specific bands indicate the variation expected from finite-sample uncertainty alone, with wider bands for smaller groups. Fixed-effects fits tend to under-cover across several groups, particularly among the small and medium groups, whereas the random-effects formulations move coverage substantially closer to the nominal level. This grouped view complements the aggregate coverage summaries by showing that the gain from hierarchical source effects is systematic across groups rather than driven by only a few large categories.

Full diagnostic panels for glycolysis and the TCA cycle are reported in Supplementary Figures S9 and S10. These panels combine multilevel predictive coverage, interval-width heatmaps, CRPS, WIS, and nominal 95% coverage summaries. They show that average scoring rules change only modestly across fitted models, whereas calibration improves substantially once source-level random effects are included. The interval-width heatmaps further indicate that the added uncertainty is concentrated in a restricted subset of kinetic quantities rather than spread uniformly across the full parameter vector.

Taken together, Figures 5 and 6 and Supplementary Figures S9 and S10 support the same substantive conclusion as the synthetic experiments. Gaussian parameter balancing can remain superficially competitive in average error or RMSE while still failing in predictive calibration. By contrast, robust likelihoods and especially hierarchical source effects deliver the more reliable uncertainty quantification needed for downstream kinetic modelling. To keep the main text focused, we defer the full derivations of the robust and hierarchical models to Supplementary Section D; detailed definitions of interval coverage, predictive coverage, PIT, and predictive scoring rules to Supplementary Sections E.2 and E.4 to E.6; and extended empirical results, including combined held-out diagnostic panels, additional group-wise and multilevel summaries, computational scaling benchmarks, and further pathway-specific sensitivity analyses, to Supplementary Sections F and F.2.

Overall, the empirical evidence is highly consistent across both pathways and across both simulation-based and held-out evaluations. Classical Gaussian parameter balancing works well when its assumptions are approximately correct, but can become overconfident under heavy tails, skewness, or source heterogeneity. Robust likelihoods and hierarchical random effects preserve the thermodynamic consistency of parameter balancing while delivering substantially better-calibrated uncertainty.

#### Computational profiling

We also profiled the computational cost of the Gaussian, Student-*t*, and skew-normal formulations, each fitted with and without random effects. To keep the main Results section focused on calibration and held-out predictive performance, the detailed runtime analyses are reported in Supplementary Section F.2. In brief, runtime increased approximately linearly with both the number of MCMC samples and the number of outer MCEM iterations across the pathway-scale experiments, with random-effects formulations incurring a moderate but predictable overhead relative to their fixed-effects counterparts. Thread-level timing experiments for the parallel cross-validation and synthetic-recovery loops are reported in Supplementary Section F.2.1. For larger runs, the structured variational approximation replaces inner stochastic updates for robust latent variables with deterministic exact-ELBO moment updates while retaining the collapsed sparse Gaussian block; algorithmic details are given in Supplementary Section D.9, with timing results in Supplementary Section F.2.2 and the grouped structured-variational scaling panels in Supplementary Figures S5 and S6.

## 4 Conclusion and future work

The main practical message of this work is that parameter balancing should be evaluated not only by the plausibility of its point estimates, but also by the reliability of its uncertainty quantification. Across both synthetic and held-out pathway analyses, the robust and hierarchical extensions changed posterior means only modestly, but produced substantially better-calibrated uncertainty intervals in the presence of heavy-tailed, skewed, or source-heterogeneous biochemical data. This matters for downstream modelling, where overconfident kinetic parameters can propagate misleading certainty into dynamic simulations, flux analyses, or whole-cell models.

By reformulating parameter balancing as a fully specified Bayesian inference problem, we provide a coherent way to combine kinetic measurements, thermodynamic data, and prior biochemical knowledge, with their relative influence determined by uncertainty rather than heuristic weighting. Posterior predictive checks and leave-one-out validation can also identify poorly explained data points, sources, or groups, which may indicate outliers, assay-specific biases, unmodelled experimental conditions, or limitations of the observation model.

The hierarchical specification used here has a shared source-level intercept across all records from the same publication or curated dataset. This parsimonious model performed well, but richer source-by-parameter random-effects structures may be useful when experimental conditions affect *K*_*M*_, *k*_cat_, or thermodynamic quantities differently. A complementary direction is to encode effects of pH, temperature, or metabolite concentrations mechanistically through thermodynamic corrections, kinetic rate laws, or condition-specific transformations.

The added flexibility comes with moderate computational cost. The Gaussian baseline admits a closed-form posterior and is extremely fast to compute, while the robust Student-*t* and skew-normal extensions require iterative inference but remain practical at pathway scale. Larger whole-cell or genome-scale applications may benefit from more scalable posterior inference strategies, including Hamiltonian Monte Carlo [33–35], variational approximations [36, 37], or sparsity-aware algorithms for structured high-dimensional Bayesian models [38, 39]. The structured variational option in Supplementary Section D.9 is one such approximation: it keeps the conjugate Gaussian update exact conditional on deterministic robust-latent moments, so it is useful for scalability studies while preserving the collapsed sampler as the default route for calibrated posterior summaries.

Our results also highlight the importance of prior specification. Informative priors are often necessary because many quantities are weakly observed or unobserved, and the Bayesian approach makes these assumptions explicit and open to sensitivity analysis. At the same time, strong priors can dominate inference in poorly informed regimes, so careful prior elicitation remains important. Likewise, the framework assumes that the dependency structure encoded by ***Q*** is fixed and correct; extending the model to allow uncertainty in the constraint structure, choice of basis quantities, or kinetic mechanism would be a valuable next step.

The framework also has practical implications for biochemical modelling workflows. Rather than returning only a single balanced parameter set, Bayesian parameter balancing yields posterior distributions and calibrated interval estimates that can be reused as structured prior information in dynamic ordinary differential equation models, steady-state flux models, or whole-cell simulations. The leave-one-record-out and leave-one-source-out analyses further show how the same workflow can distinguish interpolation to an unseen record from prediction for an unseen source. Future work should estimate robustness hyperparameters jointly where possible, scale the approach to yeast or human metabolic networks, and consider more expressive heterogeneous models, including mixture-of-experts extensions [40–43], when data support multiple biochemical regimes.

More expressive heterogeneous models could extend parameter balancing from a globally linear dependency structure to a collection of thermodynamically constrained local experts, each adapted to a different biochemical regime while still accounting for source-specific variation. This direction connects naturally to recent mixture-of-experts theory and scalable model-selection work [44–52]. Substantively, applying the framework to larger real-data case studies will be important for assessing its practical value and for identifying where robust and hierarchical extensions matter most.

Overall, this work moves parameter balancing from a useful but partly heuristic procedure to a statistically principled framework with explicit assumptions, calibrated uncertainty, and robust handling of heterogeneous biochemical data. The resulting approach preserves the biochemical constraints that make parameter balancing attractive while improving its reliability for modern systems biology applications. We expect that this Bayesian re-formulation will make parameter balancing more useful for researchers seeking thermodynamically consistent and uncertainty-aware kinetic parameter estimates from imperfect data.

## Acknowledgments

This project was funded by the Australian Research Council Centre of Excellence for the Mathematical Analysis of Cellular Systems (CE230100001).

## Supplementary Materials

### Supplementary material organization

This Supplementary Material is organised as follows. Section A provides scientific background and interpretive context for the condensed main-text presentation, including the motivation, model intuition, empirical interpretation, and practical implications of the proposed Bayesian parameter-balancing framework. Section B collects notation used in the supplementary derivations and evaluation details. Section C provides additional kinetic and thermodynamic background, including the convenience-kinetics example, Haldane relationship, and the basis-quantity parameterization that motivates the linear dependency structure ***x*** = ***Qq***. Section D gives the technical details for the robust likelihoods, hierarchical source effects, conditional Gaussian representations, fold-wise predictive construction, the collapsed Rao–Blackwellized Monte Carlo expectation– maximization (MCEM) algorithm, and the structured variational approximation for scalable fitting. Section E provides the formal definitions of the simulation-based point-estimation metrics, interval-coverage metrics, held-out predictive diagnostics, scoring rules, probability integral transform (PIT) diagnostics, and parameter-importance summaries used in the main text. Finally, Section F reports the empirical supplementary results, including point-accuracy and calibration summaries for the synthetic misspecification sweeps, raw BRENDA leave-one-record-out and leave-one-source-out diagnostics, collapsed-sampler and structured-variational computational scaling panels, thread-level timing benchmarks, structured-variational timing comparisons, reproducibility workflow details, interval-width analyses, source-level coverage comparisons, and nonparametric tests of predictive performance.

## A Additional explanatory remarks on the main framework

This section is intended as a reader-friendly bridge from the condensed main text to the more technical supplementary material. It does not introduce additional numerical results or assumptions. Instead, it summarizes the modelling intuition, explains how the main empirical comparisons should be read, and points readers to the detailed notation, derivations, diagnostics, and supplementary results that follow.

### A.1 Motivation and Bayesian interpretation

Kinetic models of cellular metabolism require many quantities, including Michaelis constants, turnover rates, elementary kinetic constants, equilibrium constants, and related thermodynamic terms. Literature and database values from resources such as BRENDA, SABIO-RK, and eQuilibrator are valuable but often sparse, noisy, heterogeneous, and mutually inconsistent [1–5]. Parameter balancing addresses this problem by working with independent basis quantities ***q*** and a dependency matrix ***Q***, so that the complete derived vector ***x*** = ***Qq*** satisfies the thermodynamic and algebraic constraints implied by the pathway. This preserves the key strength of classical parameter balancing [20, 21]: missing and conflicting measurements can be reconciled without abandoning physical consistency.

The Bayesian formulation makes this reconciliation explicit. Observed log-kinetic quantities are linked to ***q*** through ***Q***_obs_***q***, while priors 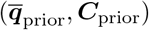 encode biochemical plausibility and dependency structure. In the Gaussian case, the resulting posterior is the conjugate linear update underlying classical parameter balancing, but written as a fully specified generative model. This view clarifies what uncertainty means: unobserved quantities remain constrained by ***Q***, conflicting measurements are weighted by their uncertainty, and posterior or posterior-predictive intervals can be checked against held-out data. The detailed kinetic motivation and notation are given in Sections B and C.

### A.2 Robustness, source effects, and prediction

The robust and hierarchical extensions address two common departures from the Gaussian fixed-effects idealization. Student-*t* and skew-normal likelihoods target record-level errors that are heavy-tailed or asymmetric, respectively. Source-level random effects target systematic differences across publications, laboratories, assay protocols, organisms, temperatures, pH values, and extraction procedures. These extensions modify the observation model but keep the same thermodynamic dependency map ***Q***, so the balanced parameters remain constrained by the biochemical structure.

The fold-wise predictive construction is important for interpreting the raw BRENDA analyses. All fitting and hyperparameter updates are performed using only the training records and the always-included thermodynamic block. Held-out records do not enter the training-fold updates. If a held-out record comes from a source seen during training, prediction can condition on the fitted source effect; if the whole source is held out, prediction integrates over a new source effect from the fitted between-source distribution. The mathematical details of these likelihoods, source effects, and predictive rules are provided in Section D.

### A.3 Empirical interpretation and limitations

The main figures should be read primarily as evidence about uncertainty calibration rather than as evidence that the central balanced parameter values change dramatically. In the synthetic pathway studies, all model specifications recover the broad thermodynamic and kinetic structure, and most posterior means remain similar across likelihood families. The main differences appear in interval width and coverage: Gaussian intervals behave well under near-Gaussian noise but under-cover when the data are heavy-tailed or skewed, whereas the Student-*t* and skew-normal models improve calibration under their corresponding departures. In the raw BRENDA held-out analyses, source-level random effects provide the strongest calibration gain, especially for leave-one-source-out prediction, indicating that between-source heterogeneity is a major component of predictive uncertainty at pathway scale.

The practical message is therefore that parameter balancing should be judged not only by plausible point estimates, but also by calibrated uncertainty. The present hierarchical model uses one shared intercept per source, which is a parsimonious first-order representation of broad source-level shifts. Richer source-by-quantity structures or mechanistic condition corrections may be useful when different assay conditions affect parameter classes in different ways. The supplementary sections below give the formal diagnostics and empirical detail needed to assess these choices: evaluation metrics in Section E, computational and predictive results in Section F, and scalable exact-ELBO structured variational fitting in Section D.9.

## B Supplementary notation and abbreviations

Table 1 summarizes the notation used in the main paper. The supplementary material introduces additional symbols for the robust likelihoods, hierarchical source effects, and fold-wise predictive diagnostics developed in Sections D and E. To keep the presentation compact, we separate these symbols into model-specific notation and evaluation-specific notation.

### B.1 Robust and hierarchical model notation

The symbols in Table S1 are used primarily in the supplementary derivations for the raw-data observation model, variance-matched robust likelihoods, latent-variable augmentations, and hierarchical source effects; see especially Sections D.2, D.3, D.5 and D.6.

### B.2 Evaluation and cross-validation notation

The symbols in Table S2 are used in the supplementary definitions of simulation metrics, held-out predictive assessment, and calibration diagnostics; see Sections E.2 to E.7.

## C Kinetic background and thermodynamic parameterization

This supplement provides additional kinetic and thermodynamic background for the parameter balancing framework used in the main text. Our aim is to make explicit how biochemical rate laws, thermodynamic constraints, and reparameterization together motivate the linear dependency structure ***x*** = ***Qq*** that underlies the Bayesian formulation. We first review the metabolic modelling context, then use a simple reversible reaction to illustrate the role of Haldane constraints, and finally explain how a suitable choice of basis quantities yields a thermodynamically consistent linear parameterization.

### C.1 Metabolic modelling context

Consider a metabolic reaction network, for example a pathway such as glycolysis or the tricarboxylic acid (TCA) cycle. Under standard well-mixed assumptions, the network dynamics can be written as a system of ordinary differential equations (ODEs)

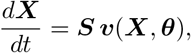

where ***X*** is the vector of metabolite concentrations, ***S*** is the stoichiometric matrix, ***v***(***X, θ***) is the vector of reaction rates, and ***θ*** denotes the collection of kinetic parameters governing those rates. In biochemical kinetics, these parameters typically include catalytic constants, Michaelis constants, enzyme abundances, metabolite concentrations, and thermodynamic quantities such as equilibrium constants or standard chemical potentials.

Although the present work does not fit the ODE dynamics directly, this mechanistic setting determines which biochemical quantities must be estimated and which constraints they must satisfy. In particular, a realistic parameter set must be compatible not only with the observed data, but also with the pathway stoichiometry and the laws of thermodynamics. Parameter balancing addresses this problem by inferring a complete, thermodynamically consistent parameter vector from incomplete, noisy, and heterogeneous measurements.

**Table S1:** Additional notation for robust likelihoods and hierarchical source effects.

| Symbol | Meaning |
| --- | --- |
| $M, D, N$ | Numbers of derived quantities, basis quantities, and raw observations, respectively |
| $m, d, n$ | Indices for derived quantities, basis quantities, and raw observations, respectively |
| $G, g$ | Number of sources and source index |
| $q_d$ | $d$-th component of $\mathbf{q}$ |
| $q_{\text{true},d}$ | $d$-th component of the ground-truth basis vector |
| $\mathbf{x}^* = (x_1^*, \dots, x_N^*)^\top$ | Raw log-transformed observation vector |
| $\sigma_n^2$ | Reported log-scale variance for raw record $n$ |
| $\mathbf{y}_0$ | Thermodynamic data block included in every fold |
| $\mathbf{Q}_0$ | Design matrix for the thermodynamic block |
| $\Sigma_0$ | Covariance matrix for the thermodynamic block |
| $\mathbf{A}_0 = \mathbf{Q}_0^\top \Sigma_0^{-1} \mathbf{Q}_0$ | Information contribution of the thermodynamic block |
| $\mathbf{b}_0 = \mathbf{Q}_0^\top \Sigma_0^{-1} \mathbf{y}_0$ | Information-vector contribution of the thermodynamic block |
| $g(n)$ | Source label of record $n$ |
| $\mu_n$ | Latent mean for record $n$ |
| $\epsilon_n$ | Residual error for raw record $n$ |
| $w_n$ | Latent precision in the record-wise Student-$t$ model |
| $s_n^2 = \frac{\nu-2}{\nu} \sigma_n^2$ | Variance-matched scale in the Student-$t$ model |
| $r_n^2$ | Standardized squared residual for raw record $n$ |
| $\alpha$ | Shape parameter in the skew-normal model |
| $\delta_\alpha = \alpha / \sqrt{1 + \alpha^2}$ | Skew-normal transformation of $\alpha$ |
| $m_1 = \sqrt{2/\pi}$ | Constant in the skew-normal mean correction |
| $c_\alpha^2 = 1 - \delta_\alpha^2$ | Variance adjustment in the skew-normal model |
| $\omega_n^2$ | Variance-matched scale in the skew-normal model |
| $\xi_n$ | Mean-correction term in the skew-normal model |
| $U_n$ | Latent half-normal variable in the skew-normal augmentation |
| $E_n$ | Standard normal latent variable in the skew-normal augmentation |
| $\tilde{x}_n^*(\ell)$ | Effective response given latent variables $\ell$ |
| $w_n(\ell)$ | Effective weight given latent variables $\ell$ |
| $\mathbf{Z}_{-c}$ | Source-incidence matrix on the training fold |
| $\mathbf{W}$ | Diagonal matrix of effective weights |
| $\lambda$ | Prior scale hyperparameter in fold-wise empirical Bayes |
| $\mathbf{q}_{\text{prior}}$ | Fold-wise prior mean in empirical Bayes |
| $\mathbf{C}_0$ | Baseline prior covariance template |
| $\mathbf{P}$ | Block precision matrix for $(\mathbf{q}, \boldsymbol{\eta})$ |
| $\mathbf{P}_q^{\text{marg}}$ | Marginal precision of $\mathbf{q}$ after integrating out random effects |
| $\mathbf{r}, \mathbf{r}_q^{\text{marg}}$ | Right-hand-side vectors in Gaussian updates |
| $\bar{\mathbf{q}}$ | Monte Carlo average of sampled $\mathbf{q}$ values |
| $\mathcal{D}_\lambda$ | Quadratic statistic used in updating $\lambda$ |
| $S_{\eta\eta}^{\text{RB}}$ | Rao-Blackwellized statistic for updating $\tau^2$ |
| $\hat{\tau}^2$ | Fitted between-source variance used in prediction |
| $\hat{\nu}, \hat{\alpha}$ | Fitted robustness parameters |
| $\eta_{\text{new}}^{(s)}$ | Predictive source effect for an unseen source |
| $\mu_n^{(s)}$ | Predictive mean for held-out record $n$ under draw $s$ |
| Gibbs | Gibbs sampler |
| MCEM | Monte Carlo expectation-maximization |

**Table S2:**
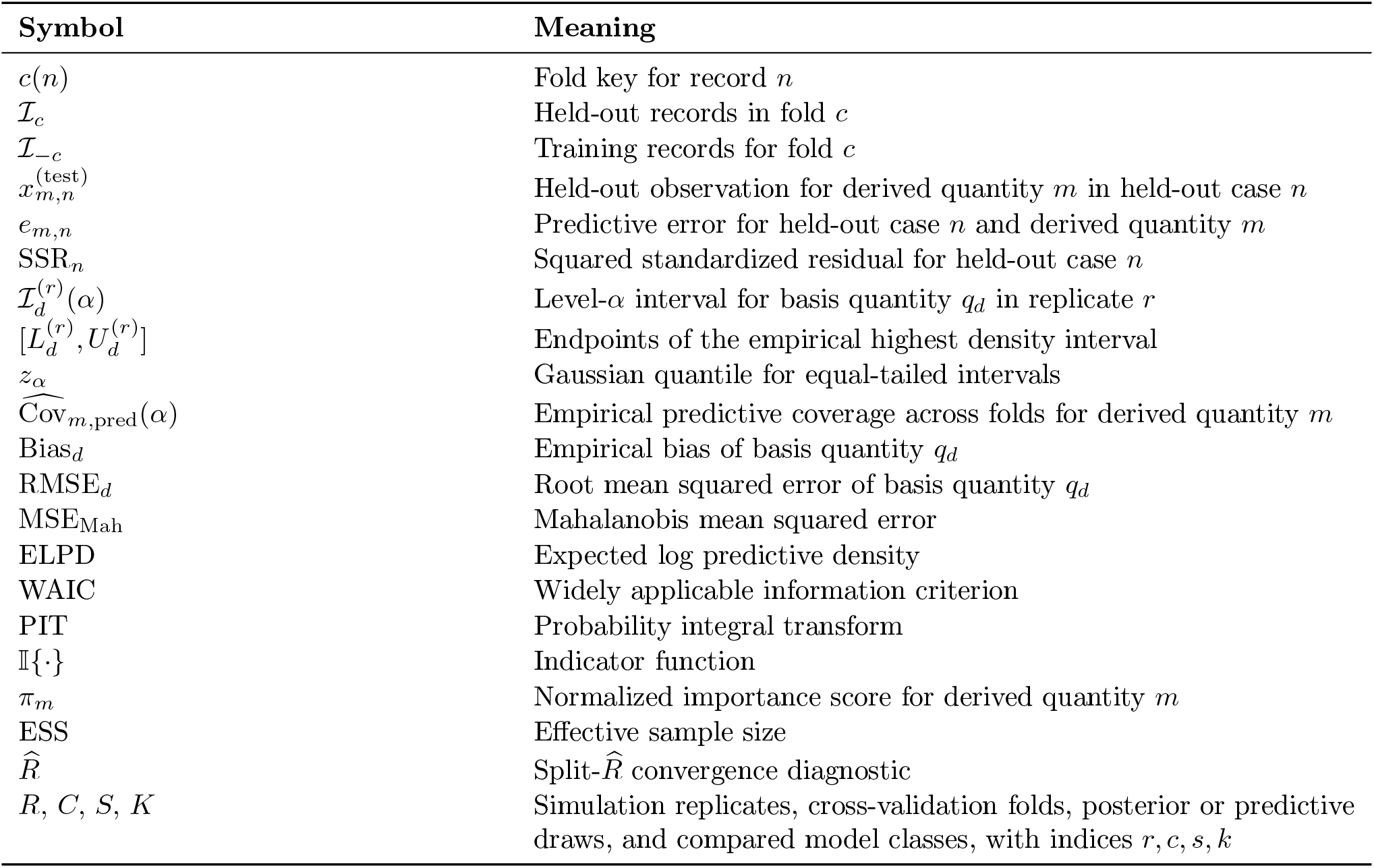
Additional notation for evaluation metrics and fold-wise predictive diagnostics.

| Symbol | Meaning |
| --- | --- |
| $c(n)$ | Fold key for record $n$ |
| $\mathcal{I}_c$ | Held-out records in fold $c$ |
| $\mathcal{I}_{-c}$ | Training records for fold $c$ |
| $x_{m,n}^{(\text{test})}$ | Held-out observation for derived quantity $m$ in held-out case $n$ |
| $e_{m,n}$ | Predictive error for held-out case $n$ and derived quantity $m$ |
| $\text{SSR}_n$ | Squared standardized residual for held-out case $n$ |
| $\mathcal{I}_d^{(r)}(\alpha)$ | Level-α interval for basis quantity $q_d$ in replicate $r$ |
| $[L_d^{(r)}, U_d^{(r)}]$ | Endpoints of the empirical highest density interval |
| $z_\alpha$ | Gaussian quantile for equal-tailed intervals |
| $\widehat{\text{Cov}}_{m,\text{pred}}(\alpha)$ | Empirical predictive coverage across folds for derived quantity $m$ |
| $\text{Bias}_d$ | Empirical bias of basis quantity $q_d$ |
| $\text{RMSE}_d$ | Root mean squared error of basis quantity $q_d$ |
| $\text{MSE}_{\text{Mah}}$ | Mahalanobis mean squared error |
| $\text{ELPD}$ | Expected log predictive density |
| $\text{WAIC}$ | Widely applicable information criterion |
| $\text{PIT}$ | Probability integral transform |
| $\mathbb{I}\{\cdot\}$ | Indicator function |
| $\pi_m$ | Normalized importance score for derived quantity $m$ |
| $\text{ESS}$ | Effective sample size |
| $\widehat{R}$ | Split-$\widehat{R}$ convergence diagnostic |
| $R, C, S, K$ | Simulation replicates, cross-validation folds, posterior or predictive draws, and compared model classes, with indices $r, c, s, k$ |

### C.2 Illustrative reversible reaction example

To illustrate the central ideas, consider a reversible reaction of the form

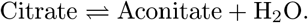

A simple mass-action description is

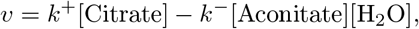

where *k*^+^ and *k*^*−*^ are the forward and reverse rate constants. This form is useful for intuition, but enzyme-catalyzed reactions are more commonly represented by saturation-type laws such as Michaelis–Menten or related reversible kinetics.

Following [13], we adopt convenience kinetics as a flexible reversible rate law that remains biochemically plausible while allowing a systematic thermodynamic parameterization. For the reaction above, one such form is

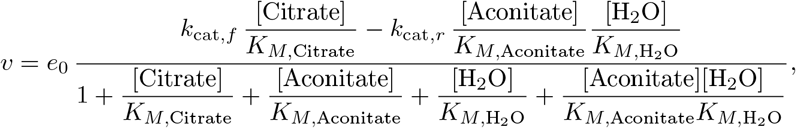

where *e*_0_ is the enzyme concentration, *k*_cat,*f*_ and *k*_cat,*r*_ are the forward and reverse catalytic constants, and the *K*_*M,·*_ terms are Michaelis constants. This example already shows the main challenge: a realistic kinetic description typically involves several interacting parameters, many of which are only partially observed in practice.

### C.3 Haldane relationship and thermodynamic consistency

At equilibrium, the net flux must vanish. For reversible enzyme kinetics, this requirement induces algebraic constraints linking catalytic constants, Michaelis constants, and the reaction equilibrium constant. For the example above, the associated Haldane relationship is

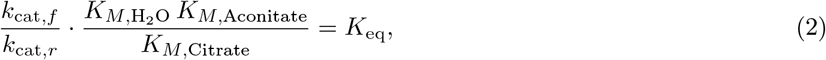

where *K*_eq_ is the equilibrium constant, which may be derived from thermodynamic quantities such as standard Gibbs free energies [3].

Equation Equation (2) shows that the kinetic quantities in Section C.2 are not independent. If one were to draw *k*_cat_ and *K*_*M*_ values independently from literature ranges, the resulting parameter set would generally fail to satisfy Equation (2), and hence would be thermodynamically inconsistent. In small examples this may already lead to implausible kinetics; in larger pathways and genome-scale networks, where many such constraints must hold simultaneously, naive combination of literature values becomes even less reliable.

This is the main motivation for parameter balancing: the task is not merely to smooth noisy measurements, but to infer a complete parameter set that is statistically regularized and physically admissible.

### C.4 Basis quantities and reparameterization

Parameter balancing resolves these dependencies by introducing a set of independent basis quantities ***q***, from which all derived quantities ***x*** are computed. For the reversible reaction above, one may introduce a reparameterization in terms of independent transformed quantities such as a reaction-specific rate parameter and species-specific affinity or binding terms. For example, one may write

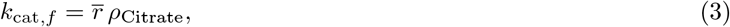

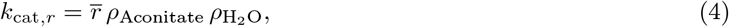

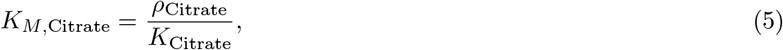

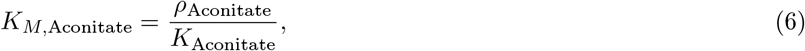

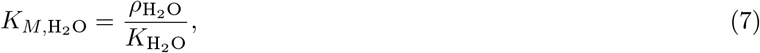

where 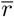 is a reaction-specific rate parameter and the *ρ* and *K* terms encode transformed affinity or binding information. Substituting Equation (3) into Equation (2) yields

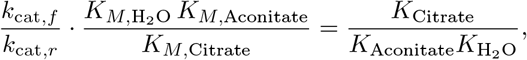

so that the equilibrium constraint is automatically satisfied when

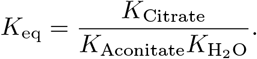

In the bond-graph and modular-kinetics literature [15, 19], 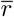 is often interpreted as a transformed rate parameter, whereas the *ρ* terms capture species-dependent energetic contributions and the *K* terms encode effective binding information. The exact interpretation depends on the chosen kinetic formalism, but the common idea is that the original kinetic parameters are replaced by a smaller set of independent quantities from which they can be reconstructed in a thermodynamically consistent way.

The exact form of the reparameterization depends on the kinetic formalism and network representation; see [13, 20] for details. For the present work, the key point is that after appropriate transformation, the resulting dependencies become linear. We therefore write

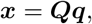

where ***q*** collects the basis quantities, ***x*** collects the full set of derived kinetic and thermodynamic quantities, and ***Q*** is a constant dependency matrix determined by the pathway structure and the chosen thermodynamic parameterization. This linear form is what makes parameter balancing computationally convenient. For example, one row of ***Q*** may encode that log *k*_cat,f_ is the sum of log 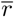 and log *ρ*_Citrate_, whereas another row may encode a different combination for log *k*_cat,*r*_ or log *K*_*M*,Citrate_. More generally, whenever multiplicative kinetic relationships become additive on the transformed scale, they can be represented by suitable rows of ***Q***.

### C.5 Connection to the Bayesian parameter balancing model

The main-text Bayesian formulation builds directly on Section C.4. Observed or curated quantities are represented by the data vector ***x***\*, and the observation map ***Q***_obs_ selects the relevant rows of ***Q***. The Gaussian baseline model is

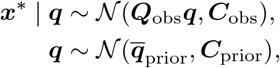

so that parameter balancing becomes Bayesian inference on the basis quantities ***q***. The posterior distribution combines the biochemical constraints encoded by ***Q***, the uncertainty structure encoded by ***C***_obs_, and prior biochemical plausibility encoded by 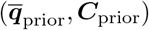.

Posterior summaries are then propagated back to the derived scale through ***Q***. In this way, the thermodynamic and kinetic consistency of the full parameter set is inherited automatically from the basis representation, rather than being imposed after the fact.

### C.6 Practical interpretation

The discussion above illustrates the central rationale of parameter balancing. The goal is not simply to interpolate missing values, but to infer a complete parameter vector that is numerically plausible, statistically regularized, and biochemically admissible. The dependency matrix ***Q*** provides the bridge between these requirements:

1. it encodes pathway-specific biochemical structure,
2. it enforces thermodynamic consistency through the choice of basis quantities, and
3. it enables statistical inference through a linear observation model.

This structure is what makes both the classical Gaussian model and the robust Bayesian extensions in the main text computationally tractable. The robust likelihoods modify the observation model, and the hierarchical extensions add source-specific variation, but the biochemical dependency structure itself remains encoded through the same linear map ***Q***. As a result, the framework can incorporate richer statistical modelling without losing the thermodynamic coherence that motivates parameter balancing in the first place.

## D Detailed robust likelihoods and hierarchical source effects

This section provides the mathematical details underlying the robust and hierarchical extensions introduced in the main text. Throughout, the thermodynamic dependency structure remains unchanged: the basis quantities ***q*** determine the derived quantities through the linear map ***x*** = ***Qq***, and observed quantities are linked to ***q*** through the corresponding observation map. The extensions below modify only the observation model, allowing robustness to heavy tails, asymmetry, and between-source heterogeneity.

### D.1 Why robust likelihoods are needed

The Gaussian parameter balancing model is attractive because it yields closed-form posterior updates and calibrated intervals when the observation model is correctly specified. In practice, however, curated biochemical datasets often contain outliers, heavy-tailed deviations, asymmetric measurement errors, and source-specific biases. Under such misspecification, the Gaussian model can become overconfident: posterior means may remain reasonable, but the corresponding credible intervals may under-cover because the assumed likelihood understates the probability of large or asymmetric deviations.

This issue is especially relevant in parameter balancing, where data are often assembled from multiple laboratories, assay protocols, and reporting standards. In the linear-Gaussian setting, posterior intervals coincide with classical confidence intervals and therefore have the expected coverage under correct specification. This property need not survive under misspecification, which is why empirical coverage and posterior-predictive calibration are central diagnostics in our study. The robust likelihoods considered here are designed to address these effects directly. The Student-*t* model protects against rare but extreme observations through heavier tails, whereas the skew-normal model accommodates directional asymmetry in the residual distribution.

### D.2 Raw-data observation model with source effects

For the raw kinetic-data analysis, let

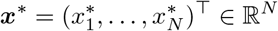

denote the vector of log-transformed records, and let ***Q***\*∈ ℝ^*N ×D*^ be the raw-observation design matrix whose *n*-th row maps record *n* to the corresponding derived quantity. Each record has reported log-scale variance 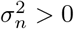, and we write

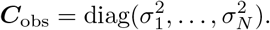

In addition to the raw BRENDA records, we retain the thermodynamic block

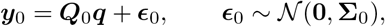

which is present in every fold. Its information contribution is

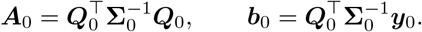

To model between-source heterogeneity, let *g*(*n*) ∈ {1, …, *G*} denote the source label of record *n*, and introduce source-specific random intercepts

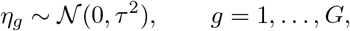

independently. The latent mean for record *n* is then

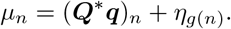

N

Hence *τ* ^2^ quantifies systematic between-source variation beyond the reported within-record measurement variances. This source-effect formulation is the same object used in the raw-data cross-validation protocol, where held-out sources in leave-one-source-out cross-validation are predicted by integrating over a new intercept *η*_new_ ~ 𝒩 (0, *τ* ^2^) rather than by conditioning on any training-fold estimate.

### D.3 Variance-matched robust observation families

We consider three observation families for the within-record residual 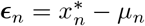, each parameterized so that the variance matches 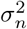. This ensures that differences in performance are not artifacts of trivial rescaling.

#### Gaussian model

The Gaussian baseline is

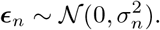

#### Student-*t* model

For heavy-tailed noise, we use a location-scale Student-*t* family. Writing *T*_*n*_ ~ *t*_*v*_ with *v >* 2, we define

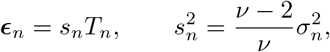

so that

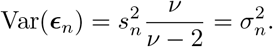

Equivalently, introduce a latent precision *w*_*n*_ through the scale-mixture representation

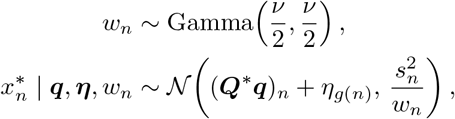

where ***η*** = (*η*_1_, …, *η*_*G*_)^⊤^. The full conditional of *w*_*n*_ is

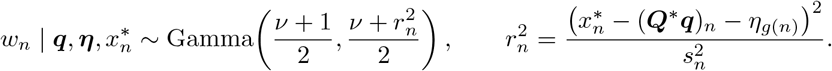

Large residuals therefore induce smaller effective precision, which down-weights outlying observations.

#### Skew-normal model

For asymmetric noise, we use the skew-normal family in Azzalini’s parameterization. Let

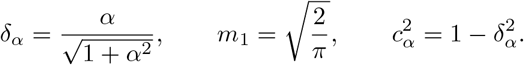

We choose

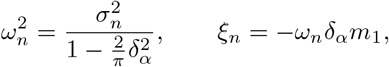

Writing SN for the skew-normal distribution, set

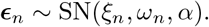

Then

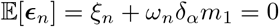

and

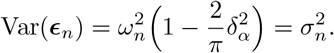

A convenient latent-variable representation is

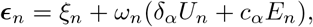

where *U*_n_ ~ | 𝒩 (0, 1) | and *E*_n_ ~ 𝒩 (0, 1) are independent. Conditional on *U*_n_,

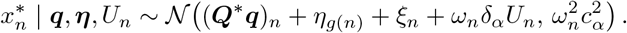

Thus the skew-normal model preserves conditional Gaussianity while allowing directional asymmetry in the measurement error.

If gross outliers are modeled explicitly, we introduce indicators *z*_*n*_ ∈ 0, 1 with *z*_*n*_ ~ Bernoulli(*π*), and replace the conditional mean and variance by *μ*_*n*_ + *z*_*n*_Δ, 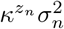, with fixed Δ ∈ R and *κ >* 1. In the analyses where contamination modelling is disabled, one simply sets *z*_*n*_ ≡ 0.

Hence all three families are matched at the same second-moment scale, so differences in performance are not artifacts of trivial rescaling. The Student-*t* and skew-normal families target complementary departures from Gaussian noise: the Student-*t* model captures rare but extreme deviations through heavier tails, whereas the skew-normal model captures directional asymmetry in the measurement error. This contrast is illustrated in Figure S1, where decreasing *v* thickens the tails of the Student-*t* distribution at fixed standard deviation, while increasing |*α*| induces stronger left- or right-skewness in the skew-normal family.

**Figure S1:**
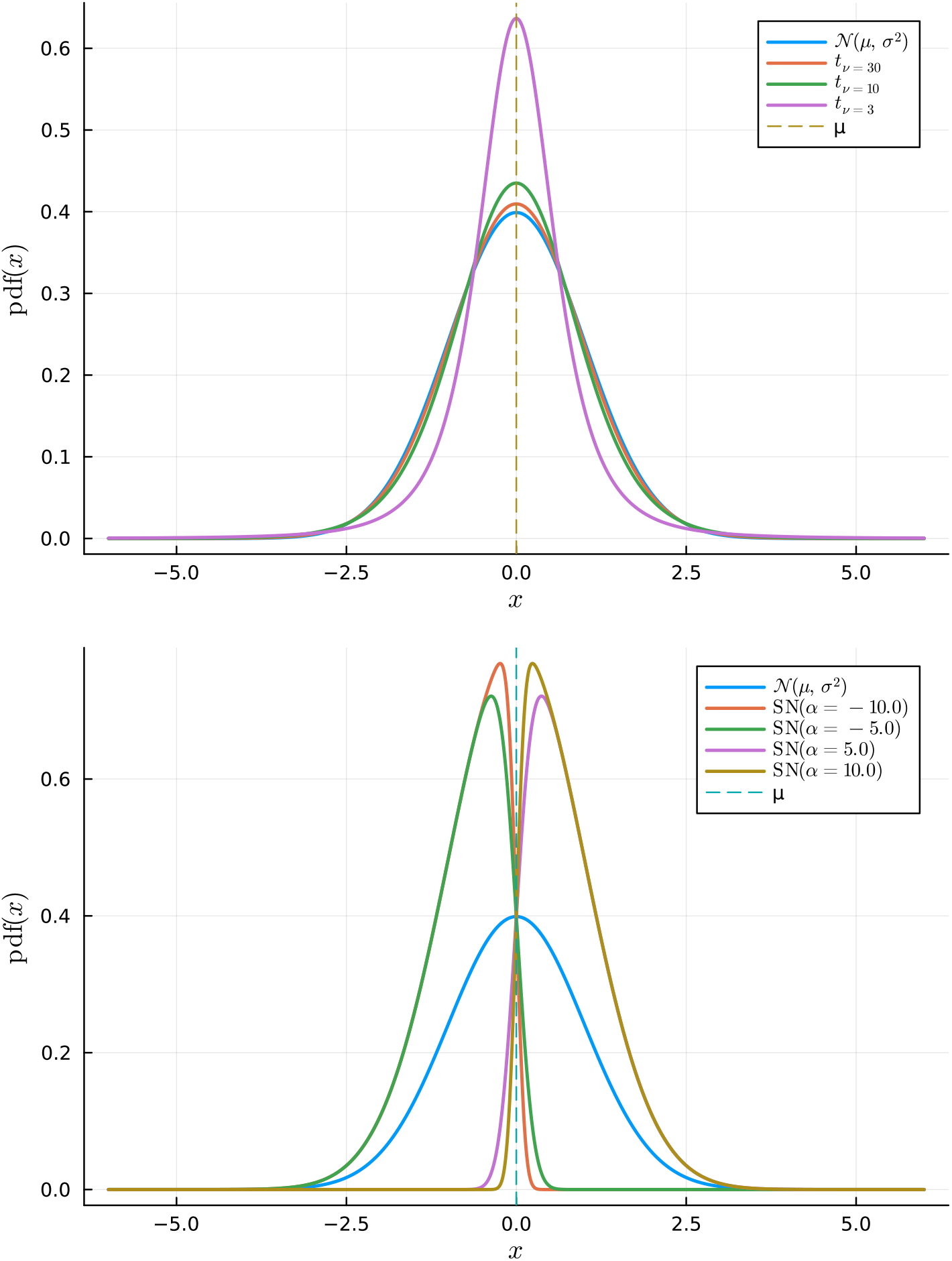
Robust noise families. (a) Heavy tails via Student-*t* (decreasing *v* thickens tails at fixed standard deviation). (b) Asymmetry via skew-normal (sign and magnitude of *α* control the direction and strength of skew while keeping the mean *μ* and scale *σ* fixed).

### D.4 Fold-wise cross-validation on raw kinetic data

We evaluated predictive calibration on raw BRENDA records using leakage-free fold-wise cross-validation. Let *c*(*n*) denote a fold key. Depending on the experiment, *c*(*n*) may be the record index, the source label *g*(*n*), the Enzyme Commission classification of the associated enzyme, or the derived-quantity index. For a fold level *c*, define

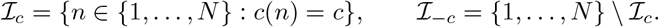

All model fitting, hyperparameter learning, and robustness-parameter handling are performed using only the training records indexed by ℐ_*−*c_, together with the always-included thermodynamic block. No statistic computed from 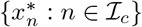 enters any training-fold update.

For Student-*t* and skew-normal models, the robustness-shape parameter is handled within the training fold and never estimated from held-out records. In the raw BRENDA analyses, the Student-*t* degrees of freedom were fixed at *v* = 50, giving a weakly heavy-tailed comparator to the Gaussian model. In the synthetic Student-*t* recovery experiments, the fitted Student-*t* model used the prespecified value *v* = 5, matching the data-generating design. In the controlled degrees-of-freedom sweep, the Student-*t* comparator used the current sweep value. For skew-normal analyses, *α* was initialized at the symmetric value and then updated by a one-dimensional profile step within the training fold, except in the controlled skewness sweep, where the comparator used the current sweep value to isolate the effect of asymmetric misspecification. Conditional on the current or fixed robustness-shape value, the fold-specific empirical-Bayes hyperparameters are estimated by MCEM.

Viewed at the level of exact objective updates, this block-wise scheme is closest to an expectation conditional-maximization (ECM) or expectation/conditional maximization either (ECME) construction: the location and variance hyperparameters (***q***_prior_, *λ, τ* ^2^) are updated by conditional maximisation of an augmented criterion, whereas an estimated robustness-shape parameter is handled by a one-dimensional observed-likelihood or profile-likelihood step. Exact ECM and ECME updates inherit the monotone-ascent property of expectation–maximization (EM) when each conditional step is solved against the appropriate complete- or observed-data objective [53–55]; analogous shape updates are standard in maximum-likelihood fitting of Student-*t* models [56]. Because our expectation-step (E-step) quantities are approximated by Monte Carlo, this monotonicity should be interpreted as an exact-algorithm template rather than as a finite-*S* guarantee for the MCEM procedure.

### D.5 Conditional Gaussian representation

The robust likelihoods admit latent-variable augmentations under which the complete-data likelihood is Gaussian in (***q, η***). Let *𝓁* denote the collection of latent variables introduced by the chosen family. Conditional on *𝓁*, the training-fold likelihood can be written in the unified quadratic form

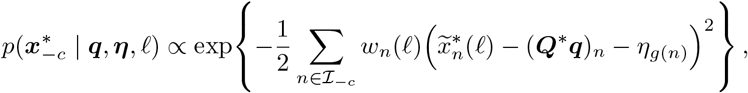

for some effective responses 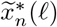 and positive weights *w*_*n*_ (*𝓁*).

For the Gaussian model,

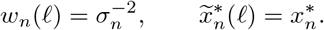

For the Student-*t* model,

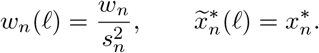

For the skew-normal model,

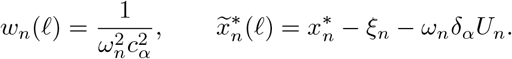

Let 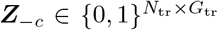 denote the source-incidence matrix on the training fold, so that (***Z***_*−*c_***η***)_*n*_ = *η*_*g*(*n*)_ for *n* ∈ ℐ_*−c*_. Let ***W*** = diag(*w*_*n*_ : *n* ∈ ℐ_*−c*_). Combining Section D.5, the thermodynamic block, and the priors

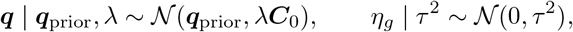

gives the conditional posterior

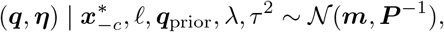

with block precision

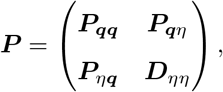

where

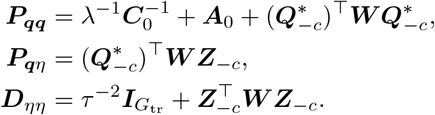

The corresponding right-hand side is

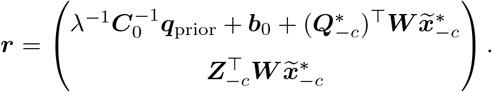

Because each training record belongs to exactly one source, 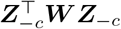 is diagonal, and therefore ***D***_*ηη*_ is diagonal as well. This diagonal structure is the key to the collapsed Gibbs update.

### D.6 Collapsed Gibbs and Rao–Blackwellized MCEM

The direct Gibbs update samples (***q, η***) jointly from the Gaussian conditional above. We instead integrate out ***η*** analytically. Since ***D***_*ηη*_ is diagonal and positive definite, the Schur complement gives the exact marginal precision of ***q***:

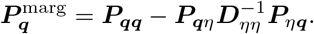

The corresponding marginal right-hand side is

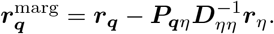

Hence

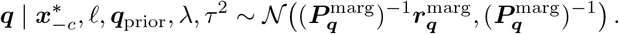

Conditional on ***q***, the source effects become independent:

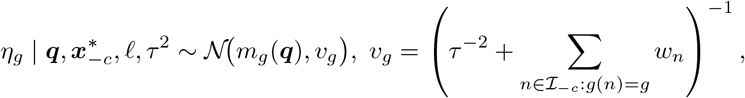

with

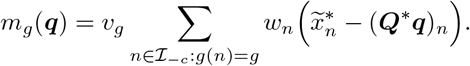

Sampling first from the exact marginal law of ***q*** and then from the exact conditional law of ***η*** | ***q*** therefore produces the same joint conditional distribution as the uncollapsed Gaussian block update.

This is a collapsed Gaussian block update in the sense of collapsed Gibbs sampling [57]: ***η*** is integrated out analytically when sampling ***q***, and is then reconstructed from its exact conditional distribution. This differs from partially collapsed Gibbs samplers, where marginalised and conditional updates may be interleaved and their order must be chosen carefully to preserve the target distribution [58]. For the present Gaussian block, the marginal-then-conditional composition samples from the same joint conditional distribution as the uncollapsed update. Collapsing tractable blocks is often associated with improved operator or autocorrelation behaviour under covariance-ordering comparisons for augmentation schemes and collapsed Gibbs samplers [57, 59], although we do not require a universal ordering for the full sampler.

Within each training fold, empirical-Bayes hyperparameters are estimated by MCEM. The Gaussian prior used by parameter balancing is

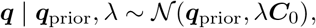

where ***q***_prior_ is the fold-specific empirical-Bayes prior center and *λ >* 0 is a global prior-scale multiplier. To regularize this update, we place the hyperprior

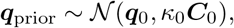

where ***q***_0_ ∈ ℝ^D^ is the fixed thermodynamic prior center used in the classical parameter-balancing initialization, and *κ*_0_ *>* 0 controls the strength of shrinkage toward this baseline. Smaller *κ*_0_ keeps ***q***_prior_ close to ***q***_0_, whereas *κ*_0_ → ∞ gives the unregularized update 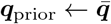.

Given Monte Carlo draws ***q***^(1)^, …, ***q***^(S)^, define

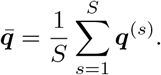

The part of the MCEM objective depending on ***q***_prior_, up to additive constants, is

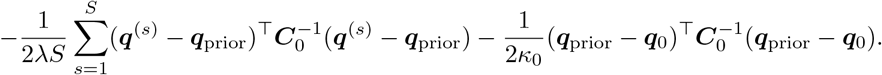

Differentiating with respect to ***q***_prior_ gives

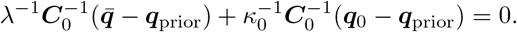

Since ***C***_0_ is positive definite, this is equivalent to

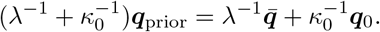

Therefore, the MCEM update for the empirical-Bayes prior center is

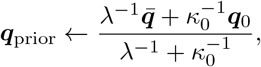

and the inverse-gamma maximum a posteriori (MAP) update for *λ* is

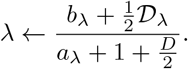

For the source-variance parameter *τ* ^2^, a direct MCEM update would use

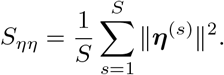

Instead, we use the Rao–Blackwellized estimator

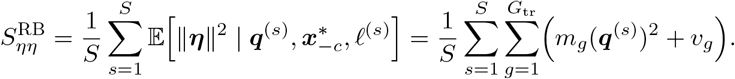

By the law of total variance,

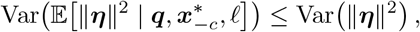

so the Rao–Blackwellized estimator removes the extra Monte Carlo variability generated by sampling ***η*** conditionally on ***q***. More precisely, for a fixed target distribution, the conditional expectation has no larger ordinary variance than the raw plug-in statistic ∥***η***^(s)^ ∥ ^2^, by the Rao–Blackwell inequality [60, 61]. For Markov-chain output, the relevant object is the asymptotic variance of the ergodic average; Rao–Blackwellisation can reduce this quantity under additional transition-kernel and reversibility or covariance-ordering conditions [59, 62], but such dominance is not automatic for arbitrary non-reversible or adaptive chains [63]. We therefore use 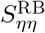 as a variance-reduction device for the *τ* ^2^ update, while treating the finite-*S* MCEM run as a monitored Monte Carlo approximation. The resulting MAP update is

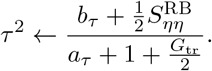

If contamination modelling is enabled, letting

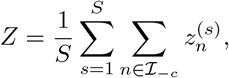

the posterior-mean update for *π* is

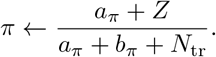

### D.7 Fold-wise predictive evaluation for unseen sources

In fold-wise prediction, if a held-out record belongs to a source represented in the training fold, prediction conditions on the sampled source effect for that source. If the source is absent from training, as in leave-one-source-out cross-validation, prediction instead integrates over a new random effect, 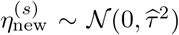 where 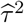 is the fitted between-source variance. This is the appropriate hierarchical predictive rule for unseen sources.

For the Gaussian model with an unseen source, the source effect can be integrated out analytically, giving 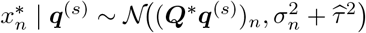. For Student-*t* and skew-normal models, prediction is evaluated by Monte Carlo over the source effect and the likelihood-specific latent variables. This is the predictive construction used in the held-out analyses reported in the main text.

### D.8 Fold-wise MCEM algorithm for robust hierarchical parameter balancing

We detail the fold-wise MCEM procedure in Algorithm S1 used for the robust and hierarchical models described in Sections D.3 to D.7. The Gaussian baseline corresponds to the special case with no latent variables and closed-form updates.

#### Algorithm S1

Fold-wise MCEM for robust hierarchical parameter balancing

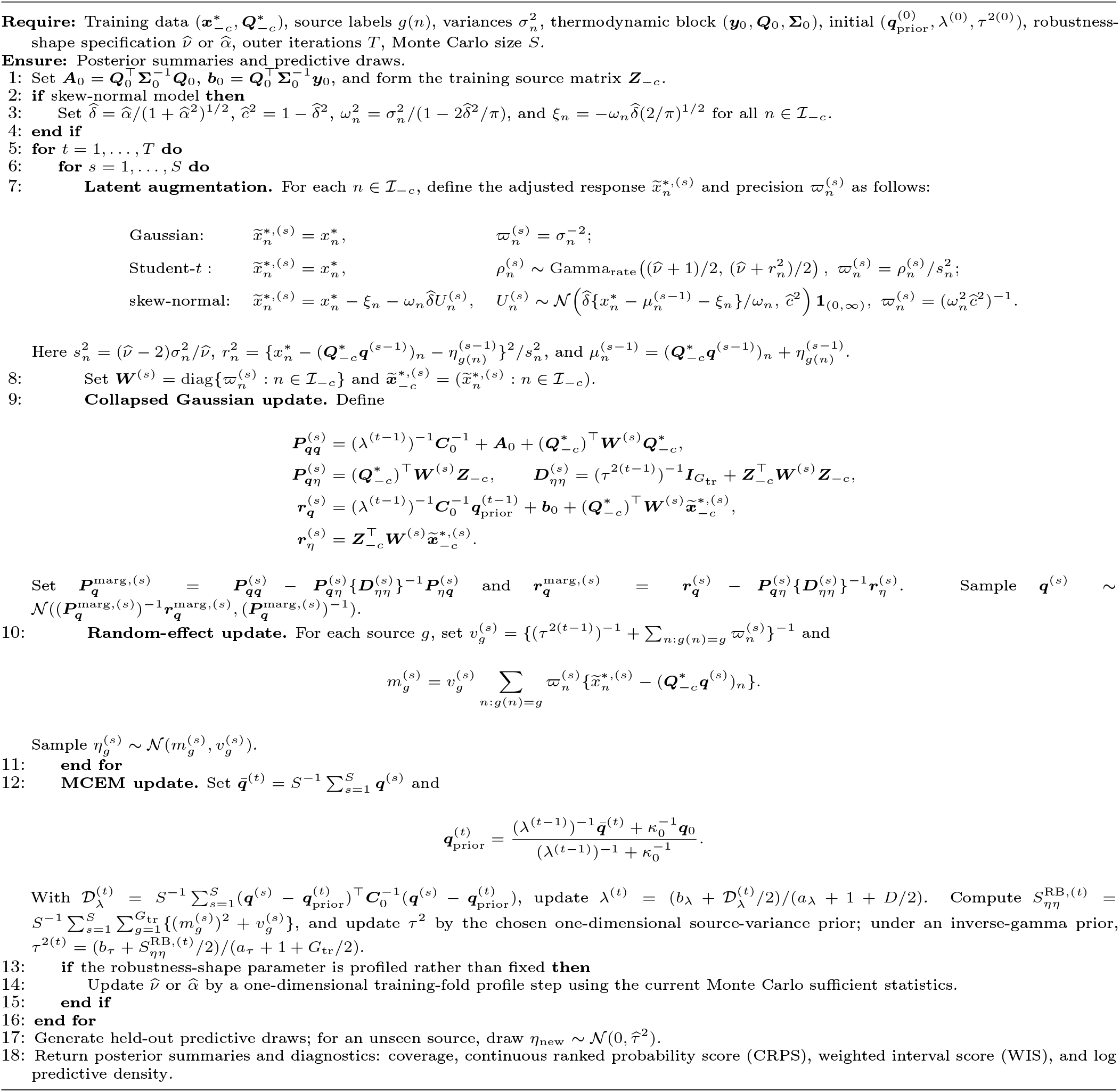

### D.9 Structured variational approximation for scalable fitting

The collapsed MCEM sampler in Algorithm S1 is the default inferential engine for the calibrated posterior summaries reported in the main paper. It accommodates the robust likelihoods by sampling the likelihood-specific latent variables and then exploiting the conditional Gaussian structure of the basis parameters and source effects. Its dominant computational cost is the repeated stochastic refreshing of the Student-*t* scales or skew-normal auxiliary variables within each outer iteration. For the larger sensitivity analyses we therefore use a structured variational approximation that preserves the joint Gaussian dependence between the basis parameters and the source effects, while replacing stochastic refreshes of the robust latent variables by deterministic coordinate-moment updates. The approximation is used as a scalability device, not as the primary source of calibrated posterior uncertainty.

For a training fold with *N* records, write the raw-data observation model as

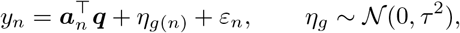

where 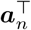 is row *n* of the observation matrix and *g*(*n*) is the source index. The thermodynamic prior contribution is summarized by (***A***_0_, ***b***_0_), and the empirical-Bayes prior is ***q*** *~* 𝒩 (***q***_prior_, *λ****C***_0_). For fixed (***q***_prior_, *λ, τ* ^2^) and fixed robustness parameter *v* or *α*, introduce local variables ***z*** = (*z*_1_, …, *z*_*N*_), with *z*_*n*_ = *w*_*n*_ for the Student-*t* likelihood and *z*_*n*_ = *U*_*n*_ for the skew-normal likelihood. The structured family is

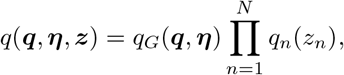

where *q*_*G*_ is a joint Gaussian factor for (***q, η***). The family is deliberately not fully factorized over biochemical quantities: dependence between ***q*** and ***η*** is retained through *q*_*G*_, while only the robust local variables are factorized across records.

The augmented joint density can be written in the schematic form

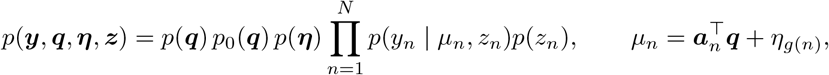

where *p*_0_(***q***) denotes the thermodynamic Gaussian information block. The evidence lower bound is

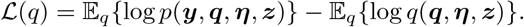

For any factor *q*_*j*_(*x*_*j*_), fixing all other factors gives the coordinate identity

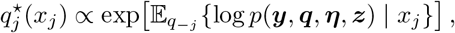

provided the normalizing constant is finite. Equivalently, the *q*_*j*_-dependent part of the ELBO can be written as a constant minus a Kullback–Leibler divergence from *q*_*j*_ to the normalized density on the right, so this coordinate is the exact ELBO maximizer.

For the Student-*t* likelihood, use the variance-matched representation

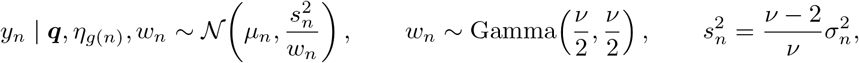

where the Gamma distribution is parameterized by shape and rate. Given the current Gaussian factor, define

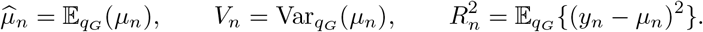

Expanding around 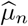 gives

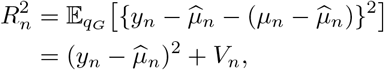

because 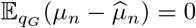. The *q*_*n*_(*w*_*n*_)-dependent expected complete log-density is

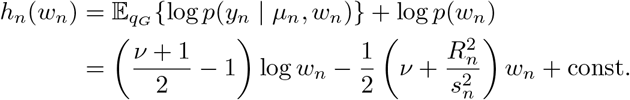

Hence the exact coordinate update is

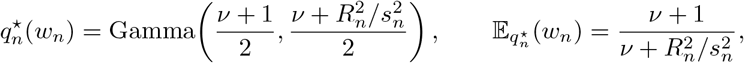

and

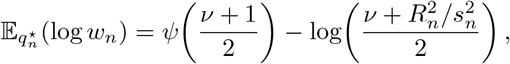

where *ψ* is the digamma function. The predictor variance *V*_*n*_ enters this update because the latent scale *w*_*n*_ multiplies the full squared residual.

For the skew-normal likelihood, define

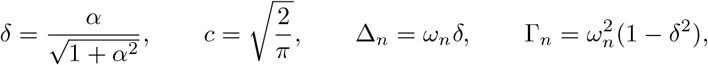

where 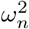 is chosen so that the marginal observation variance equals 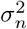. Since *δ*^2^ *<* 1 for finite *α*, Γ_*n*_ *>* 0 whenever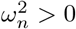. Under the centered skew-normal augmentation,

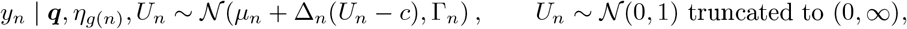

so log *p*(*U*_*n*_ = *u*) = −*u*^2^*/*2 + const for *u >* 0. Fix *q*_G_, set

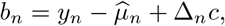

and write

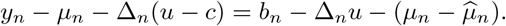

Taking expectation with respect to *q*_G_,

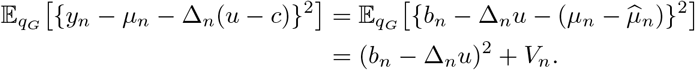

Therefore

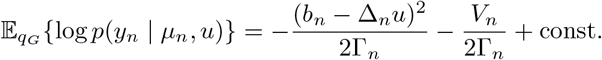

The *V*_*n*_ term affects the ELBO value but is constant in *u*, so it does not affect the local coordinate maximizer. Combining with the half-normal prior gives, for *u* > 0,

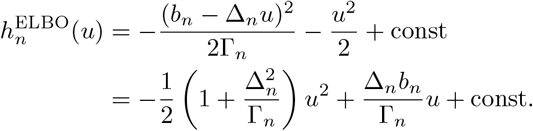

Completing the square yields the exact mean-field ELBO coordinate update

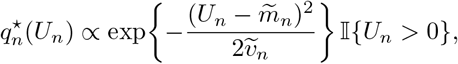

With

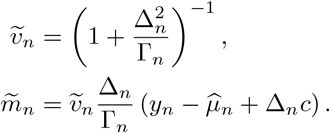

Thus 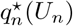 is 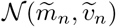 truncated to (0, ∞). The predictor variance *V*_*n*_ is absent from the denominator because the skew-normal auxiliary variable shifts the mean while the conditional variance Γ_*n*_ is fixed; *V*_*n*_ changes the ELBO value but not this local maximizer. All structured-variational numerical results use this exact coordinate update.

For a generic positive-truncated normal *U* ~ 𝒩 (*m, v*) on (0, ∞), let 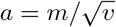. Its first moment is

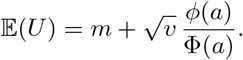

Accordingly,

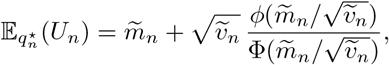

with the ratio evaluated using stable log-tail calculations when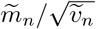 is strongly negative. If 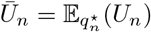, then

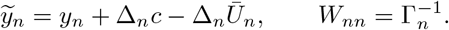

Indeed,

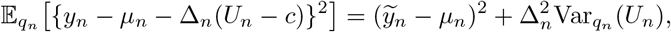

and the second term is constant in (***q, η***). The +Δ_*n*_*c* centering term is therefore required in both the local numerator and the adjusted response.

Fixing the local factors, the Gaussian coordinate update has the common quadratic form

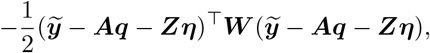

Where

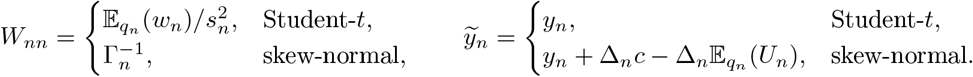

Combining this likelihood quadratic with the empirical-Bayes prior, thermodynamic block, and source-effect prior gives

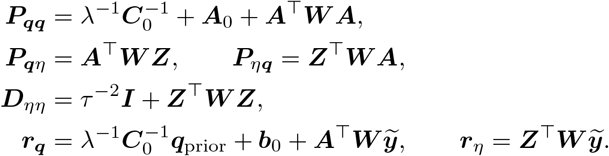

The joint Gaussian factor has precision

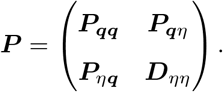

For any nonzero (***u, v***),

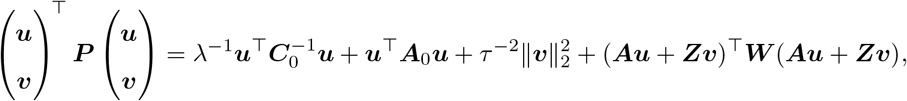

which is strictly positive when *λ >* 0, *τ* ^2^ *>* 0, ***C***_0_ ≻ 0, ***A***_0_ ⪰0, and the diagonal entries of ***W*** are positive. Thus the Gaussian coordinate is well defined.

Because each record belongs to one source, ***Z***^**⊤**^***WZ*** and hence ***D***_*ηη*_ are diagonal. Source effects can therefore be collapsed by the Schur complement:

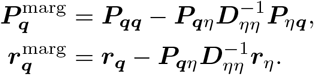

The marginal Gaussian factor for the basis vector is

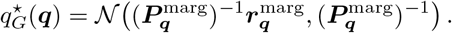

The conditional source-effect factor is

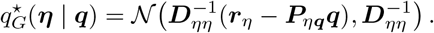

This is the same sparse Schur-complement algebra used by the collapsed sampler; sparse row supports are used when assembling ***A***^**⊤**^***WA***, and the only dense factorization is the *D* × *D* Cholesky factorization of 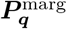.

#### Algorithm S2

Structured variational approximation for one training fold

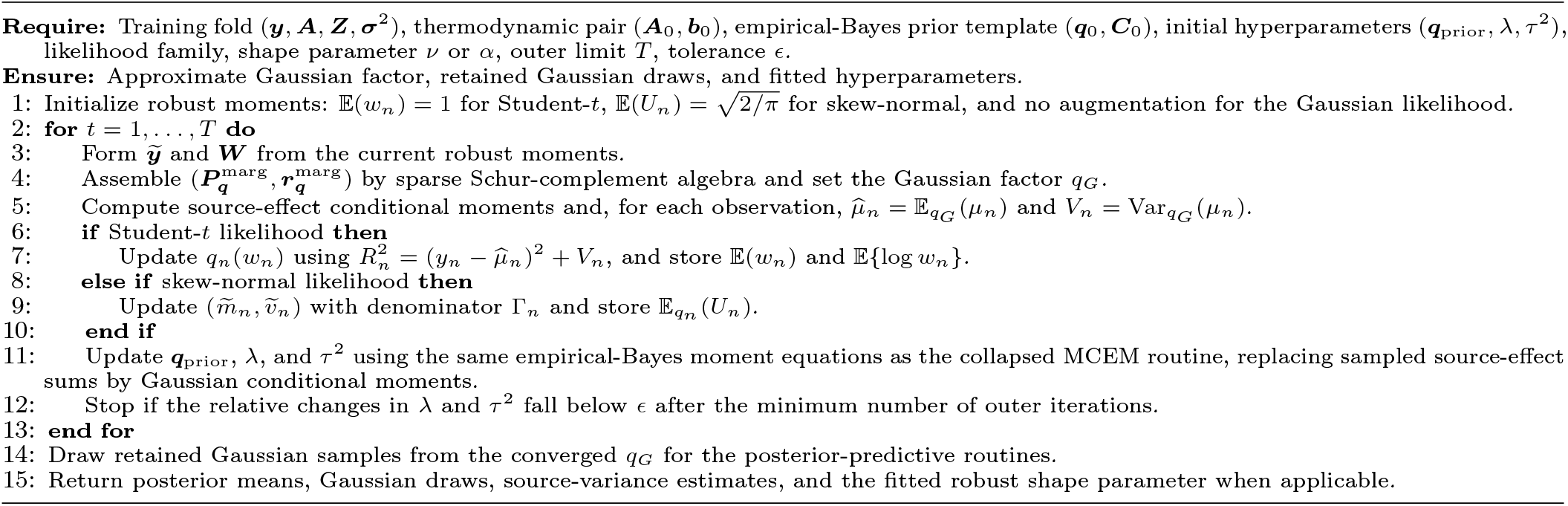

The numerical procedure checks dimensions, missing or non-finite values, positive measurement variances, sampler counts, supported likelihood labels, and unsupported contamination-mixture specifications before fitting. The approximation retains the posterior dependence among basis quantities and source effects through the collapsed Gaussian block, replacing only the robust latent-variable refresh by deterministic ELBO-coordinate moments. This design is appropriate for parameter balancing because the dominant structural information is linear and Gaussian once the robust augmentation is conditioned out. The trade-off is that deterministic moment updates can understate tail uncertainty relative to the collapsed sampler, especially under strong misspecification; consequently, the collapsed sampler remains the default for calibration results, while structured VI is reserved for scalability checks and sensitivity analyses.

### D.10 Practical computational remarks

In practice, the conditional Gaussian updates are numerically most stable with Cholesky-based linear algebra rather than explicit matrix inversion. For the Student-*t* model, smaller values of ν typically increase posterior dependence between ***q*** and the latent scales *w*_*n*_, which can slow mixing and require longer Monte Carlo runs. More generally, robust and hierarchical fits should be monitored using standard Markov chain diagnostics such as effective sample size and split-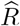. These checks were used throughout our experiments to ensure that the additional flexibility of the robust models did not compromise posterior reliability.

## E Evaluation metrics and held-out predictive diagnostics

This section provides the formal definitions of the evaluation metrics and held-out predictive diagnostics used in the main text. Our goal is to distinguish point-estimation accuracy from uncertainty calibration, and to assess whether posterior or posterior-predictive intervals behave as advertised under model misspecification and source heterogeneity.

### E.1 Simulation-based point-estimation metrics

Let ***q***_true_ denote the ground-truth basis vector used to generate synthetic data, and let 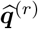 and 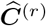 denote the posterior mean and covariance from replicate *r*, for *r* = 1, …, *R*. For basis quantity *d*, we report the empirical bias

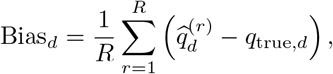

and the root mean squared error

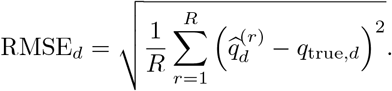

To summarize multivariate error relative to posterior uncertainty, we also use the Mahalanobis mean squared error

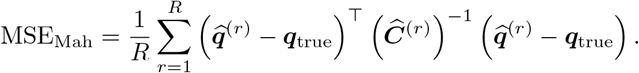

This quantity is useful because it reflects both point-estimation error and the scale of posterior uncertainty.

### E.2 Interval coverage in simulation

A central target of this work is calibrated uncertainty. For each basis quantity *q*_*d*_, replicate *r*, and nominal level *α*, let 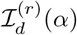 denote the corresponding posterior interval. We define empirical coverage as

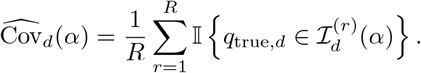

If the posterior intervals are well calibrated, then 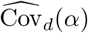 should be close to *α*.

For Gaussian posterior summaries, equal-tailed Wald-style intervals are

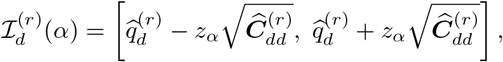

where *z*_*α*_ is the corresponding Gaussian quantile. For example, *z*_0.68_ = 1 and *z*_0.95_ ≈ 1.96.

For non-Gaussian posterior draws, we also consider highest density intervals (HDIs). Let 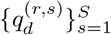 denote posterior samples for basis quantity *d* in replicate *r*. Sorting these as

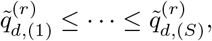

and setting *k* = ⌈*αS⌉*, the empirical HDI is defined as the shortest interval

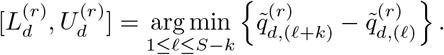

Coverage is then computed as

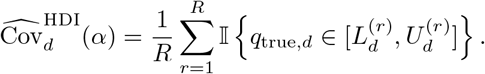

To visualize whether departures from nominal coverage exceed Monte Carlo variation, we also use a binomial reference band. Under exact calibration, the standard error of 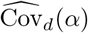 is approximately

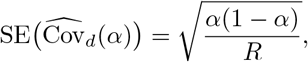

so an approximate 95% reference band is

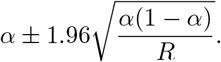

### E.3 Held-out predictive assessment

We assess predictive calibration using fold-wise cross-validation. In the record-level analysis, each fold holds out one raw BRENDA record. In the source-level analysis, each fold holds out all observations from one laboratory, study, or curated source. The latter is the primary stress test for source heterogeneity, whereas record-level leave-one-out is a complementary sensitivity analysis for point-predictive accuracy and calibration when other records from the same source may remain in the training set. Let *c* = 1, …, *C* index folds and let *n* = 1, …, *N* index the held-out predictive cases after expanding all folds. For the fold *c*(*n*) containing held-out case *n*, the model is fitted on the training data 𝒟_*\c*(*n*)_, yielding a posterior or posterior sample over ***q***, from which we construct the posterior predictive distribution

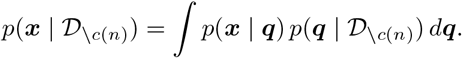

Let 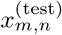 denote the *n*-th held-out observation for derived quantity *m*, and let 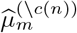 and 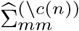 denote the predictive mean and variance from the corresponding training fold. The held-out predictive error is

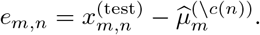

We summarize predictive bias and predictive RMSE across held-out cases by

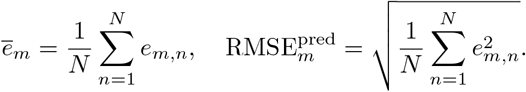

### E.4 Predictive interval coverage

For predictive calibration, we compute the empirical coverage of posterior-predictive intervals. Let 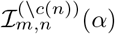 denote the level-*α* predictive interval for held-out case *n* and derived quantity *m*. Then

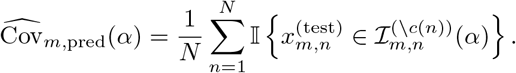

In the Gaussian predictive case,

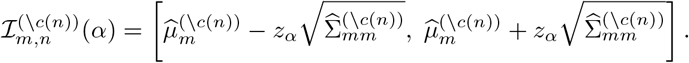

For sample-based predictive distributions, we compute HDIs or empirical equal-tailed intervals directly from predictive draws.

This metric is especially important in the present setting because a model may achieve low predictive RMSE yet still produce systematically under-dispersed predictive intervals. The held-out coverage diagnostics therefore separate point-predictive accuracy from uncertainty calibration.

### E.5 Posterior-predictive scoring rules

To complement coverage, we also report proper scoring rules for predictive distributions. Let *F*_*m,n*_ denote the predictive distribution for held-out observation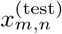.

The log predictive density (LPD) is

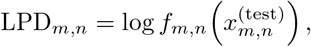

where *f*_*m,n*_ is the corresponding predictive density. Aggregating over held-out cases gives the leave-one-out (LOO) log predictive density,

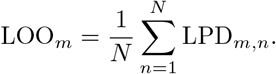

This targets the expected log predictive density (ELPD), which underlies model comparisons such as WAIC and LOO.

We also report the continuous ranked probability score (CRPS), defined for a univariate predictive distribution *F* and realized value *x* as

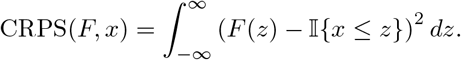

CRPS rewards both accurate location and appropriate dispersion, with smaller values indicating better predictive performance.

In addition, we use the weighted interval score (WIS), which summarizes interval forecasts across one or more nominal levels. For a central (1 − *α*)-interval with lower and upper endpoints (*l, u*), the interval score is

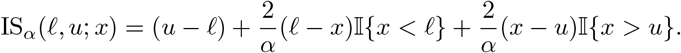

WIS averages such interval scores across several *α* values and optionally includes the predictive median. As with CRPS, smaller values are better.

### E.6 PIT diagnostics

For predictive calibration we also examine the PIT. For held-out observation 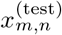 under predictive cumulative distribution function (CDF) *F*_*m,n*_, define

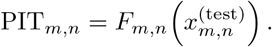

If the predictive distribution is correctly calibrated, then the PIT values should be approximately uniformly distributed on [0, 1]. U-shaped histograms indicate under-dispersion, hump-shaped histograms indicate over-dispersion, and skewed histograms indicate systematic bias.

### E.7 Parameter-importance diagnostic

To summarize how strongly predictive performance varies across quantities, we use a normalized importance diagnostic based on the variability of held-out predictive errors. For derived quantity *m*, define

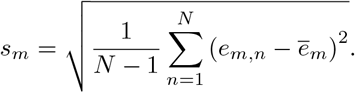

The normalized importance is then

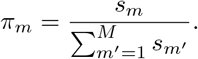

Large values of *π*_*m*_ indicate quantities whose predictive performance varies strongly across folds and hence may be especially sensitive to source heterogeneity or model misspecification.

### E.8 Interpretation of the diagnostic panels

The combined diagnostic panels in the main text and supplement summarize several complementary aspects of model performance:

1. **Coverage curves** compare empirical predictive coverage to nominal interval levels. Curves below the diagonal indicate under-coverage and overconfidence.
2. **Interval-width heatmaps** show where models increase uncertainty. This helps distinguish broad inflation of uncertainty from localized widening for a small subset of parameters.
3. **CRPS and WIS summaries** assess predictive sharpness and accuracy, but do not by themselves guarantee calibration.
4. **PIT histograms** diagnose systematic miscalibration of the predictive distribution.
5. **Coverage at** 95% provides a single headline calibration summary, but should be interpreted together with multilevel coverage curves.

Taken together, these diagnostics explain why robust and hierarchical models may appear similar to Gaussian fits in point-predictive scores while still offering substantial gains in uncertainty reliability.

## F Supplementary empirical results

This section collects additional empirical results that complement, rather than repeat, the main paper. While the main text focuses on the core synthetic-misspecification results and the primary held-out comparisons, the figures below provide a fuller view of three issues that are especially relevant for real biochemical databases: calibration on raw BRENDA records, the sharpness–calibration trade-off across model families, and the localization of uncertainty expansion under hierarchical source effects. We also report source-level comparisons and computational scaling to show that the improved predictive calibration of the hierarchical models is both systematic and computationally manageable.

### F.1 Point accuracy and predictive calibration summaries

To make the distinction between point accuracy and uncertainty calibration explicit, the full 200-replicate synthetic misspecification sweeps record mean absolute bias, mean RMSE, Mahalanobis mean squared error, and mean 95% interval width at each tail-heaviness or skewness setting. These summaries are paired with the coverage curves in main-text Figures 3 and 4 and with the point-accuracy summaries in Figure S2, which reports the point-accuracy diagnostics across synthetic misspecification sweeps. The degrees-of-freedom sweep varies the canonical Student-*t* scale and fits the robust comparator with the corresponding sweep variance; the skewness sweep uses the location-parameterized skew-normal family and fixes the robust comparator at the current sweep value. Under moderately heavy-tailed noise (*ν* = 4), the Student-*t* fit increased 95% parameter coverage from 0.891 to 0.956 in the TCA cycle and from 0.876 to 0.948 in glycolysis, while mean RMSE changed only from 2.752 to 2.691 and from 1.471 to 1.415, respectively. At the near-Gaussian endpoint (*ν* = 50), the Gaussian and Student-*t* fits had essentially the same RMSE in both pathways (2.146 versus 2.148 in the TCA cycle; 1.065 versus 1.058 in glycolysis). Under strong skewness, the skew-normal fit also improved point recovery as well as calibration; for example, at *α* = 10, TCA coverage increased from 0.738 to 0.939 and glycolysis coverage from 0.753 to 0.939. Thus, low point error alone is not sufficient to justify a parameter-balancing procedure. In our setting, the robust models are preferable primarily because they recover substantially better calibrated intervals, while sacrificing little or no point accuracy and sometimes improving it under stronger asymmetric misspecification.

We next turn to held-out predictive assessment based on leave-one-record-out and leave-one-source-out analyses. These experiments address a complementary question to the synthetic coverage studies: not only whether parameter intervals are calibrated under simulated misspecification, but whether the fitted models produce reliable predictive uncertainty for real heterogeneous biochemical records. Leave-one-record-out evaluates interpolation to an unseen record while usually retaining other records from the same source in the training set. Leave-one-source-out is more stringent for source heterogeneity, because the model must predict all records from an unseen publication or curated source. In the source-level analyses, fixed-effects 95% predictive coverage ranged from 0.680 to 0.698 in the TCA cycle and from 0.685 to 0.718 in glycolysis, whereas the random-effects fits reached 0.831–0.848 and 0.834–0.839, respectively. These gains occurred with little change in predictive RMSE: across the six source-level fits, RMSE ranged from 3.226 to 3.255 in the TCA cycle and from 2.969 to 3.006 in glycolysis.

### F.2 Computational scaling benchmarks

To complement the calibration results in the main text, we report empirical runtime scaling for the pathway-scale fits. We varied the number of outer Monte Carlo expectation–maximization (MCEM) iterations, the number of posterior samples, the basis dimension *D*, the number of sources *G*, and the total number of observations *N*. These measurements isolate the computational cost of adding robust likelihoods and source-level random effects while keeping the thermodynamic dependency structure fixed. The collapsed-sampler panels are split into pathway-level algorithmic grids and structural-size grids in Figures S3 and S4; the exact-ELBO structured variational panels use the same grouping in Figures S5 and S6. Within each figure, at most two panels are placed in a row so that the axis labels, legends, and panel annotations remain readable.

Across the experiments, runtime increases regularly over the tested ranges. Fixed-effects models remain cheaper than the corresponding random-effects models because the hierarchical variants require additional updates for source-specific effects. However, the scaling remains stable and predictable across all panels in Figures S3 to S6, indicating that the robust and hierarchical extensions are computationally practical at pathway scale.

The same benchmark grids were also evaluated with the structured variational inference option described in Section D.9. The plotting settings, model ordering, and benchmark grids match the collapsed-sampler scaling panels; only the posterior inference method is changed. Grouping the panels by inference method and scaling factor makes the comparison compact while preserving the corresponding grid-level comparisons.

The method comparisons in these scaling figures should be read as computational, rather than statistical, performance summaries. The Gaussian fixed-effects fit is the computational reference because its posterior update is closed form. Robust fixed-effects fits remain close to this reference under the structured variational approximation, since the Student-*t* scale variables and skew-normal auxiliaries are updated by deterministic conditional moments rather than repeated stochastic refreshes. The main additional cost comes from the random-effects formulations, which must update and propagate source-level latent effects. Consequently, the separation between fixed-effects and random-effects curves is usually more informative than small crossings among the Gaussian, Student-*t*, and skew-normal fixed-effects curves, especially at the shortest runtimes where wall-clock noise is visible.

Across pathway-size, observation-count, source-count, outer-iteration, and posterior-sample grids, the qualitative ordering is stable: adding source heterogeneity increases runtime, whereas adding robust likelihoods without source effects has a smaller computational effect under structured variational inference. Thus the figures support the same practical conclusion as the accuracy and calibration diagnostics: robust and hierarchical models are computationally feasible at pathway scale, but method choice should be based primarily on calibration and predictive reliability rather than runtime alone.

**Figure S2:**
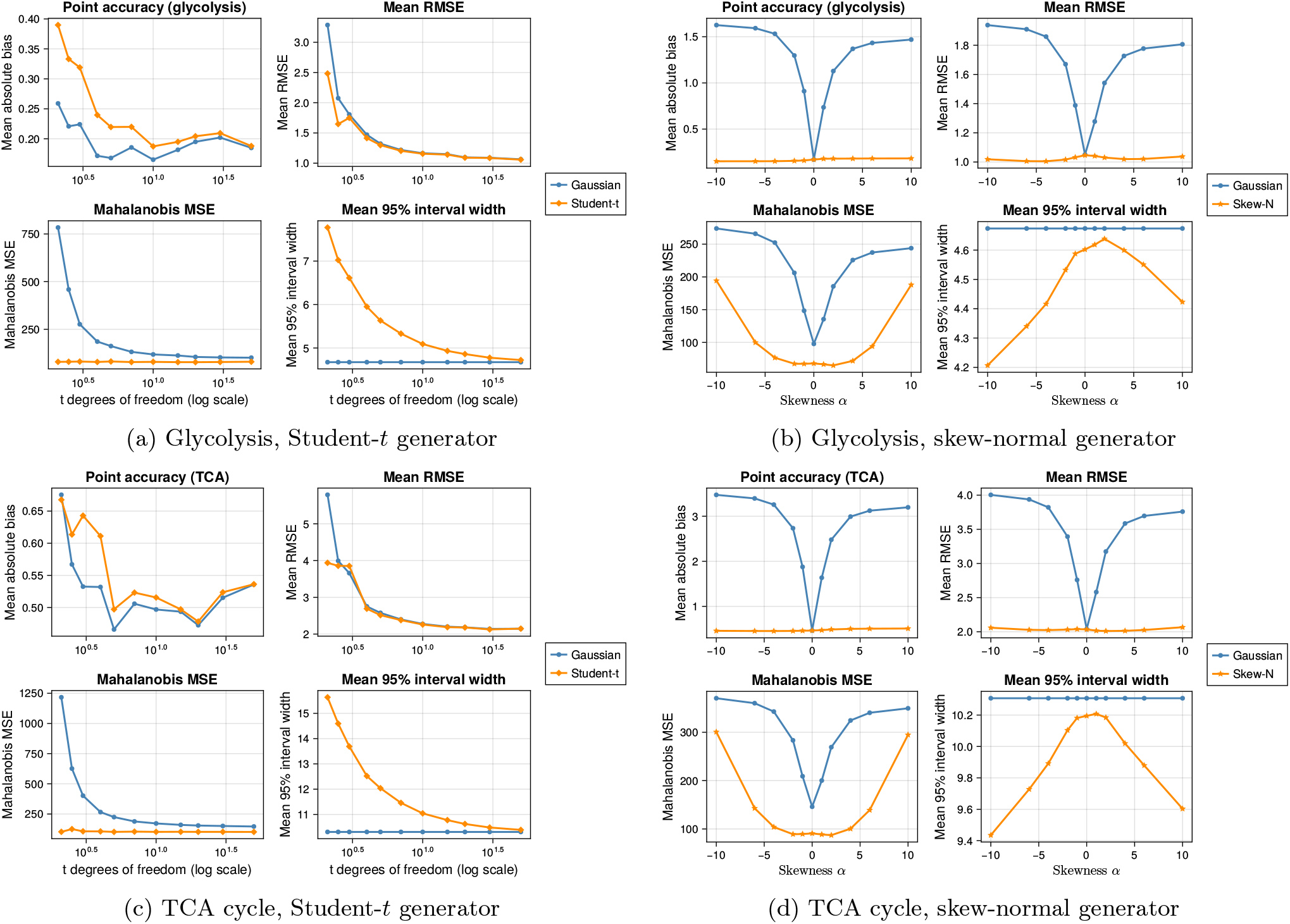
Point-accuracy diagnostics across synthetic misspecification sweeps. Each panel summarizes the 200-replicate experiments by mean absolute bias, mean RMSE, Mahalanobis mean squared error, and 95% interval width. The panels complement the coverage curves in main-text Figures 3 and 4: point error and interval calibration do not always move together, so a model can remain competitive in RMSE while still producing intervals that under-cover.

#### F.2.1 Thread-level parallel execution

The main computational bottlenecks in the reproducibility workflow are independent cross-validation folds and independent synthetic recovery replicates. The thread-level benchmark therefore measures parallel execution over leave-one-record-out folds, leave-one-source-out folds, and synthetic recovery replicates, with deterministic per-fold or per-replicate random seeds. The timing experiment in Table S3 was run with Basic Linear Algebra Subprograms (BLAS) restricted to one thread, so that the measured speedups reflect task-level parallelism rather than nested linear-algebra threading.

#### F.2.2 Structured variational inference option

The primary analyses use the collapsed Rao–Blackwellised MCEM sampler described above. This sampler exploits conjugacy by marginalising the source effects when drawing the basis vector ***q***, then sampling the source effects conditionally. The robust Student-*t* and skew-normal likelihoods nevertheless require repeated stochastic updates of the augmented latent variables inside each MCEM iteration. To support larger sensitivity analyses, we therefore evaluated a structured variational inference (VI) approximation. This option keeps the same observation model and the same sparse collapsed Gaussian algebra, but replaces the inner Gibbs updates for robust latent variables with deterministic conditional-moment updates. The Gaussian baseline is unchanged, because its posterior is already available in closed form.

For the Student-*t* model, let 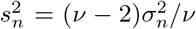 denote the variance-matched scale and let 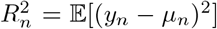, where the expectation includes posterior uncertainty in the linear predictor. The structured update uses

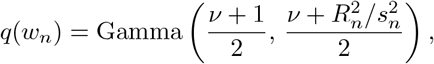

under the shape-rate convention, with 𝔼 [*w*_*n*_] and 𝔼 [log *w*_*n*_] used in the subsequent Gaussian block and shape-parameter M-step. For the skew-normal model, the positive truncated-normal auxiliary variable is set to the mean of its exact ELBO-coordinate truncated-normal approximation. If 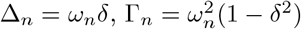, and 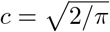 for the mean-parameterized skew-normal model, then

**Figure S3:**
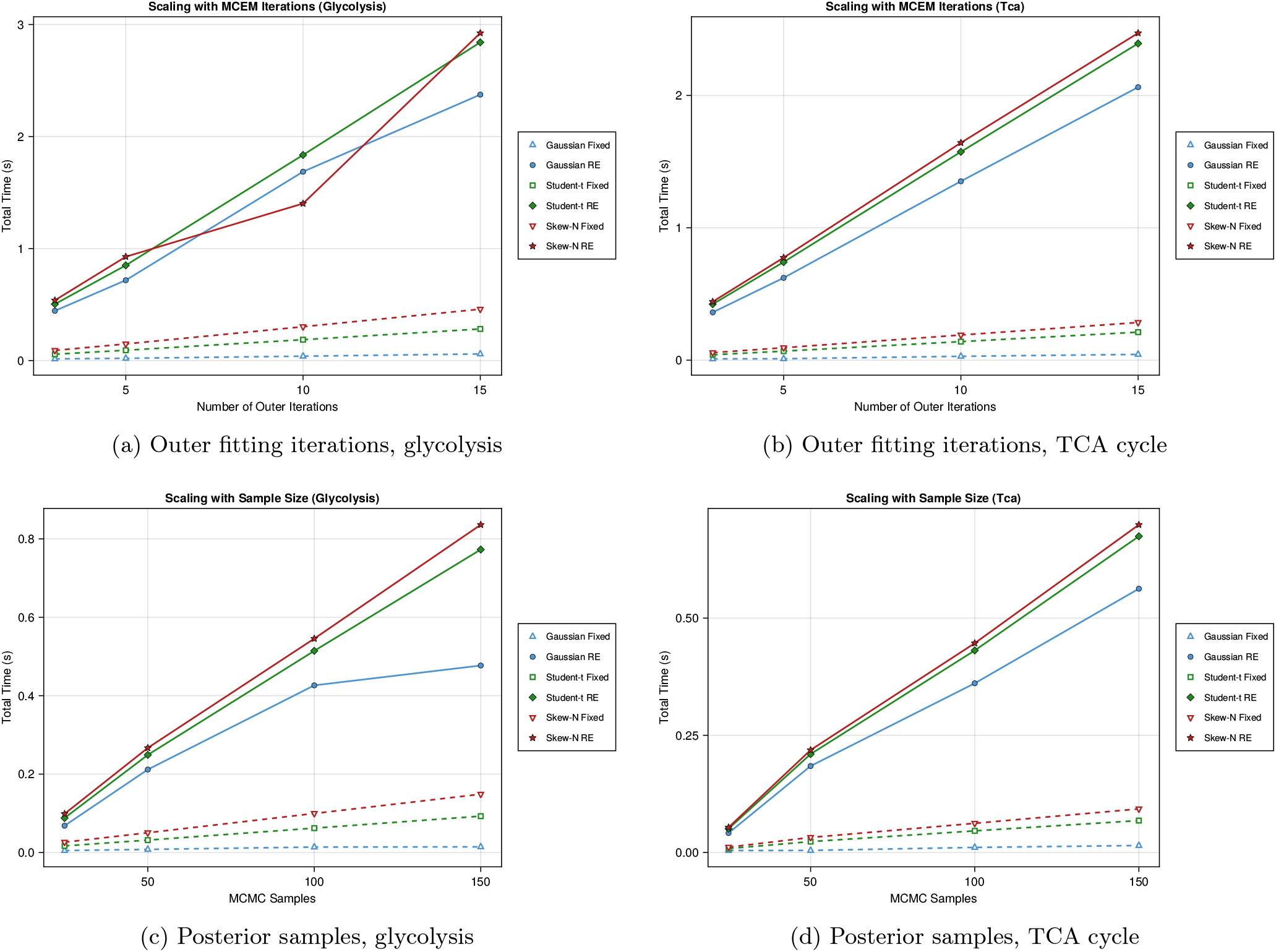
Algorithmic computational scaling for the collapsed MCEM sampler. The pathway panels vary the number of outer fitting iterations and the number of posterior samples while keeping the pathway design fixed. Runtime increases regularly over the tested ranges. FE denotes fixed effects and RE denotes random effects.

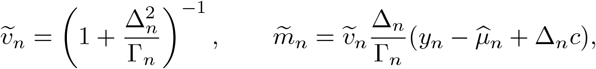

and the update sets the auxiliary value to 𝔼 [*U*_*n*_|*U*_*n*_ *>* 0] for *U*_*n*_~ *N* (*m*_*n*_,*v*_*n*_). Conditional on these deterministic moments, the posterior of ***q*** and the source effects remains Gaussian and is computed using the same sparse Schur-complement factorisation as the collapsed sampler. Thus the approximation reduces the cost of the robust part of the algorithm from repeated inner stochastic refreshes to one deterministic latent-moment pass per outer iteration, plus final Gaussian draws used by the posterior-predictive simulation step.

A small single-threaded benchmark compared the collapsed sampler with the exact-ELBO structured variational option on the same sparse hierarchical synthetic fold (*D* = 24, *N* = 160, *G* = 20, four active basis coefficients per row). The robust shape parameters were held fixed to isolate the cost of posterior inference, and timings were summarized by the median after an excluded warmup run. As shown in Table S4, point-estimation summaries were nearly unchanged, while wall-clock time decreased by a factor of 13.20 for the Student-*t* fit and 18.74 for the skew-normal fit. The structured variational runs produced smaller posterior standard deviations in this diagnostic benchmark, so the collapsed sampler remains the default for the calibrated uncertainty results reported in the main paper. The variational option is intended for computational sensitivity analyses and for scaling studies where the conjugate linear structure should be exploited before moving to more general methods such as Hamiltonian Monte Carlo.

#### F.2.3 Julia package and command-line workflow

The classical parameter-balancing software report [21] emphasizes a practical workflow in which model structure, kinetic data, thermodynamic data, prior ranges, and standardized exchange formats are combined through an online interface, a stand-alone Python command-line tool, a Python package, and a Matlab toolbox. The Julia implementation accompanying this paper follows the same workflow principle while exposing the robust Bayesian extensions used in the present analyses. The repository can be loaded as the Julia package `RobustBayesianParameterBalancing`, and the stand-alone helper `bin/parameter-balance.jl` provides command-line access to the common data-preparation and Gaussian balancing operations.

**Figure S4:**
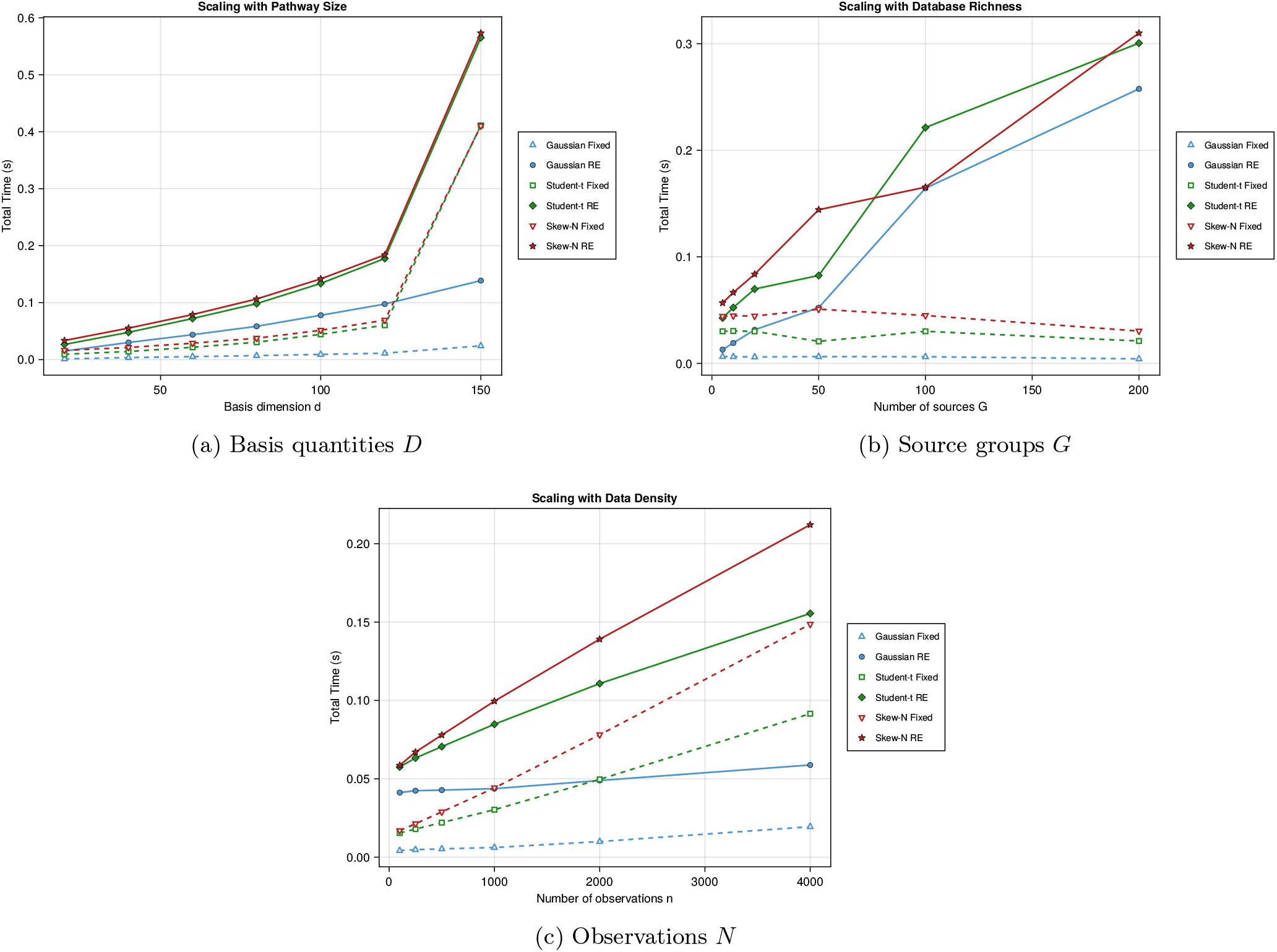
Structural computational scaling for the collapsed MCEM sampler. The panels vary the basis dimension *D*, number of source groups *G*, and number of observations *N*. Fixed-effects models remain cheaper than the corresponding random-effects models because the hierarchical variants update and propagate source-specific effects, but the overhead remains predictable at pathway scale. FE denotes fixed effects and RE denotes random effects.

The package interface separates three tasks that are conceptually distinct in parameter balancing. First, prior and option tables define biochemical plausibility ranges, mathematical quantity types, and pseudo-observation defaults. Second, matrix- and table-level helpers validate data, construct Systems Biology Table (SBtab)-like quantity tables, and export posterior summaries. Third, the inference routines call either the closed-form Gaussian update or the robust fold-wise MCEM and structured-variational routines described above. This organization makes the computational workflow reproducible from scripts while preserving interoperability with SBtab-like tables, comma-separated value and tab-separated value (CSV/TSV) files, and Systems Biology Markup Language (SBML)-oriented downstream tools.

A typical scripted workflow is therefore

~~~
using RobustBayesianParameterBalancing
prior = load_parameter_balancing_prior()
report = validate_pb_inputs(model, quantity_table; prior = prior)
result = balance_parameters((Q = Q, x = x, Cx = Cx,
                            q_prior = q_prior, C_prior = C_prior), prior)
balanced = export_balanced_table(result; format = :dataframe)
~~~

The corresponding file-based Gaussian workflow is

**Figure S5:**
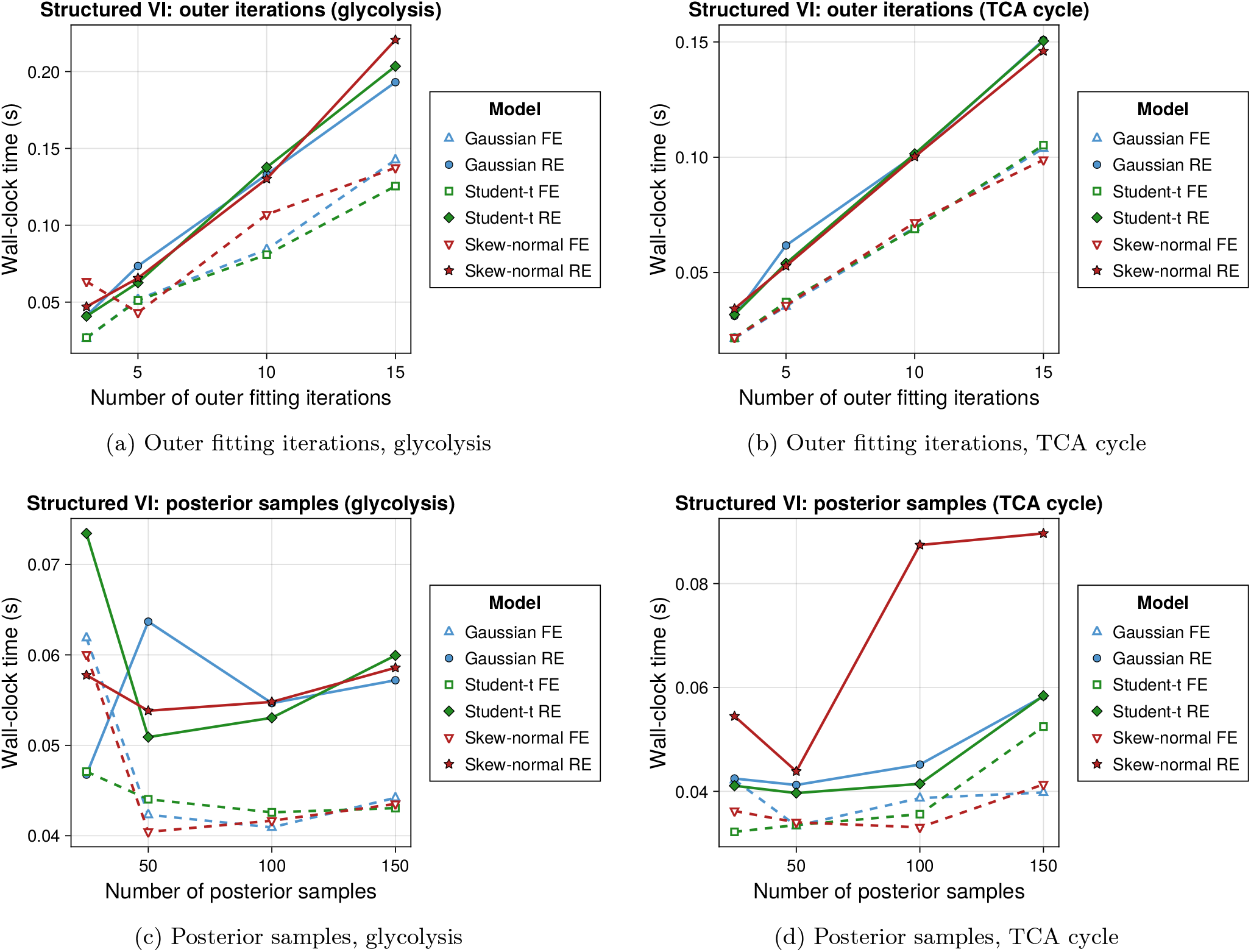
Algorithmic computational scaling for the exact-ELBO structured variational approximation. The pathway panels match Figure S3, with the collapsed sampler replaced by the structured variational approximation described in Section D.9. Runtime again increases regularly with outer fitting iterations and posterior sample count. FE denotes fixed effects and RE denotes random effects.

~~~
julia --project=. bin/parameter-balance.jl balance-gaussian \
   --Q Q.csv --x x.csv --Cx Cx.csv --q-prior q_prior.csv \
   --C-prior C_prior.csv --output balanced_parameters.csv
~~~

This command-line layer is intentionally conservative: it gives modellers a reproducible way to validate data tables, run the conjugate Gaussian baseline, and exchange balanced parameter summaries, while the robust hierarchical analyses remain available through the Julia API and the paper experiment drivers where their fold structure and Monte Carlo settings can be specified explicitly.

### F.3 Additional held-out calibration results on raw BRENDA data

The first group of figures extends the main held-out analyses by examining predictive calibration across nominal levels and by contrasting average scoring rules with empirical coverage. These summaries are useful because models with similar point-predictive accuracy can still differ substantially in uncertainty reliability.

The record-level sensitivity analysis fits the same six model specifications under leave-one-record-out cross-validation. This analysis uses the same posterior-predictive metrics as the leave-one-source-out analysis, including predictive bias, predictive RMSE, CRPS, WIS, log predictive density, PIT values, and multilevel predictive coverage. The distinction between the two holdout schemes is inferentially important. In leave-one-record-out cross-validation, a random-effects model can usually condition on source effects estimated from other records belonging to the same source, so the task measures interpolation to an unseen record. In leave-one-source-out cross-validation, the held-out source is absent from training and the random effect is integrated over as an unobserved source effect; this is therefore the stricter diagnostic for source-level heterogeneity. We accordingly use the source-level analyses for the formal paired tests below, while the record-level analyses provide a complementary check that point-predictive error and interval calibration remain separated at the individual-record scale.

**Figure S6:**
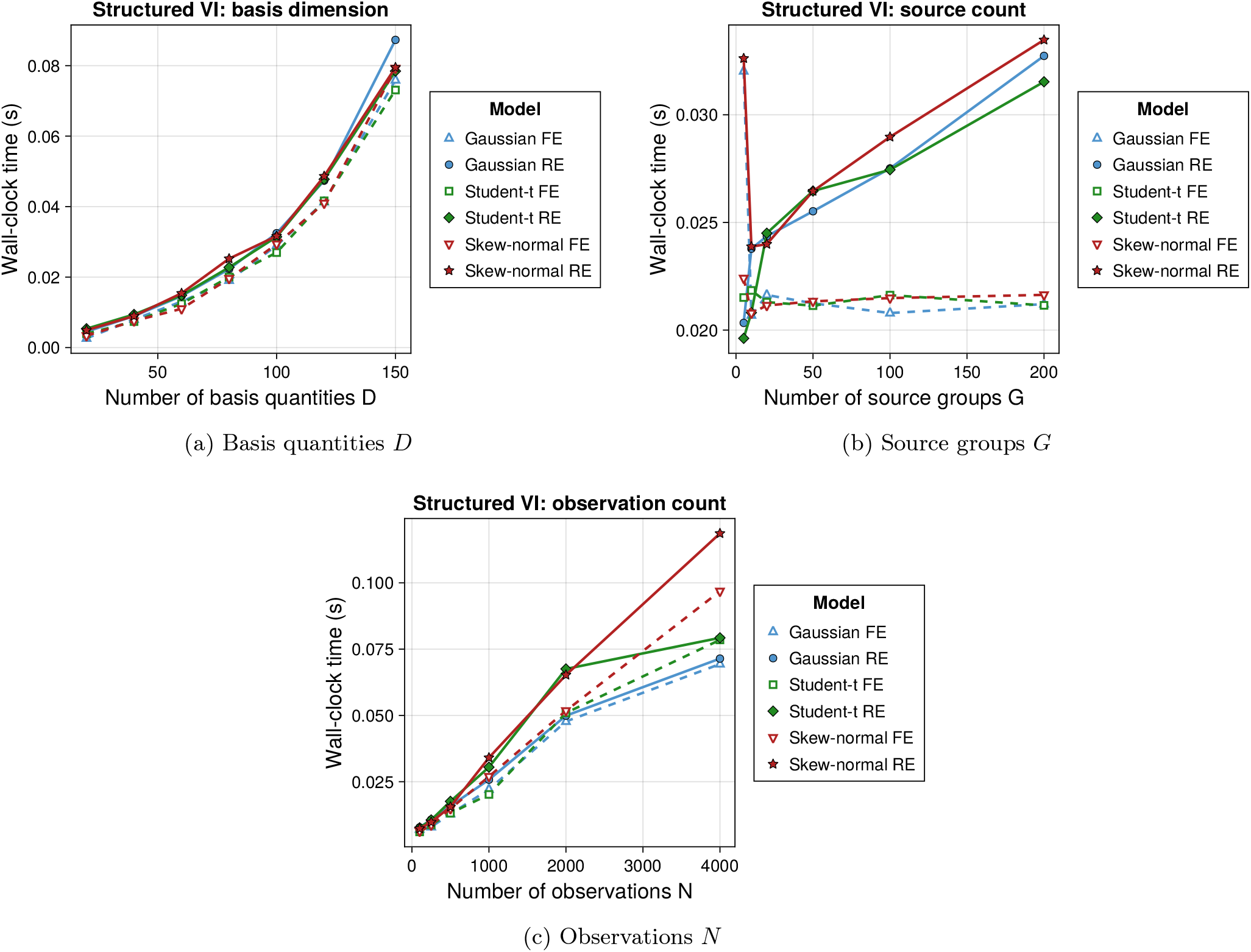
Structural computational scaling for the exact-ELBO structured variational approximation. The structural grids match Figure S4, with the collapsed sampler replaced by the structured variational approximation described in Section D.9. Robust fixed-effects fits remain close to the Gaussian fixed-effects reference because the robust local variables are updated by deterministic exact-ELBO coordinate moments; the dominant additional cost is the source-level random-effects component. FE denotes fixed effects and RE denotes random effects.

The raw-data analysis comprised 155 TCA source folds, 1167 TCA record folds, 230 glycolysis source folds, and 2625 glycolysis record folds. Summary ranges across the three fixed-effects fits and the three random-effects fits are reported in Table S6. In source-level cross-validation, random effects raised empirical 95% predictive coverage from 0.680–0.698 to 0.831–0.848 in the TCA cycle, and from 0.685–0.718 to 0.834–0.839 in glycolysis, with only small changes in predictive RMSE. In record-level cross-validation, the Student-*t* random-effects model gave the lowest CRPS and WIS in both pathways and the highest 95% coverage among the record-level fits, but coverage still remained below nominal (0.728 in the TCA cycle and 0.741 in glycolysis). These record-level results reinforce the main interpretation: improvements in point-predictive scores do not by themselves guarantee calibrated uncertainty intervals.

### F.4 Where hierarchical models increase uncertainty

Improved coverage under random effects can arise either from indiscriminate widening of all intervals or from targeted widening on the most weakly identified quantities. The next figures show that the latter pattern dominates in both pathways.

#### F.4.1 Rule for highlighting wide intervals

In the interval-width heatmaps and combined diagnostic panels, red labels mark parameters whose 95% posterior interval width falls in the top 10% of widths in more than one fitted model. This rule is applied uniformly across pathways and model families, so highlighted labels indicate coordinates that are repeatedly weakly identified rather than quantities that are wide only under a single specification.

**Table S3:** Thread-level timing benchmark for the parallelized experiment loops. Times are wall-clock seconds for a small reproducibility benchmark run with 1, 2, and 4 worker threads. Speedup is relative to the one-thread run for the same workload. The source-level leave-one-source-out (LOSO) benchmark has only 24 independent folds, so scheduler overhead can dominate at two threads; larger record-level and synthetic-replicate workloads show clearer gains.

| Workload | $T_1$ | $T_2$ | $T_4$ | Speedup 2 threads | Speedup 4 threads | Parallel unit |
| --- | --- | --- | --- | --- | --- | --- |
| Record-level LOOCV | 2.454 | 1.466 | 0.878 | 1.67 | 2.80 | 96 folds |
| Source-level LOSO | 0.594 | 0.446 | 0.218 | 1.33 | 2.72 | 24 folds |
| Synthetic recovery sweep | 0.097 | 0.056 | 0.039 | 1.72 | 2.49 | 48 replicates |

**Table S4:** Diagnostic benchmark for the structured variational inference option. Both methods were run on the same sparse hierarchical synthetic fold with BLAS restricted to one thread. Times are medians over five timed repeats after an excluded warmup run. The speedup column is relative to the collapsed sampler for the same likelihood family. SD denotes standard deviation. 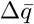 is the mean absolute difference between posterior means from the structured variational option and the collapsed sampler.

| Likelihood | Method | Time (s) | Speedup | Mean absolute error | RMSE | Mean posterior SD | $\Delta\bar{q}$ |
| --- | --- | --- | --- | --- | --- | --- | --- |
| Student-$t$ | Collapsed MCEM | 0.021 | 1.00 | 0.0694 | 0.0909 | 0.0364 | 0.0000 |
| Student-$t$ | Structured VI | 0.002 | 13.20 | 0.0693 | 0.0930 | 0.0318 | 0.0062 |
| Skew-normal | Collapsed MCEM | 0.030 | 1.00 | 0.0601 | 0.0777 | 0.0384 | 0.0000 |
| Skew-normal | Structured VI | 0.002 | 18.74 | 0.0587 | 0.0782 | 0.0293 | 0.0113 |

### F.5 Source-level coverage comparisons

The following figures move from aggregate summaries to source-level comparisons. Fixed-effects and random-effects fits are compared directly for glycolysis and the TCA cycle in Figures S11 and S13, while the three random-effects likelihood families are compared with one another in Figures S12 and S14. Together, these figures show that the gains from hierarchical modelling are not driven by a small number of unusual sources, but instead arise broadly across groups in both pathways.

### F.6 Source-level nonparametric comparison of predictive performance

#### Inferential unit

All inferential comparisons were carried out at the *source* level rather than at the level of individual observations. This choice is essential because leave-one-source-out cross-validation produces multiple held-out observations from the same source, and those observations are not statistically independent once the scientific unit of heterogeneity is the source itself. Accordingly, for each pathway, model, and endpoint, we first aggregated held-out predictive summaries within source and only then applied paired or blocked nonparametric tests across sources.

#### Source-level endpoints

Let 𝒢 denote the set of eligible sources under a given comparison. For each source *g*∈ 𝒢 and fitted model specification *h*, let 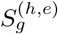 denote the source-level summary for endpoint *e*, where *e*∈ {CRPS, WIS, Cov95}. Here CRPS and WIS are source-level means of the corresponding observation-wise scoring rules, and Cov95 is the source-level mean of the indicator that the held-out observation falls inside the nominal 95% predictive interval.

For CRPS and WIS, smaller values are preferred. For Cov95, larger values are preferred because all fitted models in the present experiments remained below the nominal level 0.95.

#### F.6.1 Paired Wilcoxon signed-rank analysis for random effects versus fixed effects

Fix a pathway, a model family *f* ∈ {Gaussian, *t*, Skew}, and an endpoint *e*. Let 𝒢_*f,e*_ be the set of sources for which both the fixed-effects and random-effects versions of family *f* produced valid source-level summaries for endpoint *e*. For each source *g* ∈ 𝒢_*f,e*_, define the paired difference

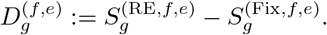

**Table S5:** Package-facing workflow and interoperability functions. The Julia package provides direct application programming interface (API) calls for scripted analyses and a command-line helper for common file-based tasks. The SBML command exports a balanced-parameter sidecar table; complete kinetic-law synthesis is left to downstream SBML tooling.

| Interface | Function or command | Purpose |
| --- | --- | --- |
| Julia API | `load_parameter_balancing_prior` | Load biochemical prior medians, spreads, bounds, and quantity metadata |
| Julia API | `load_parameter_balancing_options` | Load workflow options such as pseudo-observation and parametrisation settings |
| Julia API | `validate_pb_inputs` | Check prior tables, quantity tables, and matrix payload dimensions |
| Julia API | `make_empty_quantity_table` | Build an SBtab-like data-entry template from a reaction list |
| Julia API | `fill_pseudo_observations` | Add prior pseudo-observations for missing quantity slots |
| Julia API | `balance_parameters` | Run closed-form Gaussian balancing or delegate to robust fold inference |
| Julia API | `posterior_observable_summary` | Propagate posterior basis-parameter uncertainty to model quantities |
| Julia API | `export_balanced_table` | Export balanced posterior summaries as DataFrame, CSV, TSV, or SBtab-like text |
| Julia API | `information_gain` | Summarize entropy reduction from prior to posterior |
| Julia API | `check_bounds` | Flag posterior means or intervals outside configured biochemical bounds |
| Julia API | `kineticize_sbml` | Export balanced parameters as an SBML-compatible sidecar table |
| Command line | `prior-summary` | Print or write the normalized prior table |
| Command line | `validate-table` | Validate a quantity table against the prior definitions |
| Command line | `pseudo-fill` | Add pseudo-observation rows using a supplied template |
| Command line | `balance-gaussian` | Run Gaussian balancing from numeric matrix files |
| Command line | `kineticize` | Export a balanced-parameter sidecar for an SBML model |

Here RE denotes the random-effects fit and Fix denotes the fixed-effects fit. Thus:

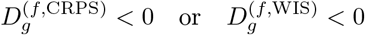

favours the random-effects model, whereas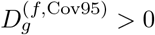favours the random-effects model.

#### Null and alternative hypotheses

For each family *f* and endpoint *e*, we tested

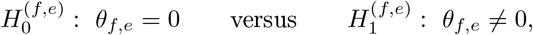

where *θ*_*f,e*_ denotes the location parameter targeted by the signed-rank test; under the usual symmetry formulation this is the center of symmetry of the distribution of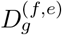,and under the Hodges–Lehmann interpretation it is the pseudomedian of that distribution [64].

#### Signed-rank statistic

Write the nonzero paired differences as 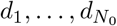,where *N*_0_ is the number of nonzero pairs after removing exact zeros. Rank the absolute values 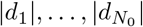from smallest to largest, obtaining ranks 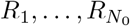. In the presence of ties, average ranks are used. Define

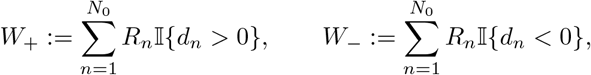

and equivalently the Wilcoxon signed-rank (WSR) statistic *T*_WSR_ := min(*W*_+_, *W*_*−*_). Under the no-tie, no-zero null, the sign pattern is equiprobable conditional on the absolute ranks, so the exact null distribution of *W*_+_ is obtained by all subset sums of {1, …, *N*_*0*_} [65]. In particular,

**Table S6:** Held-out predictive summaries from the raw-BRENDA analyses. Ranges are computed across the Gaussian, Student-*t*, and skew-normal likelihoods within each fixed-effects or random-effects class. Cov95 denotes empirical coverage of the nominal 95% posterior-predictive interval. Lower CRPS and RMSE indicate better point-predictive performance, whereas Cov95 should be interpreted relative to the nominal target 0.95.

| Pathway | Holdout unit and model class | Folds | CRPS range | RMSE range | Cov95 range |
| --- | --- | --- | --- | --- | --- |
| TCA cycle | Source, fixed effects | 155 | 1.772–1.799 | 3.236–3.255 | 0.680–0.698 |
| TCA cycle | Source, random effects | 155 | 1.692–1.710 | 3.226–3.247 | 0.831–0.848 |
| TCA cycle | Record, fixed effects | 1167 | 1.658–1.703 | 3.098–3.105 | 0.692–0.715 |
| TCA cycle | Record, random effects | 1167 | 1.601–1.656 | 2.941–2.955 | 0.692–0.728 |
| Glycolysis | Source, fixed effects | 230 | 1.605–1.649 | 2.969–2.971 | 0.685–0.718 |
| Glycolysis | Source, random effects | 230 | 1.556–1.584 | 2.994–3.006 | 0.834–0.839 |
| Glycolysis | Record, fixed effects | 2625 | 1.550–1.602 | 2.874–2.889 | 0.691–0.725 |
| Glycolysis | Record, random effects | 2625 | 1.496–1.553 | 2.783–2.798 | 0.703–0.741 |

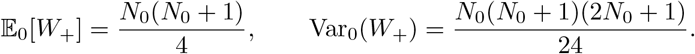

For moderate or large *N*_0_, one uses the standardized statistic

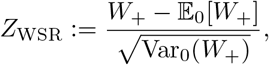

with the usual large-sample normal approximation; when zero differences are present, the normal approximation and its adjustment require care [66, 67]. In the reported computations in Table S7, the endpoint Cov95 is discrete and therefore tie-heavy, so tie-aware asymptotic *p*-values were used. For CRPS and WIS, ties were negligible.

#### Decision rule

At nominal level *α* = 0.05, the two-sided signed-rank test rejects 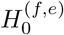 if the exact or asymptotic two-sided *p*-value associated with *T*_WSR_ is at most *α*.

#### Hodges–Lehmann estimator and confidence interval

For each paired comparison we also report the Hodges–Lehmann estimator

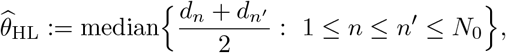

together with a 95% confidence interval obtained by inversion of the signed-rank test [64]. For CRPS and WIS, negative values of 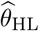 favour random effects; for Cov95, positive values favour random effects.

The exact signed-rank null distribution relies on symmetry of the paired differences 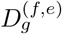 about *θ*_*f,e*_ under *H*_0_ [65]; the Hodges–Lehmann estimator 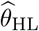 targets the corresponding pseudomedian [64]. Under symmetry this pseudomedian coincides with the population median, but without symmetry it need not do so. Because the paired differences here are source-level contrasts of the same model fitted with and without random effects, approximate symmetry under *H*_0_ is plausible but not guaranteed; we therefore report 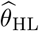 and its signed-rank confidence interval as a rank-based location summary rather than as an exact median estimate.

#### F.6.2 Friedman analysis among random-effects models

We next restricted attention to the three random-effects models: ℳ_RE_ = {Gaussian RE, *t* RE, Skew-N RE}. Fix a pathway and an endpoint *e*. Let the matched source set be 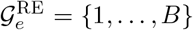, where every source in this set has valid summaries under all three random-effects models. For source *g* and model *k* ∈ {1, 2, 3}, write *Y* for the 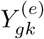source-level endpoint.

##### Within-source ranking

Within each source *g*, the three model values are ranked. For CRPS and WIS, smaller values receive better ranks; for Cov95, larger values receive better ranks. Let 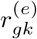 denote the resulting rank, with average ranks used for ties. Define the column rank sums

**Figure S7:**
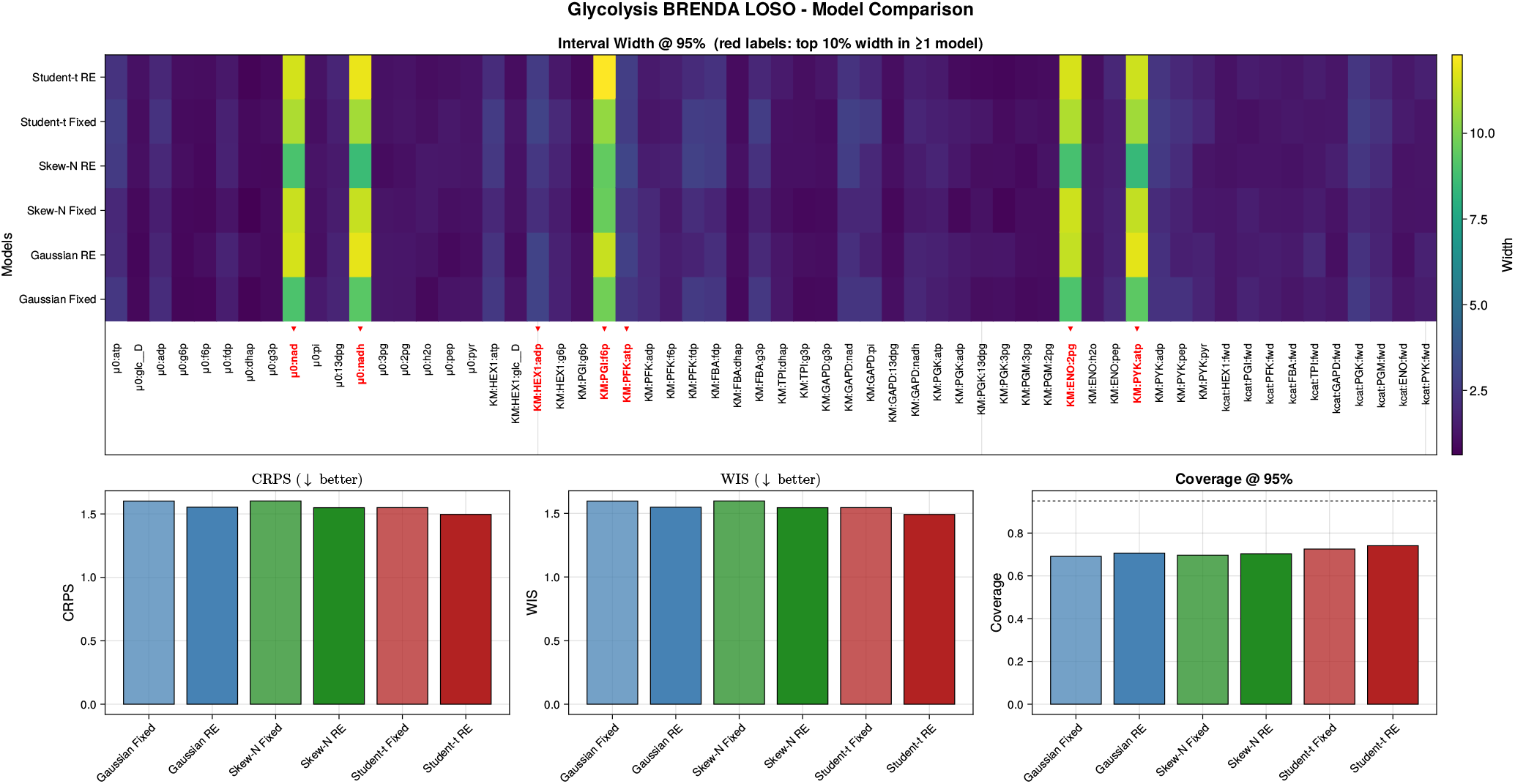
Held-out predictive assessment for glycolysis under leave-one-record-out cross-validation on raw BRENDA data. The panel layout matches the source-level diagnostic in Figure S9. Record-level leave-one-out evaluates interpolation to an unseen record while other records from the same source may remain in the training set. The 2625-fold analysis shows improved point-predictive scores for the Student-*t* random-effects model, but the 95% predictive coverage remains below nominal for all six model specifications.

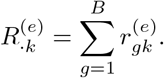

##### Null and alternative hypotheses

For each endpoint *e*, the Friedman null is

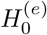 : the three RE models are exchangeable within source with respect to endpoint *e*, against the alternative that at least one model has a systematically different rank distribution across sources [68].

##### Friedman statistic

The classical Friedman statistic is

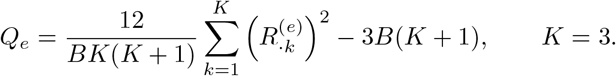

Equivalently, since *K* = 3 here,

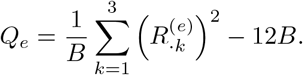

Under 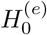 and the usual large-*B* approximation,

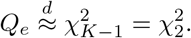

The omnibus null is rejected at level *α* = 0.05 when

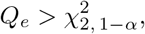

or equivalently when the corresponding *p*-value is at most *α*.

The omnibus Friedman test followed by post-hoc procedures with family-wise error-rate control is a standard protocol in benchmark-comparison studies involving several methods evaluated across multiple datasets or groups [69, 70]; we adopt the same logic here, with sources playing the role of datasets.

**Figure S8:**
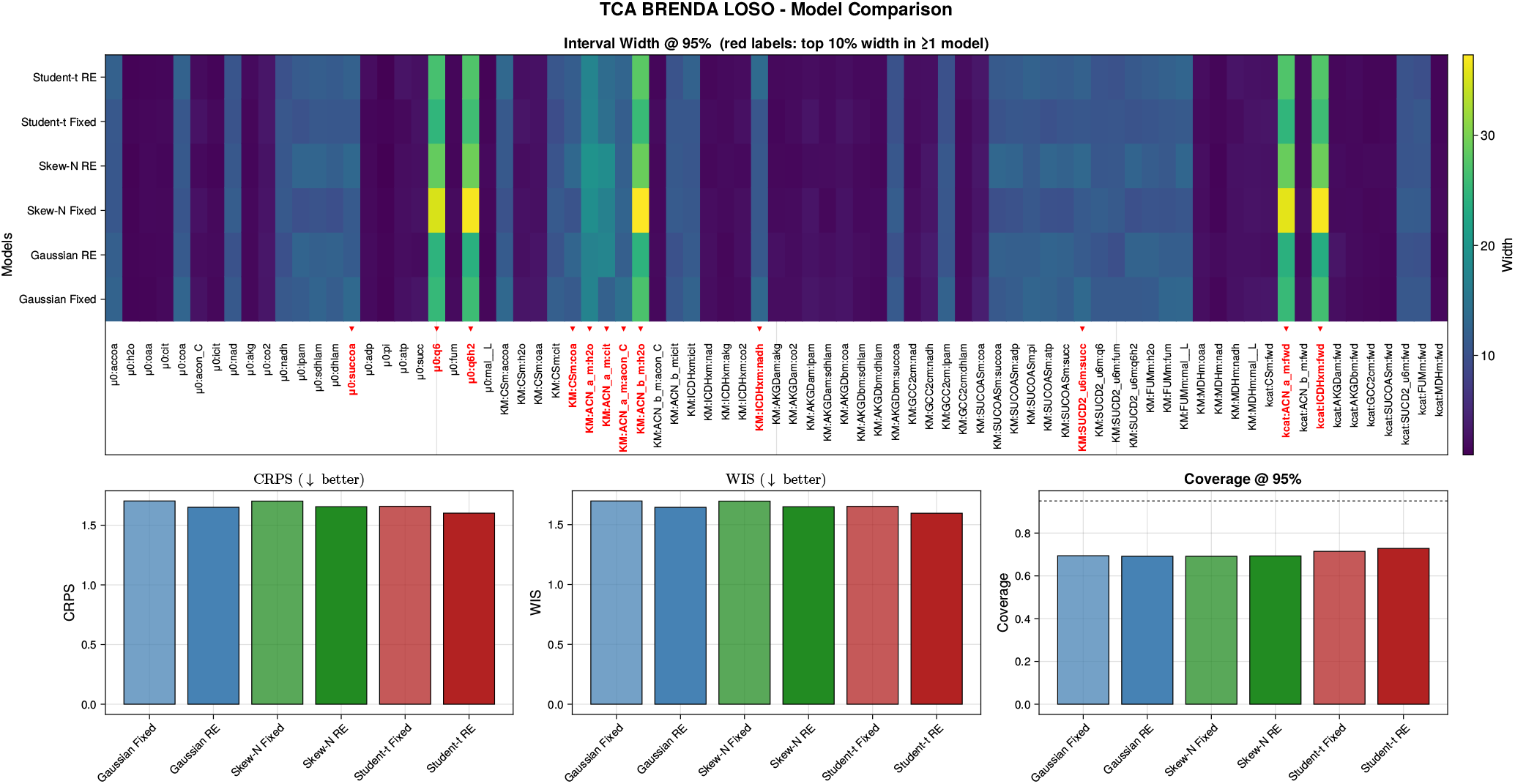
Held-out predictive assessment for the TCA cycle under leave-one-record-out cross-validation on raw BRENDA data. The 1167-fold record-level analysis complements the stricter leave-one-source-out diagnostics. As in glycolysis, record-level point-predictive scores improve for the robust random-effects fits, especially the Student-*t* random-effects model, but 95% coverage remains below the nominal level, demonstrating that point-predictive accuracy and interval calibration remain distinct diagnostics.

#### F.6.3 Holm-corrected pairwise signed-rank follow-up

Whenever the omnibus Friedman test was significant, we carried out the three pairwise source-level comparisons

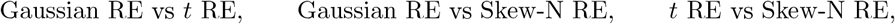

using the same paired signed-rank procedure described above. Let the resulting raw *p*-values be *p*_1_, *p*_2_, *p*_3_. After ordering them as *p*_(1)_ ≤ *p*_(2)_ ≤ *p*_(3)_, Holm’s sequentially rejective procedure rejects the smallest null if 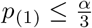, then the second if 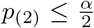, and finally the third if *p* _(3)_ ≤ *α*, stopping at the first non-rejection [71]. Table S8 reports Holm-adjusted pairwise *p*-values.

#### F.6.4 Empirical results

##### Random effects versus fixed effects

The paired source-level signed-rank analyses show that the random-effects formulation yields a statistically significant improvement in empirical 95% coverage for all three observation families in both pathways. In the TCA cycle, these gains are concentrated in calibration: the coverage improvements are highly significant, whereas the source-level CRPS and WIS differences are not statistically distinguishable from zero. In glycolysis, by contrast, random effects improve both calibration and predictive scores: for all three observation families, source-level CRPS and WIS decrease significantly and empirical 95% coverage increases significantly. Thus, source heterogeneity is a first-order contributor to predictive uncertainty in both pathways, but its impact on the full predictive distribution is appreciably stronger for glycolysis than for TCA.

##### Comparison among random-effects models

Within the random-effects class, the omnibus Friedman tests are significant for source-level mean CRPS and mean WIS in both pathways, and the Holm-corrected pairwise follow-up isolates a consistent ranking: the Student-*t* random-effects model achieves significantly smaller source-level CRPS and WIS than the Gaussian and skew-normal random-effects models, while Gaussian and skew-normal random-effects fits are not distinguishable on these two endpoints. For source-level 95% coverage, the omnibus Friedman test is significant in both pathways, but no Holm-corrected pairwise comparison is significant. This pattern indicates that, once source heterogeneity is modeled, the three random-effects families operate in a similar calibration regime at the nominal 95% level, and the remaining practically relevant difference among them is driven mainly by sharpness and score-based performance rather than by a pairwise-resolved coverage advantage.

Taken together, these source-level tests reinforce the main empirical message of the paper. First, source heterogeneity must be modeled explicitly: random effects materially improve predictive calibration, and in glycolysis they also improve the full predictive distribution. Second, after source heterogeneity is accounted for, the choice of observation family becomes a second-order but still meaningful issue: the Student-*t* random-effects specification yields the best source-level CRPS and WIS, whereas the Gaussian and skew-normal random-effects specifications remain competitive in calibration. This pathway-dependent pattern is biologically plausible for curated biochemical databases assembled across heterogeneous laboratories and assay conditions, where between-source variability and occasional heavy-tailed deviations can coexist.

**Figure S9:**
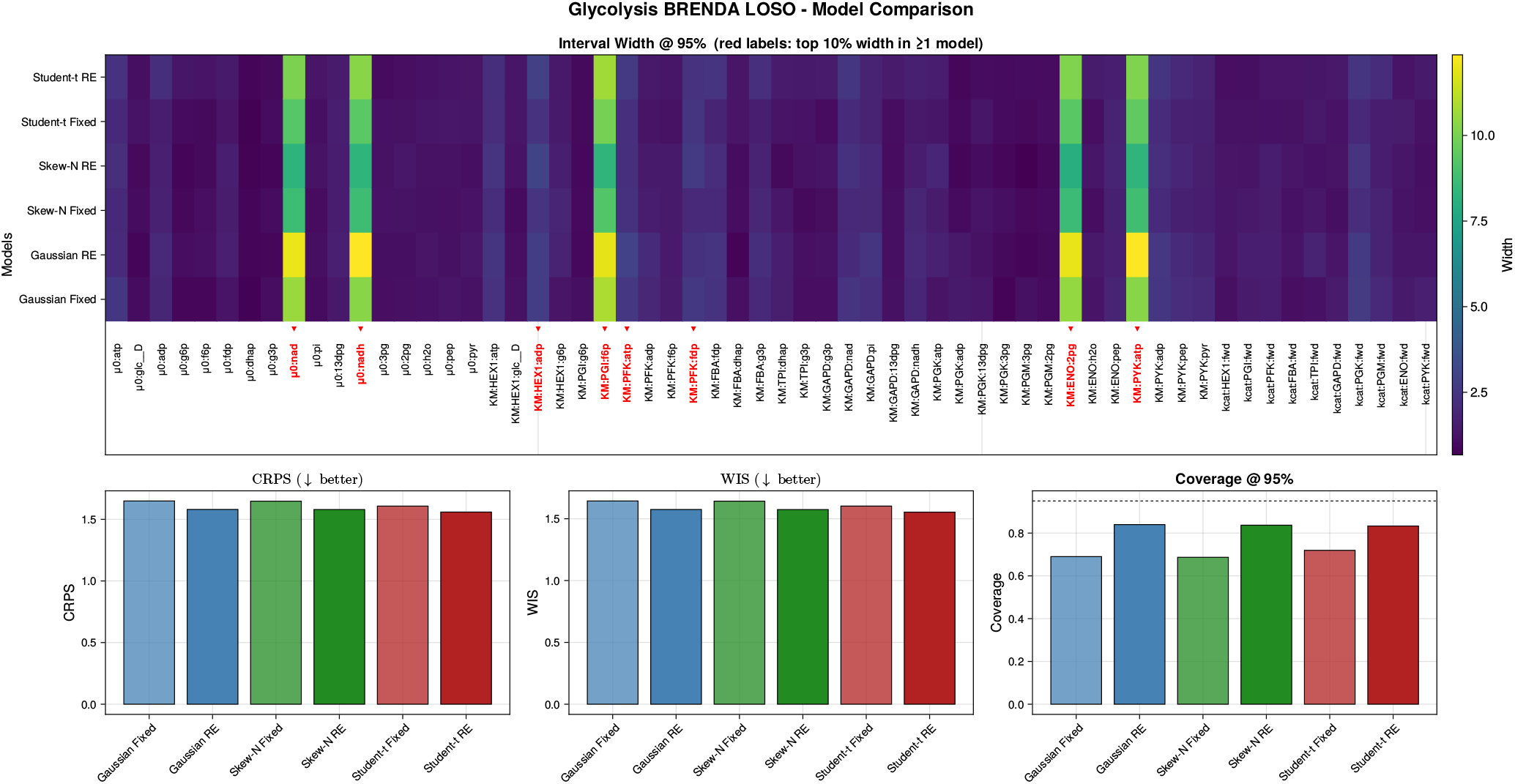
Held-out predictive assessment for glycolysis under leave-one-source-out cross-validation on raw BRENDA data. The upper-left panel shows empirical predictive coverage across nominal levels, with the dashed diagonal representing exact calibration. The upper-right panel reports parameter-specific widths of the 95% posterior intervals, with red labels marking coordinates flagged as unusually wide according to the interval-width rule defined in Supplementary Section F.4.1. The lower panels summarize CRPS, WIS, and empirical coverage at the nominal 95% level. Random-effects formulations improve predictive calibration much more clearly than they improve average scoring rules: CRPS and WIS differ only modestly, whereas fixed-effects models systematically under-cover. The widening induced by the hierarchical fits is concentrated in a small subset of parameters rather than spread uniformly across the pathway, consistent with localized source-specific heterogeneity.

**Table S7:** Paired source-level Wilcoxon signed-rank comparisons of random-effects versus fixed-effects fits. For CRPS and WIS, negative differences favour the random-effects model; for Cov95, positive differences favour the random-effects model. The Hodges–Lehmann estimate is the signed-rank location estimate, and the confidence interval is the associated 95% interval obtained by inversion of the signed-rank test.

| Pathway | Comparison | Endpoint | $n$ | Mean diff. | HL est. | CI low | CI high | $p$ |
| --- | --- | --- | --- | --- | --- | --- | --- | --- |
| TCA | Gaussian RE vs Gaussian Fixed | mean_crps | 112 | −0.0480 | −0.0081 | −0.0315 | 0.0148 | 0.7484 |
| TCA | Gaussian RE vs Gaussian Fixed | mean_wis | 112 | −0.0502 | −0.0090 | −0.0325 | 0.0122 | 0.6770 |
| TCA | Gaussian RE vs Gaussian Fixed | coverage95 | 112 | 0.1386 | 0.1250 | 0.1080 | 0.1250 | **$1.58 \times 10^{-11}$** |
| TCA | Student-$t$ RE vs Student-$t$ Fixed | mean_crps | 112 | −0.0234 | 0.0129 | −0.0078 | 0.0346 | 0.5316 |
| TCA | Student-$t$ RE vs Student-$t$ Fixed | mean_wis | 112 | −0.0248 | 0.0116 | −0.0087 | 0.0323 | 0.5723 |
| TCA | Student-$t$ RE vs Student-$t$ Fixed | coverage95 | 112 | 0.1046 | 0.0833 | 0.0767 | 0.1087 | **$2.39 \times 10^{-10}$** |
| TCA | Skew-N RE vs Skew-N Fixed | mean_crps | 112 | −0.0512 | −0.0044 | −0.0275 | 0.0154 | 0.8197 |
| TCA | Skew-N RE vs Skew-N Fixed | mean_wis | 112 | −0.0539 | −0.0062 | −0.0297 | 0.0126 | 0.7396 |
| TCA | Skew-N RE vs Skew-N Fixed | coverage95 | 112 | 0.1347 | 0.1207 | 0.1000 | 0.1250 | **$2.39 \times 10^{-11}$** |
| Glycolysis | Gaussian RE vs Gaussian Fixed | mean_crps | 125 | −0.0667 | −0.0623 | −0.0727 | −0.0516 | **$8.25 \times 10^{-8}$** |
| Glycolysis | Gaussian RE vs Gaussian Fixed | mean_wis | 125 | −0.0664 | −0.0620 | −0.0721 | −0.0518 | **$5.94 \times 10^{-8}$** |
| Glycolysis | Gaussian RE vs Gaussian Fixed | coverage95 | 125 | 0.1265 | 0.1216 | 0.1000 | 0.1250 | **$1.65 \times 10^{-13}$** |
| Glycolysis | Student-$t$ RE vs Student-$t$ Fixed | mean_crps | 125 | −0.0427 | −0.0338 | −0.0444 | −0.0234 | **0.0021** |
| Glycolysis | Student-$t$ RE vs Student-$t$ Fixed | mean_wis | 125 | −0.0426 | −0.0338 | −0.0446 | −0.0233 | **0.0019** |
| Glycolysis | Student-$t$ RE vs Student-$t$ Fixed | coverage95 | 125 | 0.1097 | 0.0929 | 0.0833 | 0.1000 | **$5.27 \times 10^{-13}$** |
| Glycolysis | Skew-N RE vs Skew-N Fixed | mean_crps | 125 | −0.0650 | −0.0601 | −0.0709 | −0.0499 | **$1.56 \times 10^{-7}$** |
| Glycolysis | Skew-N RE vs Skew-N Fixed | mean_wis | 125 | −0.0650 | −0.0598 | −0.0700 | −0.0498 | **$8.83 \times 10^{-8}$** |
| Glycolysis | Skew-N RE vs Skew-N Fixed | coverage95 | 125 | 0.1379 | 0.1250 | 0.1154 | 0.1339 | **$2.47 \times 10^{-14}$** |

**Table S8:** Source-level Friedman tests among the three random-effects models, followed by Holm-adjusted pairwise signed-rank *p*-values. The omnibus Friedman test compares Gaussian RE, Student-*t* RE, and Skew-N RE within pathway and endpoint.

| Pathway | Endpoint | Friedman $\chi^2$ | Friedman $p$ | G vs $t$ | G vs Skew-N | $t$ vs Skew-N |
| --- | --- | --- | --- | --- | --- | --- |
| TCA | mean_crps | 17.2774 | **$1.77 \times 10^{-4}$** | **0.008210** | 0.484299 | **0.003544** |
| TCA | mean_wis | 17.0710 | **$1.96 \times 10^{-4}$** | **0.011275** | 0.587684 | **0.005524** |
| TCA | coverage95 | 255.4968 | **$3.31 \times 10^{-56}$** | 0.124816 | 0.695312 | 0.103688 |
| Glycolysis | mean_crps | 28.1826 | **$7.59 \times 10^{-7}$** | **$2.75 \times 10^{-6}$** | 0.244782 | **$1.89 \times 10^{-4}$** |
| Glycolysis | mean_wis | 27.0348 | **$1.35 \times 10^{-6}$** | **$4.00 \times 10^{-6}$** | 0.213030 | **$2.93 \times 10^{-4}$** |
| Glycolysis | coverage95 | 349.6261 | **$1.20 \times 10^{-76}$** | 0.901223 | 0.459272 | 0.901223 |

**Figure S10:**
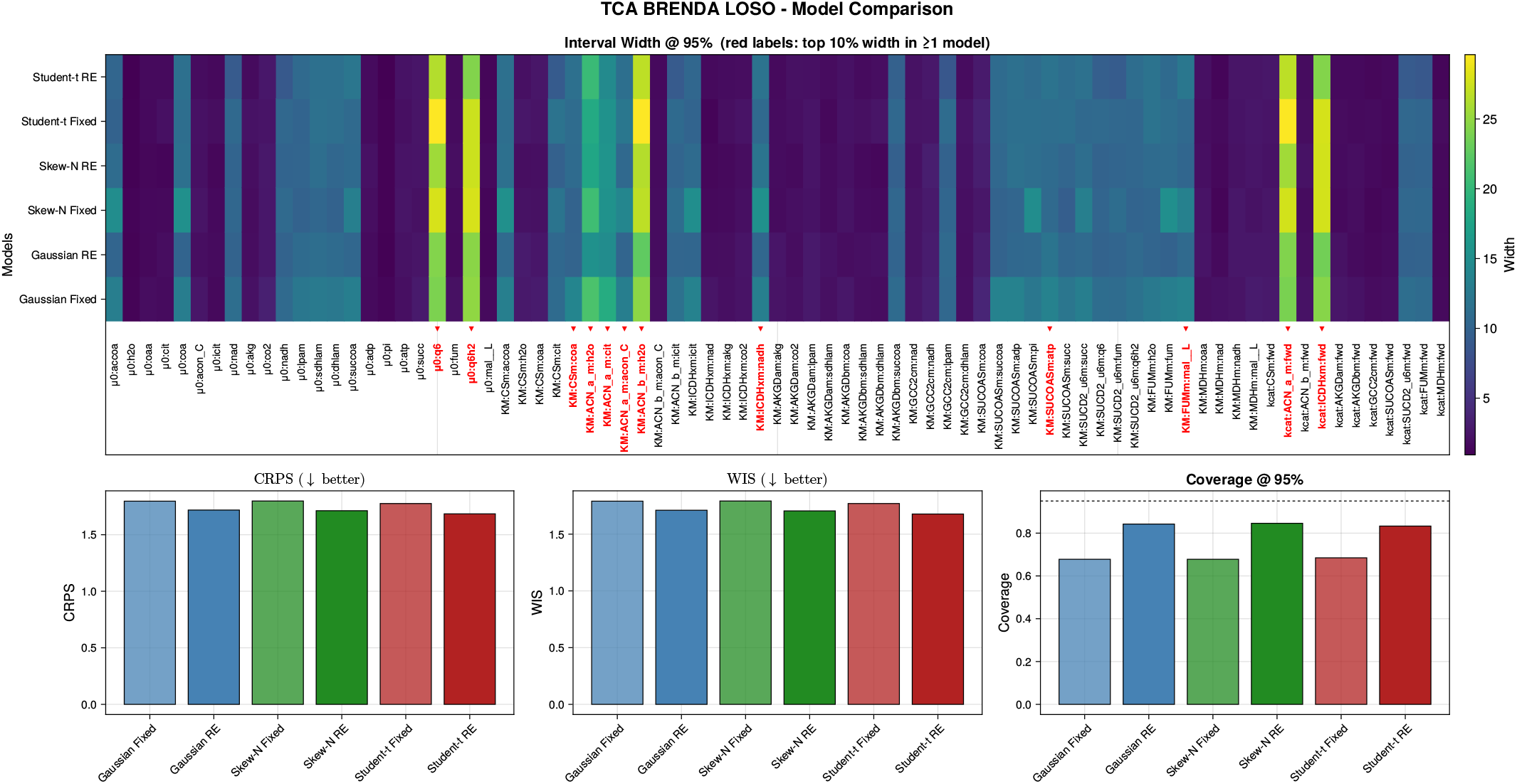
Held-out predictive assessment for the TCA cycle under leave-one-source-out cross-validation on raw BRENDA data. The panel layout mirrors Supplementary Figure S9: multilevel predictive coverage, interval-width diagnostics, CRPS, WIS, and empirical 95% coverage are shown for the competing fixed-effects and random-effects models. Random-effects formulations again improve calibration more clearly than they change average scoring rules. The Student-*t* random-effects model is relatively sharp by CRPS and WIS, whereas the Gaussian and skew-normal random-effects fits are closer to nominal coverage. The interval-width heatmap shows that the added uncertainty is concentrated in a restricted subset of parameters rather than applied uniformly across the pathway.

**Figure S11:**
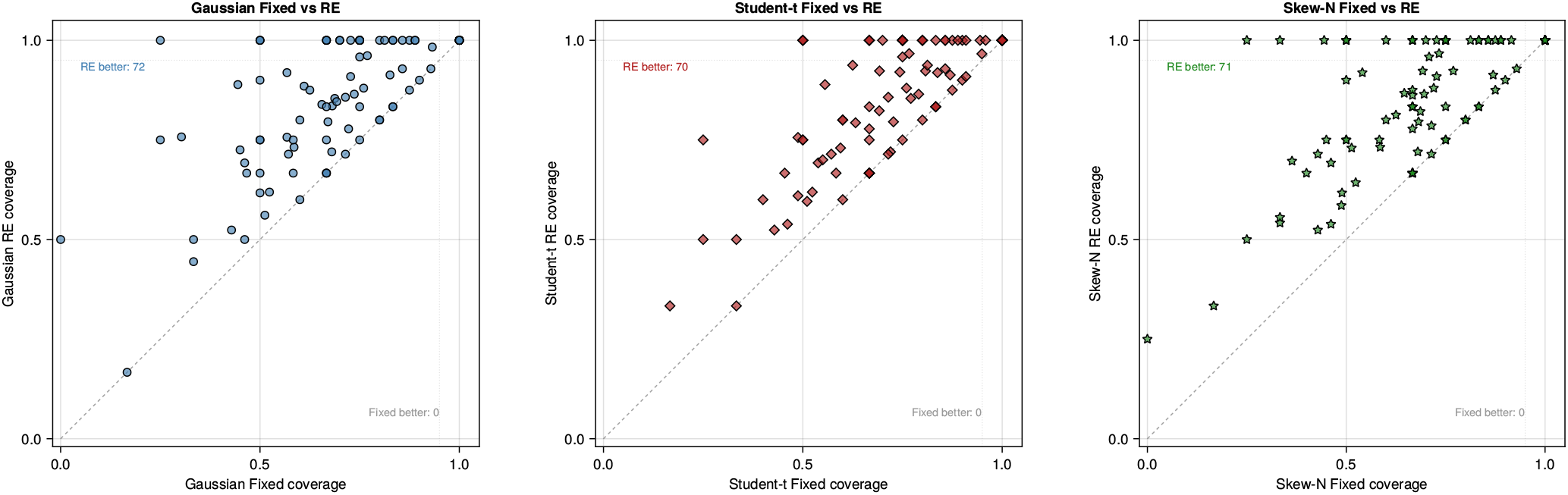
Source-level comparison of empirical 95% predictive coverage for glycolysis, contrasting fixed-effects and random-effects fits within each observation family. Each point corresponds to one source group; the diagonal indicates equal coverage, so points above the line favor the random-effects specification. Across all three noise models, nearly all groups lie on or above the diagonal, and the count annotations summarize how often the hierarchical fit performs better. The improvement is therefore broad-based rather than driven by a few isolated sources.

**Figure S12:**
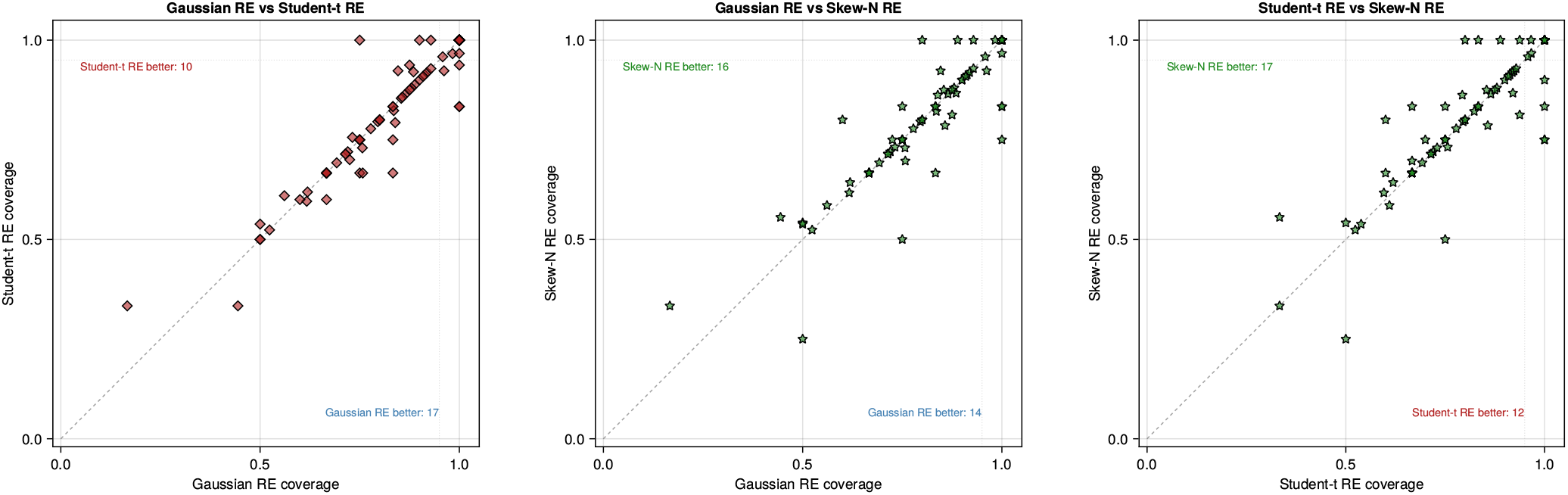
Source-level comparison of empirical 95% predictive coverage among the three random-effects models for glycolysis. Points above the diagonal favor the model on the vertical axis. Most source groups lie close to the diagonal, showing that once source heterogeneity is modeled explicitly, the remaining differences among Gaussian, Student-*t*, and skew-normal random-effects fits are comparatively modest. The skew-normal random-effects model has a slight edge in the count annotations, but the three hierarchical models are much more similar to one another than any of them is to its fixed-effects analogue.

**Figure S13:**
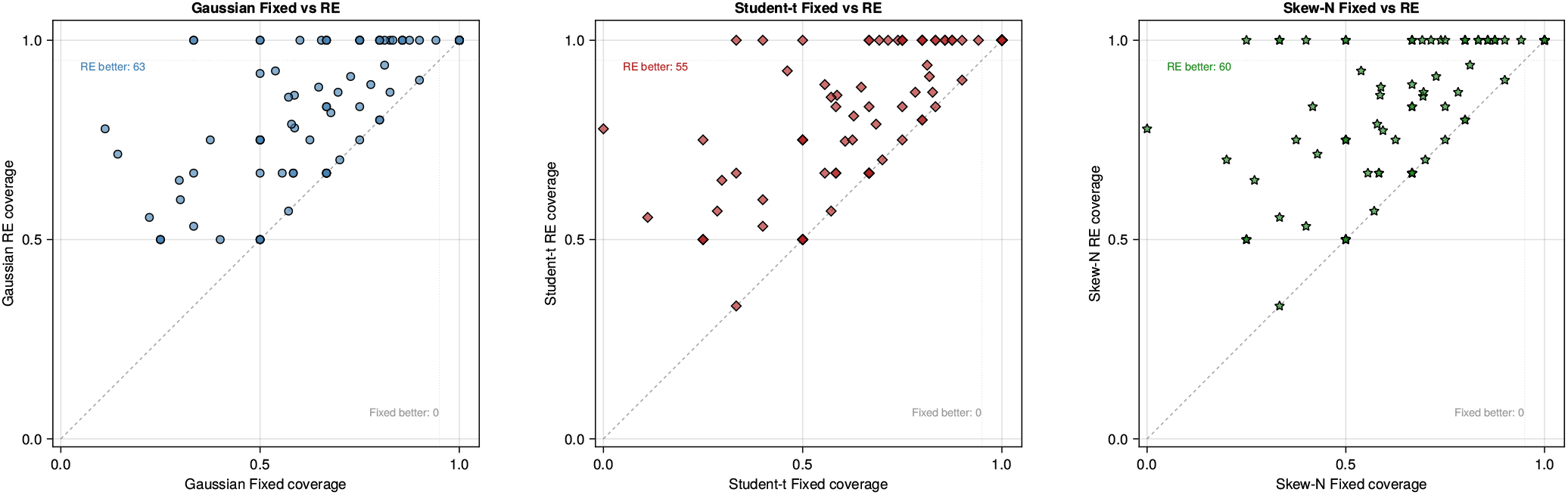
Source-level comparison of empirical 95% predictive coverage for the TCA cycle, again contrasting fixed-effects and random-effects fits within each observation family. The diagonal denotes equal source-wise coverage, and points above the line indicate groups for which the hierarchical fit performs better. For all three observation families, the random-effects specification dominates almost uniformly. The improvement is especially visible for source groups that were markedly under-covered under fixed effects, suggesting that the hierarchical component corrects systematic source-level miscalibration rather than merely inflating uncertainty everywhere.

**Figure S14:**
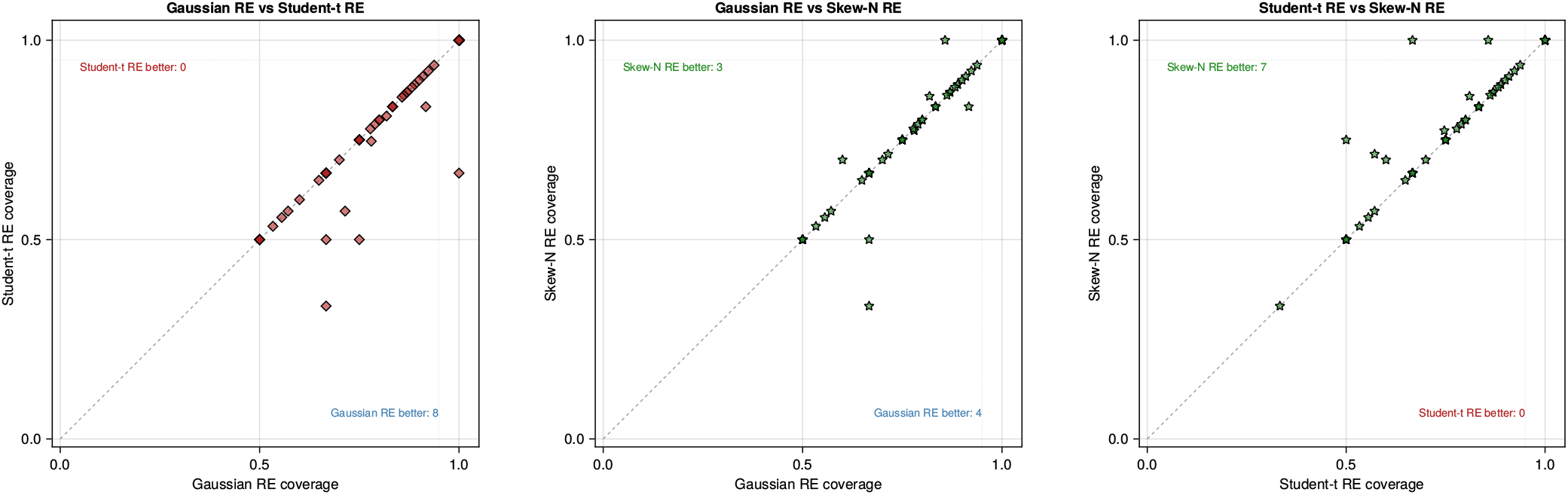
Source-level comparison of empirical 95% predictive coverage among the three random-effects models for the TCA cycle. The Gaussian and skew-normal random-effects fits are very similar overall, with only a small number of groups favoring one over the other, whereas both outperform the Student-*t* random-effects fit on more groups than the reverse comparison. Even so, most points remain close to the diagonal, indicating that the residual differences among hierarchical likelihoods are secondary relative to the much larger gain from introducing source-level random effects in the first place.

